# Concurrent profiling of multiscale 3D genome organization and gene expression in single mammalian cells

**DOI:** 10.1101/2023.07.20.549578

**Authors:** Tianming Zhou, Ruochi Zhang, Deyong Jia, Raymond T. Doty, Adam D. Munday, Daniel Gao, Li Xin, Janis L. Abkowitz, Zhijun Duan, Jian Ma

## Abstract

The organization of mammalian genomes within the nucleus features a complex, multiscale three-dimensional (3D) architecture. The functional significance of these 3D genome features, however, remains largely elusive due to limited single-cell technologies that can concurrently profile genome organization and transcriptional activities. Here, we report GAGE-seq, a highly scalable, robust single-cell co-assay that simultaneously measures 3D genome structure and transcriptome within the same cell. Employing GAGE-seq on mouse brain cortex and human bone marrow CD34+ cells, we comprehensively characterized the intricate relationships between 3D genome and gene expression. We found that these multiscale 3D genome features collectively inform cell type-specific gene expressions, hence contributing to defining cell identity at the single-cell level. Integration of GAGE-seq data with spatial transcriptomic data revealed *in situ* variations of the 3D genome in mouse cortex. Moreover, our observations of lineage commitment in normal human hematopoiesis unveiled notable discordant changes between 3D genome organization and gene expression, underscoring a complex, temporal interplay at the single-cell level that is more nuanced than previously appreciated. Together, GAGE-seq provides a powerful, cost-effective approach for interrogating genome structure and gene expression relationships at the single-cell level across diverse biological contexts.

## INTRODUCTION

Connecting genotype to phenotype remains a challenge due to the complex principles governing genome functions. Mammalian genomes exhibit an intricate organization within the three-dimensional (3D) cell nucleus^1^, encompassing various architectural structures across genomic scales. These structures, including chromosome territories^2^, A/B compartments^3^, subcompartments^3, 4^, topologically associating domains (TADs)^5, 6^ and subTADs^7, 8^, and chromatin loops^9, 10^, play critical roles in gene regulation, cellular development, and disease progression^11–16^. Single-cell analysis can provide unique insights into these processes, uncovering the variability of 3D genome features in individual cells that might be masked in bulk analyses^14, 17, 18^. However, understanding how changes in multiscale 3D genome structure within a single cell influence its transcriptional program and cellular phenotypes remains a major challenge in epigenomics.

Cellular and molecular heterogeneity is a fundamental aspect of cell differentiation and tissue development. Single-cell technologies, such as scRNA-seq and single-cell Hi-C (scHi-C), have advanced our understanding of cellular heterogeneity^19–21^ and 3D genome organization^17, 22–28^. Yet, to fully unravel the connections between 3D genome organization and transcriptional activities in a cell, technologies that can concurrently measure these two molecular properties in the same cell are needed. Current computational methods can provide integrative analysis of scHi-C and scRNA-seq to a certain extent^27, 29, 30^. However, it is currently unattainable to accurately correlate a cell’s 3D genome organization with its gene expression programs using separately generated scHi-C and scRNA-seq data. While existing imaging-based methods do offer this capability, their genomic resolution and throughput are limited^31–34^, demanding the development of new high-throughput genomic technologies capable of co-assaying 3D genome and gene expression in the same cell.

Here, we report GAGE-seq (genome architecture and gene expression by sequencing), a highly scalable and cost-effective method for simultaneously profiling chromatin interactions and gene expression in single cells. GAGE-seq, thanks to its combinatorial barcoding strategy, offers a higher methodological throughput, as well as greater efficiency and effectiveness than recent technologies such as HiRES^35^. We utilized GAGE-seq to profile 9,190 cells across diverse mammalian cell lines and tissue types, including mouse brain and human bone marrow. Specifically, we characterized the intricate connections between the multiscale 3D genome features and cell type-specific gene expression, *in situ* dynamics of both 3D genome and transcriptome in the tissue context, and the temporal interplay between 3D genome rewiring and transcriptional reprogramming during normal human hematopoiesis. This work presents an experimental and analytical framework for examining genome structure and gene expression relationships in single cells across diverse biological systems.

## RESULTS

### Overview of GAGE-seq

GAGE-seq is a high-throughput, effective, and robust single-cell multiomics technology that simultaneously profiles the 3D genome and transcriptome in individual cells (**Fig. 1a**). GAGE-seq leverages the highly scalable “combinatorial indexing” paradigm previously employed in sci-Hi-C^22, 36–38^, as well as other single-cell methods^39–42^ (**Fig. 1a**). The procedure can be summarized as follows: (i) The RNA in cross-linked and permeabilized cells or nuclei is reverse transcribed (RT) with a biotinylated poly(T) or random hexamer primer containing DNA sequences, facilitating the ligation of the first-round barcoded cDNA adaptors (**Fig. S1, Table S1**); (ii) Cross-linked chromatins are efficiently fragmented (the first round chromatin fragmentation) using two 4-cut restriction enzymes (RE), CviQI and MseI, both producing the same adhesive DNA end 5’-TA, enabling the marking of chromatin interactions via proximity ligation; (iii) After a second round of chromatin fragmentation to introduce adhesive DNA ends for ligating the first-round barcoded DNA adaptors (**Fig. S1, Table S1**), cells/nuclei are distributed to a 96-well plate, where the first-round barcodes for DNA or cDNA are introduced through ligation of barcoded adaptors; (iv) Intact cells/nuclei are then pooled, diluted, and redistributed to a second 96-well plate, where the second-round barcodes for DNA or cDNA are introduced through ligation; (v) After reverse-crosslinking to release barcoded nucleic acids, all genomic DNA and cDNA are pooled, and biotinylated cDNA fragments are separated from genomic DNA with streptavidin beads; (vi) Sequencing libraries for scHi-C and scRNA-seq are separately generated and sequenced (**Methods**); and finally, (vii) Matched scHi-C and scRNA-seq profiles are identified according to the well-specific barcoding combinations (**Fig. 1a, Fig. S1, Table S1, Methods**). This combinatorial cellular indexing strategy can be further extended to achieve even larger throughput using additional rounds of ligation-mediated barcoding.

**Figure 1.**
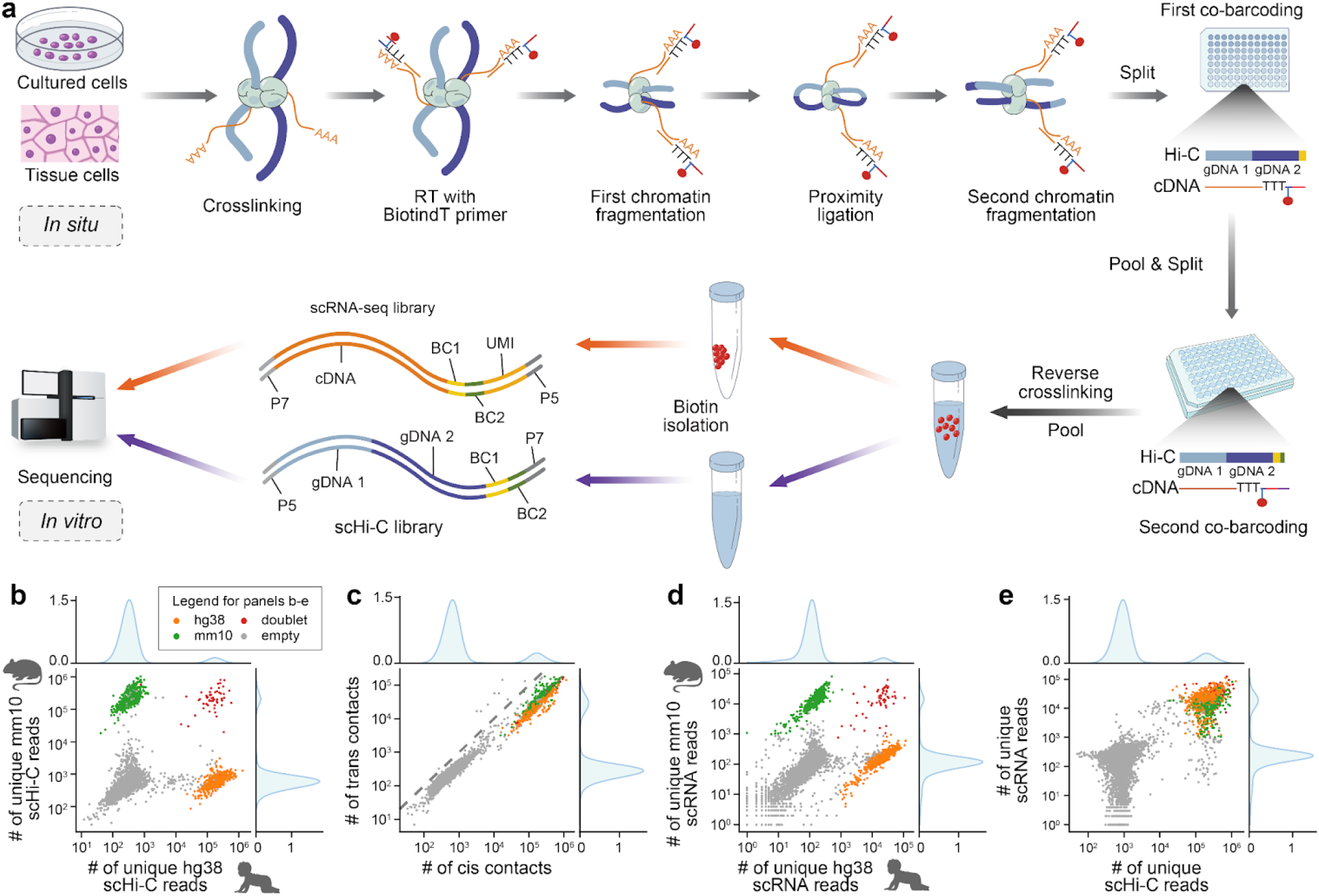
Overview and validation of GAGE-seq. **a**. Schematic representation of the GAGE-seq workflow detailing the simultaneous single-cell profiling of 3D genome architecture and gene expression. **b-e**. Validations demonstrating the specificity of GAGE-seq using mixed experiments with the human (K562) and mouse (NIH3T3). **b** and **d**. Scatter plots showing the collision level in the GAGE-seq scHi-C (**b**) and scRNA-seq (**d**) libraries, and histograms showing the binomial distribution of reads mapped to hg38 (top) and mm10 (right). **c**. Scatter plot showing the cis:trans ratio of scHi-C reads. **e**. Scatter plot showing the well-separation of scHi-C and scRNA reads of valid cellular indices from that of empty indices. Mouse is colored in green, human in orange, collisions in red, and empty indices in gray.

The GAGE-seq strategy can profile tens of thousands of single cells in a few days without the need for physical isolation of the cells and special instruments/reagents, and can be applied to various biological contexts.

### Quality validation and benchmarking of GAGE-seq

To validate the quality and specificity of GAGE-seq data, we first performed experiments using a mixture of human (K562) and mouse (NIH3T3) cell lines (**Fig. 1b-e, Methods, Supplementary Methods**). The successful separation of human and mouse reads in both scHi-C and scRNA-seq data demonstrated the accuracy of GAGE-seq, with 683 human and 568 mouse cells identified out of 1,500 expected, along with 57 doublets observed in line with the expected 4.4% collision rate (**Fig. 1b-e**). Cells passing stringent quality criteria exhibited an average of 181,240 (K562, 39.2% duplicate rate) and 206,113 (NIH3T3, 38.0% duplicate rate) chromatin contacts (>1Kb intra-chromosomal) from scHi-C part, as well as an average of 24,784 (K562, 35.7% duplicate rate) and 16,596 (NIH3T3, 31.2% duplicate rate) unique molecular identifiers (UMIs) obtained from 3,699 (K562) and 2,256 (NIH3T3) genes per cell for the scRNA-seq part (**Fig. 1, Table S2**). These robust results underscore GAGE-seq’s capacity to concurrently measure single-cell chromatin interactions and transcriptome with high sensitivity and accuracy. In addition, GAGE-seq’s efficient fragmentation of crosslinked chromatin before proximity ligation, enabled by two four-cutters (**Fig. 1a, Methods**), allows for efficient detection of multi-way chromatin interactions, with >25% of all identified chromatin contacts in each scHi-C library (**Table S2**).

We expanded the validation of GAGE-seq to additional cell lines, GM12878 and MDS-L, further reinforcing its robustness, specificity, sensitivity, and reproducibility (**Fig. 2, Fig. S2, S3, Methods, Supplementary Methods**). At the whole-genome and whole-library level, we found that the chromatin interaction and gene expression profiles generated by GAGE-seq were strongly correlated with published datasets (**Fig. 2a-b**). The low collision rate (**Fig. 1b**), the binomial distribution of scHi-C reads (**Fig. 1b, Fig. S2a, S3a**), the typical chromatin contact decay curve (**Fig. 2c**), the high *cis*-*trans* ratio (**Fig. 1c, Fig. S2c, S3c, Table S2-S4**), and the aggregated pseudobulk and single-cell chromatin contact maps (**Fig. 2d, Fig. S4, S6**), as well as pseudobulk and single-cell A/B compartment scores and insulation scores (**Fig. 2e**), further confirmed the specificity of the GAGE-seq scHi-C signals. The specificity of the GAGE-seq scRNA-seq signals was demonstrated through the low collision rate (4.6% in the K562/NIH3T3 library) (**Fig. 1d**), the binomial distribution of RNA reads (**Fig. 1d, Fig. S2d, S3d**), and the fact that the majority of RNA reads (86%) mapped to the gene body (**Fig. 2f**), complemented by the pseudobulk and single-cell RNA signal distribution at individual gene loci (**Fig. 2g, Fig. S5**). Notably, similar to SHARE-seq^43^, GAGE-seq scRNA-seq reads were found to be 25%-50% intronic (**Fig. 2f**), indicating enriched nascent RNA. The high reproducibility of GAGE-seq across replicates was demonstrated at multiple levels (**Fig. 2a,b,d,e,g,h,i**), and its methodological resolution (library complexity) of scHi-C matched existing lower-throughput, unimodal methods, such as Dip-C^26, 27^, as well as sn-m3C-seq^44, 45^ (**Fig. 2j**). The scRNA-seq data quality generated by GAGE-seq was also comparable to existing methods (**Fig. 2k**). In line with previous scHi-C studies^23, 36^, GAGE-seq scHi-C data revealed cell cycle stages (**Fig. S6, Supplementary Methods**).

**Figure 2.**
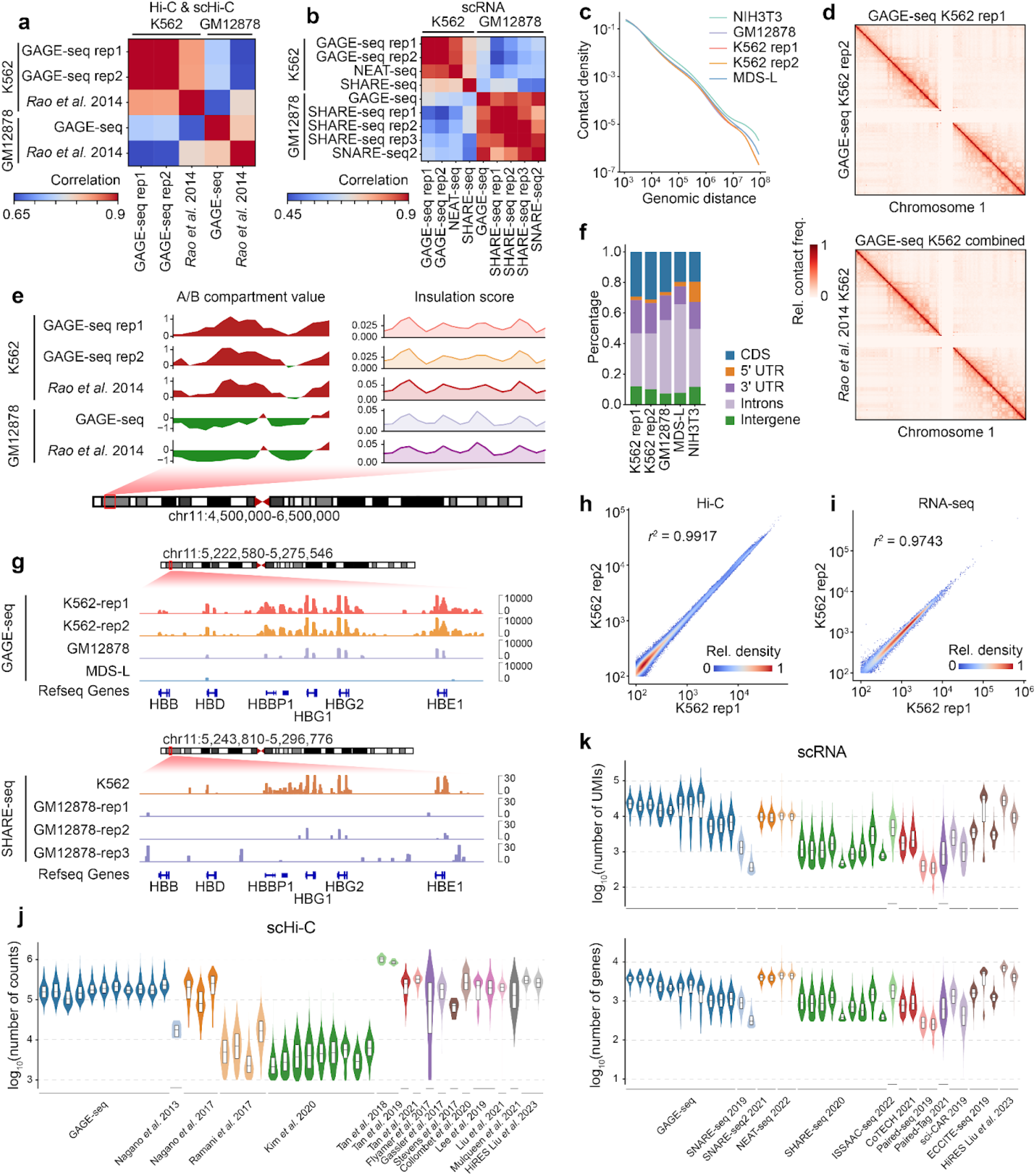
High-quality scHi-C and scRNA-seq data generated by GAGE-seq. **a**. Correlation between the aggregated scHi-C profiles from GAGE-seq replicates and the bulk *in situ* Hi-C data^3^. **b**. Comparison of aggregated scRNA-seq profiles of GAGE-seq replicates with NEAT-seq^68^, SHARE-seq^43^, and SNARE-seq2^67^. **c**. Decay curves of chromatin contact for the GAGE-seq scHi-C libraries. **d**. Comparison of aggregated contact maps between two GAGE-seq K562 replicates (upper), and between the combined GAGE-seq K562 library and an *in situ* Hi-C library^3^ (lower). **e**. Comparison of A/B compartments and TAD-like domain calling at the human beta-globin locus between GAGE-seq (pseudo bulk) and *in situ* Hi-C^3^. **f**. RNA read distribution across gene bodies in the GAGE-seq scRNA libraries. **g**. Aggregated single-cell gene expression profiles at the GAPDH locus. Upper panel: scRNA-seq signals of GAGE-seq libraries of K562, GM12878, and MDS-L cells (hg38). Lower panel: scRNA-seq signals of SHARE-seq in GM12878 cells (hg19)^43^. **h.** Reproducibility between two biological replicates of GAGE-seq scHi-C libraries. **i**. Reproducibility between two biological replicates of GAGE-seq scRNA libraries. **j**. Comparison of GAGE-seq scHi-C library size with published scHi-C^17, 22–27, 37, 62–65^ and co-assay methods^35, 44, 45^. **k**. Comparison of scRNA-seq library size (upper) and the number of detected genes (lower) with published co-assay methods^35, 43, 66–74^.

Compared to the recent HiRES method^35^, GAGE-seq offers several major advantages (**Fig. 2j-k, Fig. S10, S11**). By using a combinatorial barcoding strategy, GAGE-seq achieves a higher methodological throughput, which is an order of magnitude higher than HiRES. In addition, GAGE-seq is more efficient as HiRES relies on quasilinear pre-amplification, known to lead to low mappability of sequence reads^46^. Moreover, GAGE-seq is more cost-effective, as HiRES requires higher sequencing costs per cell and substantial costs from enzymes required in the single-cell quasilinear pre-amplification, PCR amplification and the Tn5-enzyme mediated tagmentation in multi-well plates.

These comprehensive validation experiments and benchmarking collectively affirm the high-quality data produced by GAGE-seq, attesting to its reliability in jointly profiling single-cell chromatin organization and transcriptome.

### GAGE-seq reveals complex cell types in mouse cortex

To demonstrate the utility of GAGE-seq in unveiling complex cell types based on single-cell 3D genome features and gene expression within a tissue context, we turned our focus to the adult mouse brain cortex, known for its varied cell types. Applying GAGE-seq on cells from the mouse cortex (8-9 weeks old), we generated 3,296 high-quality joint single-cell profiles of chromatin interactions and transcriptomes (**Methods, Supplementary Methods**). On average, we observed 231,136 chromatin contacts per cell (at ∼50% duplication rate), and 20,160 UMIs and 1,883 genes per cell (∼59% duplication rate), in line with the adult mouse whole brain data from the recently published HiRES method (**Fig. S7-S9, Fig. 2j-k, Table S5**).

Our GAGE-seq scRNA-seq data successfully identified 28 known cell types for three major lineages in the mouse cortex, including 15 excitatory neuron subtypes, 8 inhibitory neuron subtypes, and 5 glia cell subtypes, such as astrocytes and oligodendrocytes (**Fig. 3a-b, Fig. S12-S14**). These cell identities were confirmed by the unique expression patterns of marker genes (**Fig. 3b**). Notably, our GAGE-seq scRNA-seq data enabled the delineation of many rare neuronal subtypes not identified by HiRES^35^, such as L5 PT CTX, Sncg, and Meis2 (**Fig. 3a-b, Fig. S13-S14**). Although previous studies have suggested that 3D genome features encode cell identity information^35, 47^, scHi-C data often identified fewer cell types in complex tissues than scRNA-seq did^27, 44, 45, 48^. By using our recent method Fast-Higashi^49^ for scHi-C embedding, GAGE-seq distinguished all 28 transcriptome-defined cell types, including the aforementioned L5 PT CTX, Sncg, and Meis2 rare subtypes (**Fig. 3c, Fig. S13-S15**). The scHi-C-based delineation supports these cell types with distinct 3D genome features, with insulation scores surrounding gene bodies showing cell type-specific connection with gene expression (**Fig. 3d**; see later section with more analysis).

**Figure 3.**
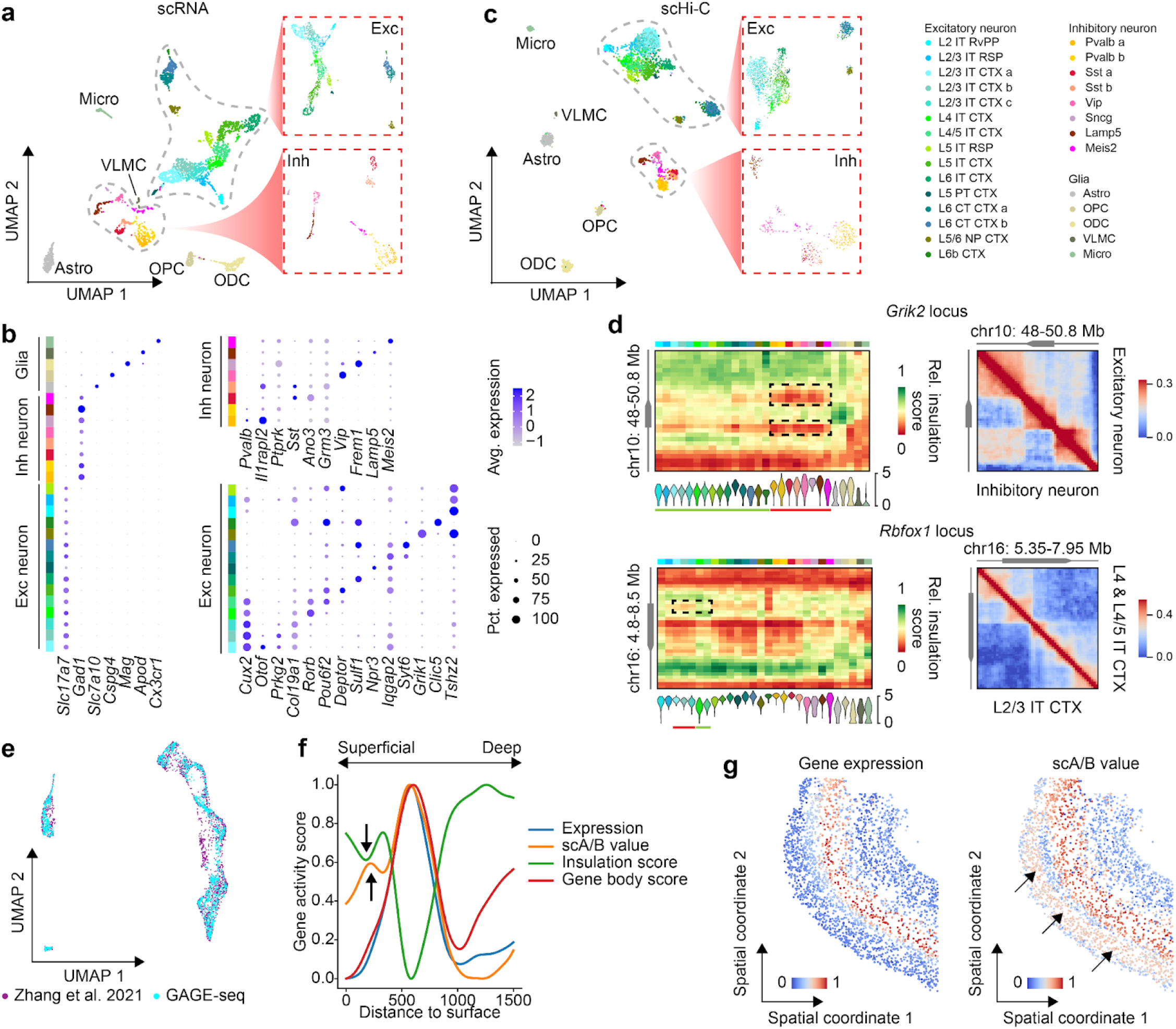
Cell types in mouse cortex characterized by GAGE-seq scHi-C and scRNA-seq. **a** and **c**. UMAP visualization of mouse cortex scRNA-seq (a) and scHi-C profiles (c) from GAGE-seq. Insets: UMAP visualization of excitatory neuron subtypes (top) and inhibitory neuron subtypes (bottom). **b**. Cell type-specific expression (based on scRNA-seq in GAGE-seq) of known marker genes, including glial types, neuronal types, and neuron subtypes. **d**. Visualization of cell type-specific 3D chromatin architecture and gene expression at representative gene loci. Left: aggregated single-cell insulation score (100-Kb resolution, upper) and gene expression (lower) at the *Girk2* locus and the *Rbfox1* locus. Right: aggregated contact maps (50-Kb resolution) of the *Girk2* locus (top panel, excitatory vs inhibitory neurons) and the *Rbfox1* locus (low panel, L4 & L4/5 IT CTX vs L2/3 CTX). Cell types selected in the right panels are highlighted by green lines (higher expression) or red lines (lower expression) in the corresponding left panels. **e.** UMAP visualization of the integration of GAGE-seq and a MERFISH dataset^50^. **f**. Inferred spatial patterns of gene expression and 3D genome features of L5 IT CTX marker genes. **g**. *In situ* plots of inferred single-cell gene expression (left) and scA/B value (right) for L5 IT CTX marker genes. Layer 3 was highlighted by black arrows in panels (f) and (g).

These results established that both the scRNA-seq and scHi-C data from GAGE-seq can resolve refined cell types, such as neuron subtypes in mouse cortex, facilitating the analysis of cell type-specific connections and variability between 3D genome features and gene expression. The high congruence between cell types defined by scHi-C and scRNA-seq from the GAGE-seq suggest a strong correlation between the two modalities in mouse cortex.

### Spatial integration of GAGE-seq data reveals *in situ* 3D genome variation in tissues

Using GAGE-seq to map the 3D genome and transcriptome of single cells, we sought to examine the *in situ* variation of the 3D genome in the adult mouse cortex. We leveraged GAGE-seq scRNA-seq as a “bridge” for this analysis. Recently, the spatial transcriptomics method MERFISH successfully discerned the spatial organization of distinct cell populations in the mouse primary motor cortex^50^. We started by integrating our GAGE-seq scRNA-seq data with the MERFISH data using Seurat^51^, allowing us to establish a connection between the two datasets (**Methods**).

We focused on the excitatory neuron cell types present in both GAGE-seq and MERFISH datasets. Within the integrated embedding space, cells primarily clustered by cell type, and cells from both datasets integrated cohesively, indicating high correlation between cell types identified by the two methods (**Fig. 3e, Fig. S16, Methods**). We next characterized the *in situ* variation of both marker gene expression and 3D genome features of these maker gene loci in the mouse cortex. As a proof of principle, we investigated the *in situ* pattern of marker genes for L5 intratelencephalic (IT) CTX. The observed and inferred gene expression demonstrated a high degree of congruence, further supporting the reliability of the integration (Spearman’s *r*=0.76, two-sided *P*=0; **Fig. S17b-c, j-k**). Layer 5, where L5 IT CTX cells reside, corresponded with the highest expression level, scA/B value^27^, gene body score (**Supplementary Methods**), and a low single-cell insulation score (**Fig. 3f-g, Fig. S17**), reinforcing the overall correlation between expression and 3D genome structure.

Interestingly, despite consistently low expression levels and gene body scores in more superficial layers, the scA/B value increased and the single-cell insulation score decreased slightly around layer 3, a cortical layer containing the L2/3 IT CTX cells that are not adjacent to the tissue boundary, suggesting potential discrepancies of expression and various 3D genome features at finer spatial resolution (highlighted by arrows in **Fig. 3f-g, Fig. S17g-o**).

Together, such joint profiling of scRNA-seq and scHi-C by GAGE-seq allows for the superimposition of 3D genome features on *in situ* tissues. The spatial patterns of expression and multi-scale 3D genome structures generally align, although they do show discrepancies at finer resolutions.

### Impact of multiscale 3D genome features on gene expressions in single cells

With high-resolution, paired GAGE-seq scRNA-seq and scHi-C data, we next rigorously examined the relationship between gene expression and various multiscale 3D genome features, including A/B compartments, TAD-like domains, and chromatin loops.

Our analysis of the 3,461 genes expressed in inhibitory neurons (n=508) or excitatory neurons (n=1,938) revealed a strong correlation between cell type-specific gene expression and scA/B value, a quantitative measure of compartmentalization variations^27, 29^ (**Fig 4a**, top panels). Inhibitory neurons, for instance, showed a much higher expression for 432 genes which corresponded to a higher scA/B value (t-test *P*=1.1e-46; **Fig. 4a**, top middle panel).

**Figure 4.**
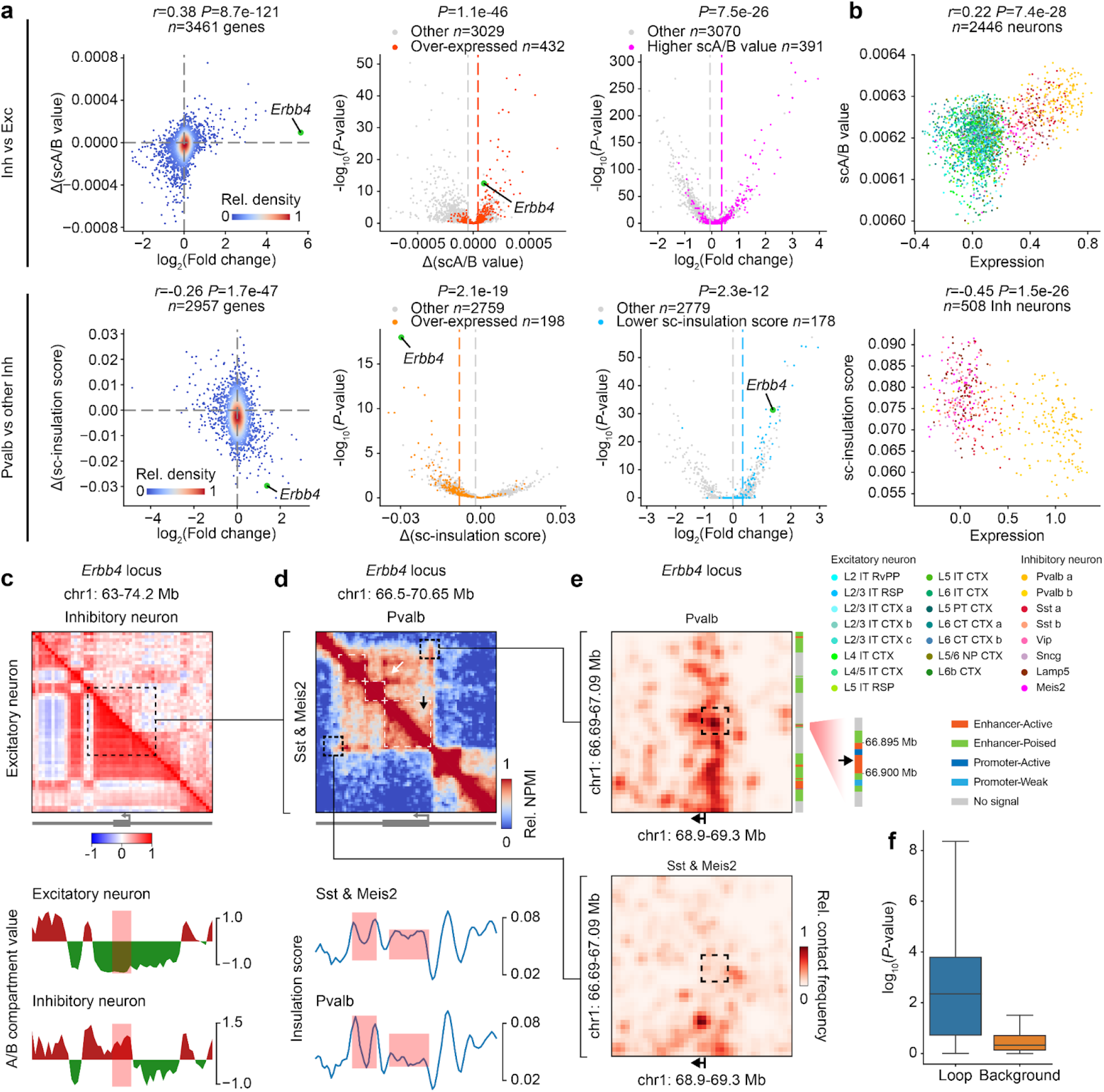
Multiscale 3D genome features inform cell type-specific gene expressions in the mouse cortex at single-cell resolution. **a**. Genome-wide correlations between gene expression and 3D genome features in different neuron cell types. Upper row: correlation for inhibitory (n=508) vs. excitatory neurons (n=1938). Lower row: correlation for Pvalb (n=188) vs. other inhibitory neurons (n=320). Left column: correlation between differential expression and differential 3D genome feature. Middle column: volcano plot of differential scA/B value and single-cell insulation score; Right column: volcano plot of differential expression. Left column: Pearson’s correlation coefficients and the *P*-values from one-sided tests for nonzero correlations are shown. Middle and right columns: *P*-values from one-sided t-tests with unequal variance are shown. **b**. Correlation at the single-cell level between gene expression and scA/B value (upper) or single-cell insulation score (lower) for genes overexpressed in inhibitory neurons (432 genes) and Pvalb cells (198 genes), respectively. Spearman’s correlation coefficients and the *P*-values from one-sided tests for nonzero correlations are shown. **c**. Comparison of A/B compartment (identified at 200-Kb resolution) state of the *Erbb4* locus between inhibitory and excitatory neurons. Correlation matrices of aggregated scHi-C contact maps (top) and the corresponding A/B compartment scoretracks (bottom) are shown. **d**. Comparison of the pseudo-bulk contact map (at 50-Kb resolution) of the *Erbb4* locus between Pvalb and the other inhibitory subtypes. Pseudo-bulk scHi-C contact maps (upper) and the corresponding insulation scores (bottom) are displayed. Two Pvalb-specific bright strides are highlighted by a white arrow and the melted TAD by a black arrow in the top panel. The gene body is shown right under the matrices in the top panels of (c) and (d). Regions with differential 3D genome features are highlighted with light red boxes in the bottom panels of (c) and (d). **e**. An example loop in Pvalb (upper) and Sst and Meis2 (lower) inhibitory subtypes at 10-Kb resolution. Aggregated convolution-smoothed contact maps and regulatory element annotations^72^ in Pvalb are shown. The black maplet arrows show the TSS of Erbb4. **f**. The comparison of loop contacts (blue) and non-loop contacts (orange) regarding their correlation with expression, showing more significant correlation for loop contacts. *P*-values from two-sided tests for nonzero Spearman’s correlation coefficients are shown.

Most of the 391 genes with much higher scA/B value in inhibitory neurons were also expressed at a significantly higher level in these cells than in excitatory neurons (t-test *P*=7.5e-26, **Fig. 4a**, top right panel). Overall, there is a high correlation between differential gene expression and differential scA/B value (Pearson’s *r*=0.38, *P*<1e-100, **Fig. 4a**, top left, **Fig. S18**). Additionally, at the chromatin domain level, we identified a negative correlation between cell type-specific gene expression and the associated single-cell insulation score across cell types (**Fig. 4a**, bottom panels, **Fig. S19**), indicating that TAD-like domain variation surrounding the gene body is accompanied with changes in transcriptional activity of the gene. This phenomenon was also observed at cell type level previously^29^ and may be related to domain melting, which was noted earlier in highly expressed long genes in mouse hippocampus and midbrain neurons^47^.

We therefore examined the relationship between single-cell insulation score surrounding the gene body and the potential occurrence of domain melting within our diverse collection of cell types revealed by GAGE-seq (**Supplementary Methods**). We focused on the four genes (*Grik2*, *Dscam*, *Rbfox1*, and *Nrxn*) known to undergo domain melting^47^, profiling their scA/B value, single-cell insulation score, and single-cell gene expression. Notably, these genes displayed high expression across nearly all of the 28 cell subtypes revealed by GAGE-seq, with the exception of *Dscam* and *Grik2* in VLMC and Micro cells (**Fig. S20, Fig. 3d**). As expected, the scA/B value profiles indicated that *Dscam*, *Rbfox1*, and *Nrxn3* are in the active A compartment in the majority of cell subtypes (**Fig. 3d, Fig. S20**). The *Grik2* locus was in a weak B compartment across all the cells, despite its high expression (**Fig. S20**). Aggregated single-cell insulation scores varied across the gene body, with lower insulation scores in most cell subtypes often correlating with higher gene expression (**Fig. 3d, Fig. S20**). The aggregated chromatin contact maps indicate potential occurrence of domain melting around these gene bodies (**Fig. 3d, Fig. S21**). A similar phenomenon was also detected for the *Rbfox1* locus across different excitatory neurons (**Fig. 3d**, low panels). These observations indicated co-occurrence of potential domain melting, low single-cell insulation score surrounding the gene body, and high gene expression.

We next further confirmed the above observed connection between multiscale 3D genome features and gene expression at a single cell resolution. Higher gene expression in a cell compared to others frequently corresponded to a higher scA/B value and lower single-cell insulation score in the same cell (**Fig. 4b, Fig. S22**). For instance, of the 432 genes showing a significantly elevated scA/B value in inhibitory neurons, most displayed a higher expression level in these neurons than in excitatory neurons (Spearman’s *r*=0.22, *P*=7.4e-28, n=2446 cells; **Fig. 4b**, top panel). At the chromatin domain level, the 198 genes expressed highly in Pvalb cells exhibited notably lower single-cell insulation scores than in the other inhibitory neurons (Spearman’s *r*=0.45, *P*=1.5e-26, n=508 cells; **Fig. 4b**, low panel). Therefore, the connection between multiscale 3D genome features and gene expression can be recapitulated at the single-cell resolution.

We then confirmed our observations on single loci. As a proof of principle, we focused on the Pvalb inhibitory subtype (including both Pvalb a and Pvalb b). We first selected genes that have 1) significantly higher scA/B values and expression in inhibitory neurons compared to excitatory neurons (**Fig. 4a**, top panels, **Fig. S23**), and 2) significantly higher expression and lower single-cell insulation scores in Pvalb compared to other inhibitory neurons (**Fig. 4a**, bottom panels, **Fig. S24**, **Supplementary Method**). This approach led us to the *Erbb4* gene. The *Erbb4* gene plays a pivotal role in the central nervous system and has been linked to schizophrenia^52^. As expected, we observed differential A/B compartment states correlated with cell type-specific expression of the *Erbb4* gene (**Fig. 4c)**, and differential single-cell insulation score that manifests domain melting in the gene locus (**Fig. 4d**, low panel). The TAD-like domain structure of the *Erbb4* gene body in Sst and Meis2 cells appears to be melted in Pvalb cells (i.e., less pronounced), which is again accompanied with high gene expression in Pvalb cells (**Fig. 4d**, top panel). In addition, it appears that the *Erbb4* gene body interacts more frequently with the downstream two small TAD-like domains in Pvalb cells than in Sst and Meis2 cells (**Fig. 4d**, top panel). On the finer scale, we also observed a cell type-specific putative enhancer-promoter chromatin loop at the TSS of the *Erbb4* gene in Pvalb cells (**Fig. 4e-f, Supplementary Methods**).

Together, GAGE-seq enabled us to uncover the intrinsic link between multiscale 3D genome features and cell type-specific gene expression at the single cell level.

### Developmental stages of human hematopoiesis characterized by GAGE-seq

Human definitive hematopoiesis, a developmental process that produces all types of blood cells, originates from CD34+ stem cells in the bone marrow (BM), offering an ideal model system to explore the dynamic relationship between 3D genome structure and gene expression. We generated GAGE-seq profiles of 2,815 human BM CD34+ cells after stringent quality filtering (**Fig. S25-S27, Table S6, Methods, Supplementary Methods**), obtaining an average of 265,336 chromatin contacts (at ∼50% duplication rate) and detecting on average 5,504 UMIs and 985 genes per cell (at ∼63% duplication rate), which is in line with the publicly available scRNA-seq datasets (**Fig. S25-S27, Table S6**). BM CD34+ cells, part of hematopoietic stem/progenitor cells (HSPCs), exhibit considerable molecular heterogeneity. The continuous nature of hematopoiesis^53^ makes distinguishing various types of HSPCs in the CD34+ population based solely on transcriptomic data a challenge. Our goal here is not to exhaustively identify all cell types in the CD34+ population but to elucidate the dynamic relationship between genome structure and function. To mitigate the potential impact of 3D genome’s cell-cycle dynamics^23^, we restricted our analysis to high-quality G0/G1 phase cells (837 cells).

Unsupervised clustering of our GAGE-seq scRNA-seq data yielded six clusters (five clusters with continuous diffusion and one distinct cluster), each displaying unique gene signatures (**Fig. 5a-b**). Based on the gene expression signatures and known marker genes^54^, we annotated these clusters into known cell types, namely hematopoietic stem cell (HSC), multipotent progenitor (MPP), lymphoid-primed MPP (LMPP), multi-lymphoid progenitor (MLP), megakaryocyte-erythroid progenitor (MEP), and B lymphocyte natural killer cell progenitors (B-NK) (**Fig. 5a-b**). These clusters represent all three major blood cell lineages but show a preference towards the lymphoid lineage. As anticipated, progenitor subpopulations at earlier differentiation stages, HSC, MPP, LMPP, MLP and MEP, were clustered more closely, whereas the more lineage-committed B-NK cells separated from the rest (**Fig. 5a**). Our GAGE-seq scHi-C data also successfully resolved these six cell types (**Fig. 5a-b**), further demonstrating the ability of the 3D genome to encode cell type information.

**Figure 5.**
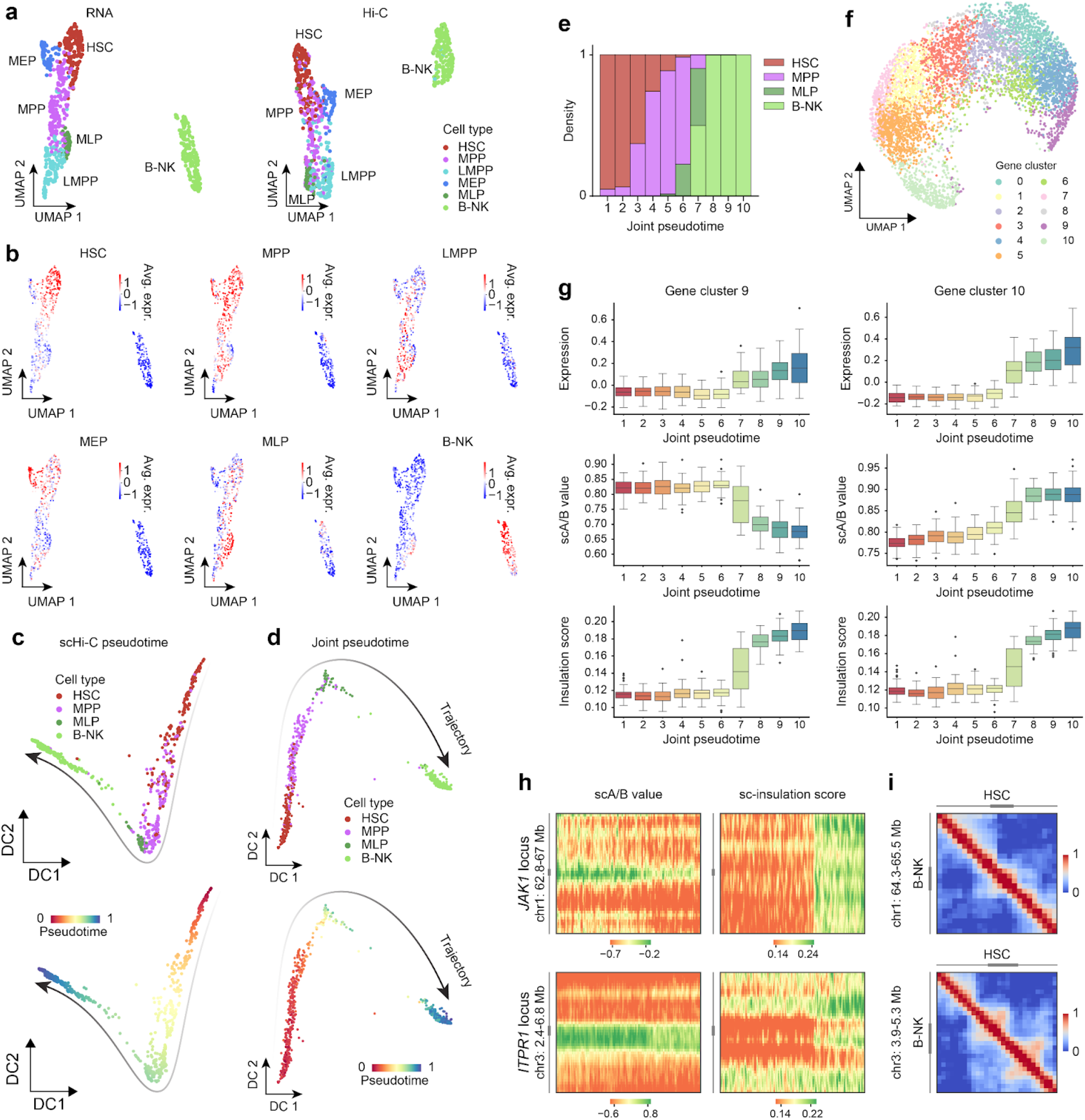
Interplay between 3D genome variation and gene expression changes in human bone marrow differentiation. **a**. UMAP visualization of GAGE-seq scRNA-seq (left) and scHi-C profiles (right) of human bone marrow CD34+ cells. **b**. Average expression of known marker genes on the UMAP plot. The 6 panels include n=124, 78, 24, 82, 126, and 356 genes for HSC, MPP, LMPP, MEP, MLP, and B-NK, respectively. **c**-**d**. Inferred B-NK lineage trajectory and pseudotime from scHi-C profiles (c) and jointly from scRNA-seq and scHi-C profiles (d), displayed by cell type (upper) and pseudotime (lower). **e**. Cell type compositions across 10 equally divided pseudotime bins. **f**. UMAP visualization of gene clusters determined by the temporal trend of expression and scA/B value. **g**. Temporal trends of gene expression (upper row), scA/B value (middle row), and single-cell insulation score (lower row) of gene clusters 9 (left column) and 10 (right column). **h**. scA/B (left) and single-cell insulation score (right) of the *JAK1* (upper) and *ITPR1* (lower) loci (at 100-Kb resolution). Each row represents a cell, ordered by the joint pseudotime from left to right. Heat maps were smoothed by a Gaussian kernel with a receptive field of 10 neighboring cells and 1 neighboring bin in each direction. **i**. Pseudo-bulk contact maps (at 50-Kb resolution) of HSC and B-NK at the *JAK1* (upper) and *ITPR1* (lower) loci.

Four of the six identified cell types (HSC, MPP, MLP and B-NK) represent the early differentiation stages of B-NK cell lineage. We used our GAGE-seq dataset to reconstruct the developmental trajectory of this lineage, demonstrating the dynamic interplay between genome structure and gene expression along this trajectory. Transcriptome and 3D genome-based pseudotime trajectories, inferred from GAGE-seq data, were highly congruent (**Fig. 5c, Fig. S28-S29, Methods**), suggesting that global 3D genome temporal variations overall mirror transcriptional changes and differentiation progression. Further, we created an integrated pseudotime trajectory (**Fig. 5d, Methods**), which was confirmed by the accurate alignment of the four cell types along the differentiation pseudotime and the observation that earlier-stage progenitors (e.g., HSCs) decrease while later-stage cells (e.g., B-NK) increase along the pseudotime (**Fig. 5d-e**), reaffirming that lineage differentiation is a continuous process^53^.

These results underscore the ability of GAGE-seq’s joint profiling of scRNA-seq and scHi-C at delineating major differentiation stages, and inferring the differentiation trajectory and pseudotime in human BM CD34+ cells. The concordance between unimodal pseudotime and the joint pseudotime suggests the general temporal trend of concurrent changes of gene expression and 3D chromatin structures.

### Temporal interplays between 3D genome and gene expression during lineage differentiation

Examining 3D genome dynamics and temporal gene regulation during differentiation pseudotime, we found that while scA/B values temporally shifted across most marker genes, all remained in the A compartment (**Fig. S30-S32**). The scA/B value in the MPP marker genes exhibited little temporal variation, implying regulatory mechanisms beyond A/B compartmentalization (**Fig. S30**). In the HSC marker gene set, scA/B value changes lagged behind gene expression, whereas the MLP and B-NK2 gene sets transitioned to a more active chromatin state before transcriptional activation, denoting a reverse pattern (**Fig. S30**).

When analyzing the aggregated single-cell insulation scores of these marker genes along the differentiation pseudotime (**Fig. S30**), unlike mouse cortex cells, we found no reverse correlation to gene expression. An abrupt insulation score increase during MLP to B-NK lineage commitment signaled global 3D genome rearrangement, corroborated by a surge in mid-range intra-chromosomal contacts and decreased short- and long-range contacts in B-NK progenitors (**Fig. S33**). TAD-like domain rearrangements at marker gene loci in the B-NK cells further supported these findings (**Fig. S31, S32**). Despite the increased portion of chromatin contacts in B-NK cells residing within the mitotic contact range^55^, the chromatin contact maps in these gene loci exhibited clear A/B compartment and TAD-like domain organization (**Fig. S31-S32, Fig. S35-S36**), possibly tied to the quiescent state (G0 phase) of the bone marrow B-NK immature cells^56^.

We then performed an unsupervised clustering to further unravel relationships between gene expression and 3D genome features in the B-NK differentiation trajectory, based on all genes expressed in at least twenty single cells in the trajectory (**Methods**). We identified 11 distinct gene clusters (**Fig. 5f**). Intriguingly, aside from the aforementioned scenarios in the marker gene sets (**Fig. 5g** right panel**, Fig. S34**), we found that 5 of these 11 clusters displayed a negative correlation between the changes in gene expression and scA/B value as the pseudotime progressed (**Fig. 5g** left panel**, Fig. S34**). We closely examined gene cluster 9, where expression is elevated while the scA/B value decreases along the pseudotime. We selected two genes, *JAK1* and *ITPR1*, which exhibit the highest similarity with the average temporal patterns of this gene cluster. The scA/B value at the gene bodies of *JAK1* and *ITPR1* indeed decreases over pseudotime, but we did not observe any A/B compartmentalization switch (**Fig. 5h** left panels), indicating that scA/B value changes may reflect the gradual change within the A compartment of the local chromatin environment of the gene loci. Therefore, we have systematically identified a diverse set of gene groups with distinct temporal patterns, including those with discordant patterns in expression and scA/B value, as reported previously^27^, during differentiation trajectory.

In relation to the chromatin domains, a uniform temporal trend was observable in the aggregated single-cell insulation scores across all gene clusters, mirroring the pattern seen in the marker gene sets (**Fig. 5g, Fig. S34**). This trend suggests global 3D genome changes, manifested by widespread TAD-like domain re-organizations, in B-NK cells. Specifically, for genes *JAK1* and *ITPR1,* the single-cell insulation score increases abruptly from MLP to B-NK, showing a positive correlation with gene expression (**Fig. 5h** right panels). This was confirmed by aggregated contact maps (**Fig. 5i, Fig. S35-S36**). In addition, genes with different sizes appear to behave differently (**Fig. S37**). For instance, among the under-expressed genes in B-NK cells (n=576 genes), shorter genes (< 200kb, n=285) showed a more prominent increase in single-cell insulation score during the MLP to B-NK transition compared to longer genes (>=200kb, n=191) did (one-sided t-test *P*-value=2e-11, **Fig. S37**).

Together, our findings unveiled a landscape of diverse interplays between gene expression and chromatin compartmentalization during the B-NK lineage differentiation. The observed global changes in chromatin interactions within the B-NK cells induced widespread gene locus-associated domain re-organizations. Interestingly, these re-organizations appear to be independent of gene expression, yet they do not induce a broad-scale A/B compartment switching at the gene loci.

## DISCUSSION

Our new high-throughput multiomic single-cell technology, GAGE-seq, delivers an integrative approach to co-assay 3D genome structure and gene expression in individual cells with high resolution. We show that GAGE-seq can reveal complex cell types from complex tissues not identified by other existing methods. Additionally, its data integration with spatial transcriptomic data points to great potential to reach a deeper understanding of 3D genome variation within complex tissues. Importantly, GAGE-seq also facilitates the reconstruction of differentiation trajectories based on 3D genome features, transcriptomes, or both. The high congruence between these modalities underscores the intimate connection between the temporal variations of the 3D genome and transcriptional rewiring during cell differentiation. Notably, GAGE-seq has revealed much more nuanced relationships between 3D genome features and gene expression during bone marrow B-NK lineage differentiation, creating a resource for future studies to disentangle causal gene regulatory changes in differentiation through the lens of 3D genome in single cells.

GAGE-seq is characterized by its efficiency, scalability, robustness, cost-effectiveness, and adaptability. We envision that GAGE-seq, along with our analytical tools, could significantly enhance the current toolkit for single-cell epigenomics. With wide-ranging applications, GAGE-seq can deepen our understanding of genome structure and function, providing insights into normal development and disease pathogenesis. Future refinements, such as enhancing barcoding strategy for higher throughput and improving detection of chromatin contacts, may allow GAGE-seq to construct high-resolution cell atlases and assess the role of pathogenic noncoding single-nucleotide variants on chromatin loops^57^ in a massively parallel manner. Additionally, we anticipate a future application where GAGE-seq will be integrated with spatial labeling technologies, producing spatially-resolved scHi-C and scRNA-seq data. Such advancements will likely open up new avenues of investigation, such as exploring the role of the 3D genome in various tissue development and disease progression. Ultimately, GAGE-seq may offer the opportunity to coalesce multiscale molecular modalities in single cells, leading to a more comprehensive understanding of genome structure, cellular function, and their spatiotemporal variability.

## METHODS

### GAGE-seq experimental details

#### Preparation of 96-well plate of barcoded adaptors

Two separate barcoding rounds of ligation reactions are used in CARE-seq, with two different 96-well barcoding plates for each round (scHi-C and scRNA-seq, respectively, as detailed in **Table S1**). The design of the scRNA-seq part barcodes resembles that of Split-seq^42^ and SHARE-seq^43^ (**Table S1**). The molecular structure of the scHi-C part barcodes is depicted in **Fig. S1**.

#### Cell lysis

Crosslinked cells of K562, NIH3T3, GM12878, MDS-L, human bone marrow Cd34+ cells were thawed from −80°C or liquid nitrogen. 0.2 ml of high-salt lysis buffer 1 (50 mM HEPES pH 7.4, 1 mM EDTA pH 8.0, 1 mM EgTA pH 8.0, 140 mM NaCl, 0.25% Triton X-100, 0.5% IGEPAL CA-630, 10% glycerol, and 1× proteinase inhibitor cocktail (PIC)) was added per 1 × 106 cells. The cell solution was mixed thoroughly and incubated on ice for 10 min. After this, cells were pelleted at 500xg for 2 min at 4°C and then resuspended in 0.2 ml of high-salt lysis buffer 2 (10 mM Tris-HCl pH 8, 1.5 mM EDTA, 1.5 mM EgTA, 200mM NaCl, 1× PIC). The solution was incubated on ice for 10 min. Following this, cells were then pelleted at 500xg for 2 min at 4°C and then resuspended in 200 μl of 1 × T4 DNA ligase buffer (NEB, B0202S) containing 0.2% SDS. They are then incubated at 58°C for 10 min. To quench the reaction, 200 μl ice-cooled 1x NWB and 10 μl 10% Triton X-100 (MilliporeSigma, 93443) were added to the tube. Finally, cells were spun at 500xg for 4 min at 4°C. For crosslinked mouse brain cortex cells, the treatment was simplified. The step involving high-salt lysis buffer 1 and high-salt lysis buffer 2 was omitted, and 0.1% SDS was used for cell lysis.

#### Reverse transcription

SDS treated cells were resuspended in 400 μl of RT mix (final concentration of 1x RT buffer, 500 mM dNTP, 10 mM Biotinylated RT primers, 7.5% PEG 6000 (VWR, 101443-484), 0.4U/ml SUPERase•In™ RNase Inhibitor, and 25U/ml Maxima H Minus Reverse Transcriptase (ThermoFisher Scientific, EP0752)). The RT primers contain a poly dT tail, a biotin molecule, and a universal ligation overhang. The sample then underwent a series of heating cycles. Initially, it was heated at 50 oC for 10 minutes, then it went through 3 thermal cycles (8 °C for 12s, 15 °C for 45s, 20 °C for 45s, 30°C for 30s, 42 °C for 2 min and 50 °C for 3 min). Afterwards, the sample was again incubated at 50 oC for 10 minutes. After reverse transcription, 600 μl of 1x NWB was added, the sample was centrifuged at 500x g for 3 minutes, and the supernatant was then removed.

#### 1st-round chromatin fragmentation, proximity ligation, and 2nd-round chromatin fragmentation

Cells were resuspended in 400 μl of restriction enzyme (RE) digestion mix (1x T4 ligase buffer (NEB, B0202S), 500U MseI (NEB, R0525M), 240U CviQI (NEB, R0639L), 0.32 U/ml Enzymatics RNase Inhibitor, 0.05 U/ml SUPERase RNase Inhibitor), and incubated at room temperature (25 °C) for 2 hr. Cells were then centrifuged at 500x g for 3 minutes at 4 °C, and the supernatant was removed. The remaining cell pellet was washed twice with 300 μl of 1x NWB, and as much supernatant was removed as possible. Next, the pellet was resuspended in 200 μl of ligation mix (1x T4 ligation buffer (NEB, B0202S), 50 Units T4 DNA ligase (ThermoFisher Scientific, EL0012), 0.32 U/ml Enzymatics RNase Inhibitor, 0.05 U/ml SUPERase RNase Inhibitor) and incubated at 16 °C overnight. This was followed by adding 20 μl 10x T4 ligation buffer, 1 μl SUPERase RNase Inhibitor and 20 μl DdeI (NEB, R0175L). The sample was then incubated at 37 °C for 1 hr and centrifuged at 500x g for 3 minutes, with the supernatant removed afterwards.

#### Combinatorial cellular barcoding

Cells were resuspended in 330 μl of ligation mix (1x T4 ligase buffer (NEB, B0202S), 100 Units T4 DNA ligase (ThermoFisher Scientific, EL0012), 0.25 mg/ml BSA (ThermoFisher Scientific, AM2618), 5% PEG-4000 (ThermoFisher Scientific, EL0012), 0.32 U/ml Enzymatics RNase Inhibitor, 0.05 U/ml SUPERase RNase Inhibitor) and distributed into each well (3 μl/well) of the first-round barcoding plate, which already contained 2 μl of CARE-seq 1st-round adaptors in each well. This barcoding plate was then incubated at 25 °C for 3 hr. Afterwards, cells from all 96 wells were pooled into three 1.5 ml tubes, and 5 μl of 10% NP-40 (ABCam, ab142227) was added to each tube. This is followed by centrifuging at 500x g for 3 minutes at 4 °C. The supernatant was then removed and cells were resuspended in 300 μl 1x NWB containing 0.033% SDS and combined into one 1.5 ml tube. Cells were then pelleted at 500x g for 2 minutes at 4 °C. After three additional rounds of washing with 300 μl 1x NWB containing 0.033% SDS, cells were resuspended in 200 μl 1x NWB containing 0.1% SDS and filtered with with 10 μm or 20 μm cell ministrainer (PluriStrainer, 43-10010-50 or 43-10020-40). Cells were inspected under a microscope and counted with a hemocytometer. Approximately 7,500 cells were diluted with 1.25 ml of a dilution buffer containing 0.4x NEBuffer 2 (NEB, B7002S), 2 mg/ml BSA (ThermoFisher Scientific, AM2618), and 0.08 μM RNA ligation-1 block, and distributed into each well (3 μl/well) of a 96-well plate (the 2nd-round barcoding plate). Then, 2 μl of cell lysis buffer (5x NEBuffer 2, 0.625% SDS) were then added to each well of the 2nd-round barcoding plate. The plate was incubated at 60 °C for at least 24 hr.

For the 2nd-round barcoding, 1.5 μl of pre-mixed GAGE-seq adapters (0.2 μM Hi-C-AD2 and 0.17 μM RNA-AD2) were added to the plate, followed by 23.5 μl of ligation mix (3 μl 1x T4 ligase buffer (NEB, B0202S), 0.15 μl 50 mg/ml BSA (ThermoFisher Scientific, AM2618), 1 μl 10% Triton X-100 (MilliporeSigma, 93443), 0.03 μl 20 μM 5’-P-TNA-Nextera-P5-AD, 0.03 μl 20 μM 5’-P-TA-Nextera-P5-AD, 0.03 μl 10 μM RNA ligation-1 block, and 0.8 μl T4 DNA ligase (ThermoFisher Scientific, EL0012)). The ligation was carried out at 25 °C for 24 hr, and then stopped by adding 2 μl of proteinase K digestion mix (0.2 μl proteinase K (ThermoFisher Scientific, AM2546), 0.5 μl 10% SDS and 1.8 μl water) to each well. A reverse crosslinking was carried out at 60 °C for 20 hr.

#### Reverse crosslinking and separation of scHi-C and scRNA-seq libraries

After reverse crosslinking, the sample in each 96-well plate was pooled into 12 DNA low-binding 1.5 ml tubes (Eppendorf, 022431021). Genomic DNA (gDNA) and cDNA were precipitated by adding 66 μl 3M Sodium Acetate Solution (pH 5.2) (MilliporeSigma,127-09-3), 1 μl GlycoBlue (ThermoFisher Scientific, AM9515) and 720 μl iso-propanol (MilliporeSigma, I9516) to each tube, followed by incubating at −80 °C for at least 1 hr. The samples were then centrifuged at 15000 rpm for 10 min and the pellet in each tube were resuspended in 30 μl 1x NEBuffer 2 containing 0.15% SDS. After incubation at 37 °C for 10 min, the samples were combined into one DNA low-binding tube. gDNA and cDNA were precipitated by adding 66 μl 3M Sodium Acetate Solution (pH 5.2) and 720 μl iso-propanol, followed by incubating at −80 °C for at least 1 hr. The sample was then centrifuged at 15000 rpm for 10 min and the pellet was resuspended in 100 μl buffer EB (Qiagen, 19086). For each sample of a 96-well plate, 5.5 μl of MyOne C1 Dynabeads were washed twice with 1x B&W-T buffer (5mM Tris pH 8.0, 1M NaCl, 0.5mM EDTA, and 0.05% Tween 20) and resuspended in μl of 2x B&W buffer (10mM Tris pH 8.0, 2M NaCl, and 1mM EDTA) and added to the sample tube. The mixture was incubated at room temperature for 60 min and put on a magnetic stand to separate supernatant and beads.

#### scHi-C sequencing library construction

The supernatant that contained the Hi-C library (gDNA) was precipitated by adding 60 μl 3M Sodium Acetate Solution (pH 5.2) and 660 μl iso-propanol, followed by incubating at −80 °C for at least 1 hr. The sample was then centrifuged at 15000 rpm for 10 min and the pellet was washed with 0.8 ml 80% ethanol, air dried, and resuspended in 38.5 μl water. After adding 1.5 μl 20 μM 5’-P-TNA-Nextera-P5-AD, 1.5 μl 20 μM 5’-P-TA-Nextera-P5-AD, 1.5 μl 20 μM Hi-C-AD1-Block and 2 μl T4 DNA ligase, adaptor ligation was carried out at 22 °C for at least 20 hr and stopped by adding 2 μl 10% SDS. Hi-C library was purified by adding 80 μl Ampure beads and amplified in two 100 μl pre-amplification reactions (50 μl 2x NEBNEXT Ultra II Q5 Master Mix (NEB, M0544L), 5 μl 10 μM Nextera-P5-pre-Primer, 5 μl 10 μM Trueseq-P7-pre-P-S primer and 40 μl Hi-C sample), with the following PCR program: 98 °C for 2 min, and then 9 cycles at 98 °C for 15 s, 60 °C for 30 s and 65 °C for 3 min. The pre-PCR products were purified by 0.75x AMPure beads, eluted in 42 μl buffer EB, and quantified by Qubit (ThermoFisher Scientific). For Illumina sequencing library construction, about 40 ng purified pre-amplified Hi-C sample was fragmented in two 50 μl tagmentation mix (1x TD buffer and 0.5 μl TDE1 (Illumina Tagment DNA TDE1 Enzyme and Buffer Kit, 20034198)) at 55 °C for 5 minutes in two PCR tubes. Tagmentation was stopped by adding 15 μl NT buffer per well and incubated at room temperature for 5 min, followed by adding 60 μl NPM (Illumina, FC-131-1096), 12 μl 10 μM Indexed Nextera P5 primer, 12 μl 10 μM Indexed Trueseq P7 primer and 51 μl water. The final Hi-C sequencing library was amplified, with the PCR program: 72 °C for 5 min, 98 °C for 45 s, and then 7 cycles at 95 °C for 10 s, 55 °C for 30 s and 72 °C for 1 min. The amplified library was purified by 0.7x AMpure beads and eluted in 40 μl buffer EB.

#### scRNA-seq sequencing library construction

The Myone C1 beads with the cDNA library were resuspended in 100 μl of TdT mix (1x Terminal Transferase Reaction Buffer, 0.25 mM CoCl2, 1 µM dGTP/ddGTP mix (0.95 µM dGTP (Thermo Fisher Scientific, R0161), 0.05 µM ddGTP (MilliporeSigma, GE27-2045-01), 0.2 U/μl Terminal Transferase (NEB, M0315L)) and incubated at 37 °C for 20 min. The supernatant was removed by placing the sample on a magnetic stand. Beads were washed with 200 μl buffer EB and resuspended in 400 μl of Post-TdT PCR mix (1x NEBNEXT Ultra II Q5 Master Mix (NEB, M0544L), 0.5 μM post-TdT-poly(C)12-S Primer, and 0.5 μM post-TdT-P5-T7 primer). Post-Tdt PCR was carried out with the following program: 98 °C for 2 min, and then 13-18 cycles at 98 °C for 15 s, 52 °C for 45 s and 65 °C for 3.5 min. The PCR products were purified by 0.8x AMPure beads, eluted in 32 μl buffer EB, and quantified by Qubit. For sequencing library construction, about 1.5 ng purified pre-amplified cDNA sample was fragmented in four 20 μl tagmentation mix (10 μl 2x TD buffer, 5 μl Nextera XT (Illumina, FC-131-1096), and 5 μl cDNA sample) at 55 °C for 5 minutes in four PCR tubes. Tagmentation was stopped by adding 5 μl NT buffer per tube and incubated at room temperature for 5 min, followed by adding 15 μl NPM (Illumina, FC-131-1096), 3 μl 10 μM Indexed Nextera P7 primer, 3 μl 10 μM Indexed Trueseq P5 primer and 4 μl water to each tube. The final cDNA sequencing library was amplified with the PCR program: 72 °C for 5 min, 98 °C for 45 s, and then 12 cycles at 95 oC for 10 s, 55 °C for 30 s and 72 °C for 1 min. The amplified library was purified by 0.7x AMpure beads and eluted in 40 μl buffer EB.

#### Sequencing

Both scHi-C and scRNA-seq libraries were pooled and paired end sequencing (PE 150) were performed on the HiSeq, NextSeq, or NovaSeq platform (Illumina).

### GAGE-seq data processing workflow

#### Demultiplexing

DNA and RNA reads were assigned to wells based on the two rounds of barcodes. For DNA reads, only read 2 was used for demultiplexing, allowing at most 1 mismatch in each of the two rounds of barcodes. DNA reads with more than 5 mismatches in the region between the two rounds of barcodes (the 9th-23rd nt) were discarded. After demultiplexing, the first 12 nt were removed from read 1 and the first 35 nt were removed from read 2. For RNA reads, only read 1 was used for demultiplexing, allowing at most 1 mismatch in each barcode round. RNA reads with more than 6 mismatches in the region between the two rounds of barcodes (the 19th-48th nt) or with more than 6 mismatches in the region downstream of the first round of barcode (the 57th-71th nt) were discarded.

The two reference genomes were combined into a single reference genome file used for all GAGE-seq libraries. For DNA reads, BWA^58^ was used for alignment. The combined reference genome was indexed using command bwa index-a bwtsw. Paired, trimmed DNA reads were aligned to the combined reference genome using command bwa mem-SP5M. For RNA reads, STAR^59^ was used for alignment. The GENCODE annotation files for human (v36) and mouse (vM25) were downloaded and concatenated. The combined reference genome was indexed using command--runMode genomeGenerate--sjdbOverhang 100 with the combined gencode annotation file. Only read 2 of RNA reads was aligned with the command STAR--outSAMunmapped Within.

#### Identification of contact pairs from DNA reads

Pairtools^60^ was used to identify contact pairs from paired DNA reads with command pairtools parse--walks-policy all--no-flip--min-mapq=10. After that, walk reads (i.e., DNA reads containing multiple ligation sites) were further processed. Briefly, we assumed that any pair of loci in the same DNA read forms a valid contact pair, and these contact pairs were included in the results.

#### Deduplication of contact pairs

The contact pairs were deduplicated. We extract the genomic positions of the two ends of each contact pair. We define two contact pairs as directly duplicated if the two contact pairs’ first ends lie within 500 nt apart and their second ends also within 500 nt. If two contact pairs are not directly duplicated, but are directly or indirectly duplicated with a third contact pair, we define the first two contact pairs as indirectly duplicated. Among each cluster (i.e., connected component) of (in)directly duplicated contact pairs, the one with the largest difference between its two ends’ genomic positions was retained, and the rest were marked as duplicates.

#### Deduplication of RNA reads

The RNA reads were deduplicated. Two RNA reads are defined as directly duplicated if there is at most 1 mismatch in their UMI and if their genomic positions differ by at most 5 nt. The rest of the process is similar to the deduplication of contact pairs. Only one RNA read from each duplicate cluster is retained.

### GAGE-seq integrative analysis for mouse brain cortex

#### Integration with MERFISH data

Integration of GAGE-seq data and MERFISH data was done with Seurat^51^. Only scRNA-seq profiles from the GAGE-seq data were used for this integration. In the GAGE-seq mouse brain cortex data, the following cell types of excitatory neurons were used: L2/3 IT CTX a, L2/3 IT CTX b, L2/3 IT CTX c, L4 IT CTX, L4/5 IT CTX, L5 IT CTX, L6 IT CTX, L6 CT CTX a, L6 CT CTX b, L5/6 NP CTX, and L6b CTX. In the MERFISH data, cells from L2/3 IT, L4/5 IT, L5 IT, L5/6 NP, L6 CT, L6 IT, and L6b were used. Each time, the selected cells from GAGE-seq were integrated with one slice from the MERFISH data. All genes detected and expressed in both GAGE-seq and MERFISH were used. The ‘FindIntegrationAnchors’ and ‘IntegrateData’ functions were used with default parameters, except that the number of dimensions was set to 10.

#### Inference of whole-transcriptome expression and 3D genome features for MERFISH cells

The integrated single-cell expression profiles of GAGE-seq data and MERFISH data were scaled by the ‘ScaleData’ function from Seurat^51^ with default parameters, and the first 30 PCs were calculated by the ‘RunPCA’ function. A 50-nearest neighbor regressor was created to estimate whole-transcriptome expression and 3D genome features from the 30-dimensional PC space. The regressor was trained on GAGE-seq data and then applied to the MERFISH data. The Gaussian kernel was used as the weight function. For each MERFISH cell, the bandwidth was defined as the 0.3 quantile of the distances to the 50 nearest neighbors.

### GAGE-seq integrative analysis for bone marrow

#### Trajectory and pseudotime

The pseudotime of human bone marrow cells was inferred by the ‘sc.tl.diffmap’ and ‘sc.tl.dpt’ function in Scanpy^61^, jointly from the paired scRNA-seq profiles and scHi-C profiles. Specifically, cells in the HSC, MPP, MLP, and B-NK clusters were included. The first 5 PCs of the scRNA-seq profiles were used for the scRNA-based pseudotime and the first 2 PCs of the Fast-Higashi embeddings of the scHi-C profiles were used for the scHi-C-based pseudotime. The 5 scRNA-seq PCs and the 2 scHi-C PCs were then concatenated and used for the joint pseudotime. The ‘sc.pp.neighbors’ function was used to construct the neighbor graph with 30 (scRNA-based and joint pseudotime) or 20 (scHi-C-based pseudotime) nearest neighbors per cell. The ‘sc.tl.diffmap’ and ‘sc.tl.dpt’ function was applied with 10 diffusion components to learn a latent representation focusing on the trajectory and to infer the pseudotime for single cells. The origin of the trajectory was set based on the average expression level of HSC marker genes previously identified^54^.

#### Unsupervised clustering of genes

The clustering of genes was based on the expression and scA/B value. Genes expressed in at least 20 cells were included. To generate features for genes, 1) the expression levels and scA/B values were z-score normalized per gene among all cells. 2) cells were evenly divided into 10 bins based on the pseudotime, and 3) the average values of the expression and scA/B value in each bin were calculated for each gene. This process led to 20 features for each gene. The Louvain clustering algorithm was then applied to genes with 20 neighbors, a resolution of 1.5. The correlation was used as the distance metric.

Additional experimental methods, methods for quality control and benchmarking, methods for identifying single-cell 3D genome features, and other methods are described in the **Supplementary Methods**.

## Acknowledgements

This work was primarily supported by the National Institutes of Health (NIH) grant R01HG012303 (J.M. and Z.D.), with additional funding, in part, provided by NIH Common Fund 4D Nucleome Program grants UM1HG011593 (J.M.) and UM1HG011586 (Z.D.), NIH Common Fund Cellular Senescence Network Program grant UG3CA268202 (J.M.), and NIH grants R01HG007352 (J.M.) and R61DA047010 (Z.D.). Z.D. was additionally supported by EvansMDS Discovery Research Grant 2019. J.M. received additional support from a Guggenheim Fellowship from the John Simon Guggenheim Memorial Foundation, a Google Research Collabs Award, and a Single-Cell Biology Data Insights award from the Chan Zuckerberg Initiative. R.Z. was supported by the Eric and Wendy Schmidt Center at the Broad Institute.

## Author contributions

Z.D. and J.M. conceived and oversaw the project. Z.D. conceived and developed the GAGE-seq protocol with critical contributions from T.Z. and J.M.. T.Z. developed the computational workflow and performed all the data analysis with assistance from R.Z., under the supervision of Z.D. and J.M.. D.J. provided mice and dissected the mouse brain tissues, under the supervision of L.X.. R.T.D. and A.M. prepared human PBMCs for method optimization, under the supervision of J.L.A.. D.G. performed experiments under the supervision of Z.D. T.Z., Z.D., and J.M. wrote the manuscript with input from all authors.

