## Supplemental Information for "Concurrent profiling of multiscale 3D genome organization and gene expression in single mammalian cells"


**Supplementary Methods** **Page 2**

Additional Experimental Methods **Page 2**
Additional Computational Methods **Page 4**

**Supplementary Figures** **Page 10**

### Supplementary Methods

#### Additional Experimental Methods

**Cell culture.** K562 and GM12878 cells were cultured in RPMI 1640 as previously described[1]. NIH3T3 cells (CRL-1658, ATCC) were cultured at 37°C, 5% CO2 in DMEM/F12 supplemented with 1× Pen/Strep and 10% FBS. The myelodysplastic cell line MDS-L, which was a gift from Dr. Kaoru Tohyama (Kawasaki University of Medical Welfare
), was cultured in RPMI 1640 supplemented with 10% fetal bovine serum, 2.0 mM L-glutamine, 50 microM beta-Mercaptoethanol (MilliporeSigma) and 100 U/mL IL-3 (PeproTech) as previously described[2]. For crosslinking, K562, GM12878 and MDS-L cells growing at log phase were spun down at 500 xg for 2 min, resuspended in 10 mL serum-free RPMI-1640, crosslinked with a final concentration of 2% formaldehyde (Electron Microscopy Sciences, 15714) at room temperature (RT, 25°C) for 10 min. The crosslinking was quenched with 0.125M glycine on ice for 15 min, and cells were spun down at 500xg for 2 min, resuspended in 1X DPBS (ThermoFisher Scientific), aliquoted into 1 million cell aliquots, pelleted once again at 500 xg for 5 min, decanted, then snap frozen in liquid nitrogen, and finally stored at -80°C. The adherent NIH3T3 cells were washed once with 1X DPBS , trypsinized with 0.25% Trypsin-EDTA (ThermoFisher Scientific), spun down at 500 xg for 5 min., and resuspended in 10 mL serum-free DMEM/F12. The single-cell suspension was then subjected to crosslinking as described above.

**Mouse brain cortex.** All the mice used in this study received humane care in compliance with the principles stated in the Guide for the Care and Use of Laboratory Animals, NIH Publication, 1996 edition, and the protocols were approved by the Institutional Animal Care Committee (IACUC) at the University of Washington. The C57BL/6 mice were purchased from the Jackson Laboratory (Bar Harbor, ME) and housed in AAALAC-accredited vivarium. Room temperature was maintained at 22 ± 3°C and the humidity at 55 ± 15%. Animals were kept at a 12 h/12 h dark/light cycle. Adult C57BL/6 mouse brain cortex was dissected, snap-frozen, and stored in liquid nitrogen. Single nucleus suspensions were prepared from frozen cortex dissections (1.5–2 $mm^3$) as below. A piece of frozen cortex sample was placed in a pre-chilled 1.7ml DNA low-binding microtube containing 0.15 ml ice-cold EZ lysis buffer (MilliporeSigma, NUC101). The sample was pulverized 17 times with a pestle (Thomas Scientific, 1194N23) on ice, followed by 3 minutes incubation on ice. The sample was centrifuged at 500xg, 4°C for 2 min, and the pellet was resuspended in 1 ml ice-cold NSB-sucrose buffer (10 mM HEPES pH 8.0, 0.25 M sucrose, 10 mM KCl, 3 mM MgCl2, 1 µM DTT, 3 µL SupeRaseIn (ThermoFisher Scientific, AM2694) and 3 µL Enzymatic RNAse inhibitors (Qiagen, Y9240L)), and filtered through a 70-µm cell strainer (PluriStrainer). The single nuclei suspension was set at RT for 5 min, and 66.7μL 32% formaldehyde solution (Electron Microscopy Sciences, 15714) was added to the tube (2% final concentration). Crosslinking was carried out at RT for 10 min and stopped by adding 60 μL 2.5M glycine, 5 μL 10% NP-40, 10 μL 10% BSA, 3 μL SupeRaseIn and 3 μL Enzymatic RNAse inhibitor. Crosslinked nuclei were spun down at 500xg, 4°C for 5 min, and resuspended in 0.3 ml ice-cooled 1xNWB (10 mM Tris-HCl pH8.0, 10 mM NaCl, 0.025% NP40, 0.5 mM MgCl2, 0.2% BSA), plus 1 μL SupeRaseIn and 1 μL Enzymatic RNAse inhibitor. Nuclei were examined and counted after counterstaining with Trypan blue under a microscope. Finally, crosslinked nuclei were snap-frozen in liquid nitrogen and stored at -80$^{o}$C.

**Human bone marrow CD34+ cells.** Primary human bone marrow CD34+ cells were purchased from ATCC (PCS-800-012, Lot: 80926212 and 80225277), which were purified from healthy donors with a monoclonal CD34 antibody (>=90% purity). For crosslinking, a vial of cells (~500,000 cells) were thawed at 37$^{o}$C for 2 min, transferred to an ice-cooled 1.5ml DNA low-binding tube, and centrifuged at 400xg, 4$^{o}$C for 6 min. The cells were resuspended in 1ml SFEM II (Stemcell, 09605), supplemented with StemSpan™ CD34+ Expansion Supplement (Stemcell, 02691) and UM729 (Stemcell, 72332), and incubated at 37$^{o}$C for 10 min in a $CO_2$ incubator. The cells were then set at RT for 5 min, crosslinked with 57μL 32% formaldehyde solution (Electron Microscopy Sciences, 15714) for 10 min at RT, and quenched with 60 μL 2.5M glycine, 3 μL SupeRaseIn and 3 μL Enzymatic RNAse inhibitor. The crosslinked cells were spun down at 400xg, 4°C for 6 min and immediately subjected to GAGE-seq data generation.

#### Additional Computational Methods

##### Quality control and benchmarking

**Filtering cells for estimating the collision rate.** Wells were first filtered with a less stringent set of criteria for estimating the collision rate. For the 3 cell line libraries, wells with fewer than 1K RNA reads from any species or fewer than 10K contact pairs of any species were removed. For the K562-NIH3T3 mixture library, additional filtering based on species purity was applied: wells were retained if at least 95% of RNA reads were mapped to the human genome or at least 95% of RNA reads were mapped to the mouse genome, and if at least 98% of contact pairs were mapped to the human genome or at least 99% of contact pairs were mapped to the mouse genome. A well was annotated as a doublet if the well had sufficient numbers of contact pairs and RNA reads but did not pass the species purity filter. For the 3 mouse brain cortex libraries, wells with fewer than 1K RNA reads mapped to the mouse genome or fewer than 50K contact pairs mapped to the mouse genome were removed. For the 3 human bone marrow libraries, wells with fewer than 100 RNA reads mapped to the human genome or fewer than 40K (for replicate 1), 50K (for replicate 2), or 100K (for replicate 3) contact pairs mapped to the human genome were removed.

**Filtering cells for downstream analysis.** Wells were then filtered with more stringent criteria for identifying high-quality cells for subsequent analysis. For the K562/NIH3T3 mixture experiment, a well is assigned to the K562 library (K562 replicate 1) if the well had 1) at least 5K human RNA reads, 2) between 100-7,000 human genes, 3) at least 90% of RNA mapped to the human genome, 4) at most 15% of RNA originating from human mitochondria, 5) at least 40K human contact pairs, 6) at least 90% of contact maps mapped to the human genome, 7) at least 1 contact pair per 1M bases on average on each human chromosome, 8) at least 0.5 nonzero elements per locus on average in the 500kb contact map on each human chromosome.

For the K562/NIH3T3 mixture library, wells were assigned to the NIH3T3 library if it had: 1) at least 5K mouse RNA reads, 2) between 100-4,500 mouse genes, 3) at least 95% of RNA mapped to the mouse genome, 4) at most 15% of RNA coming from mouse mitochondria, 5) at least 100K mouse contact pairs, 6) at least 99% of contact maps mapped to the mouse genome, 7) at least 1 contact pair per 1M bases on average on each mouse chromosome, 8) at least 2 nonzero elements per locus on average in the 500kb contact map on each mouse chromosome.

For the K562/GM12878 mixture experiment, wells were retained if the well had: 1) at least 5K human RNA reads, 2) between 100-7,000 human genes, 3) at least 90% of RNA mapped to the human genome, 4) at most 15% of RNA coming from human mitochondria, 5) at least 40K human contact pairs, 6) at least 90% of contact maps mapped to the human genome, 7) at least 1 contact pair per 1M bases on average on each human chromosome, 8) at least 0.5 nonzero elements per locus on average in the 500kb contact map on each human chromosome. Retained wells were then assigned to either the K562 library (K562 replicate 2) or the GM12828 library based on the first-round barcode.

For the MDS-L library, wells were retained if the well had: 1) at least 5K human RNA reads, 2) between 100-6,000 human genes, 3) at least 90% of RNA mapped to the human genome, 4) at most 5% of RNA coming from human mitochondria, 5) at least 30K human contact pairs, 6) at least 90% of contact maps mapped to the human genome, 7) at least 1 contact pair per 1M bases on average on each human chromosome, 8) at least 1 nonzero elements per locus on average in the 500kb contact map on each human chromosome.

For the 3 mouse brain cortex library, wells were retained if the well had: 1) at least 1K mouse RNA reads, 2) at most 1% of RNA coming from mouse mitochondria, 3) at least 50K mouse contact pairs, 4) at least 20 contact pair per 1M bases on average on each mouse chromosome.

For the 3 human bone marrow library, wells were retained if the well had: 1) at least 1K human RNA reads, 2) 30-10,000 human genes, 3) at most 10% of RNA coming from human mitochondria, 4) at least 40K (for replicate 1), 50K (for replicate 2), or 100K (for replicate 3) human contact pairs, 5) 10-1,000 contact pair per 1M bases on average on each human chromosome.

**Doublet removal.** In order to thoroughly remove doublets for the mouse brain cortex dataset and the human bone marrow dataset, the DoubletDetect tool (https://doi.org/10.5281/zenodo.6349517) was utilized on the 3 mouse brain cortex libraries and the 3 human bone marrow libraries. For the 3 mouse brain cortex libraries, the BoostClassifier was trained with parameters n_iters=100, n_components=28, clustering_algorithm="louvain", standard_scaling=True, pseudocount=1. Doublets were then inferred by the trained classifier with thresholding parameters p_thr=1e-2, v_thr=.3. For the 3 human bone marrow libraries, the BoostClassifier was trained with parameters n_iters=100, n_components=9, clustering_algorithm="louvain", standard_scaling=True, pseudocount=1. Doublets were then inferred by the trained classifier with thresholding parameters p_thr=1e-2, v_thr=.3. After this, cells with more than 45K nonzero elements in the contact map were removed from the 3 mouse brain cortex libraries.

**Estimation of scHi-C library size for GAGE-seq and HiRES.** The total scHi-C library size for GAGE-seq and HiRES was estimated using the Lander-Waterman equation[3], based on the number of observed unique reads and the number of sequenced reads. The Lander-Waterman equation was solved with the data right after the deduplication step. The proportion of retained contact pairs in subsequent filter steps was calculated and the solution to the Lander-Waterman equation was scaled down by the proportion.

#### Analysis methods for revealing single-cell 3D genome features

**Calculation of CpG-based scA/B compartment values for raw single-cell contact maps.** The calculation of CpG-based scA/B compartment values for single-cell contact maps was conducted in a way similar to the method in Tan *et al.*[4,5]. Specifically, we calculate the A/B value one locus at a time. For each locus, we 1) extract the corresponding row vector from the contact map, 2) normalize the row vector by the row sum, 3) take the inner product of the normalized row vector and the CpG densities. The inner product is defined as the scA/B compartment value for that locus. Unless otherwise stated, CpG-based scA/B values were used for mouse brain cortex libraries.

**Calculation of eigenvector-based A/B compartment values for bulk and pseudo-bulk contact maps.** The calculation of eigenvector-based A/B compartment values for bulk and pseudo-bulk contact maps was performed in a way similar to the method in[6]. Specifically, we 1) perform the SQVC and O/E normalizations, 2) calculate the pairwise Pearson correlation between rows, 3) perform PCA on the square Pearson correlation matrix, 4) extract the first PC, 5) z-score normalize the first PC. The A/B compartment values were defined as the values in the normalized first PC.

**Calculation of scA/B compartment values for Higashi-imputed single-cell contact maps.** The scA/B compartment values for Higashi-imputed contact maps were calculated using the 'scCompartment.py' script in the Higashi suite[7]. The CpG density was used as the calibration file. Contact maps imputed without neighbor information were used. The z-score normalized scA/B values were used. Due to the stochastic nature of chromatin structure and the limited number of cells in the analysis on the human bone marrow dataset, we utilized the Higashi suite to enhance the contact maps. Unless otherwise stated, these scA/B values were used for human bone marrow libraries.

**Calculation of insulation scores.** To calculate the insulation score for each locus on a contact map at a certain resolution, two quantities were defined for each locus. The first quantity is the number of contact pairs whose both ends are at most $h$ loci away from the locus of interest, where $h$ is a hyperparameter and is called the half window size. The second quantity is the number of contact pairs that are counted toward the first quantity and have the two ends at different sides of the locus of interest. The insulation score of a locus is then defined as the second quantity divided by the first quantity. Unless otherwise stated, this insulation score was used for single-cell and pseudo-bulk contact maps in mouse brain cortex libraries and for pseudo-bulk contact maps in human bone marrow libraries.

**Calculation of insulation score for Higashi-imputed single-cell contact maps.** The insulation scores for Higashi-imputed contact maps were calculated by the ‘scTAD.py’ script in the Higashi suite with parameters --window_ins 2000000 --window_tad 500000. Contact maps imputed without neighbor information were used. Unless otherwise stated, this insulation score was used for single-cell human bone marrow libraries.

**Calculation of gene body score.** The gene body score was calculated as the values along the main diagonal line divided by the total number of contacts in this contact map.

**Dimension reduction of scHi-C profiles by Fast-Higashi.** Fast-Higashi[8] was used for inferring a low-dimensional representation of single-cell Hi-C contact maps. Fast-Higashi was run on the autosome, using the first 100 diagonal lines, without filter or convolutional imputation. The partial RWR imputation was enabled for the human bone marrow dataset due to the limited number of cells used in the analysis, but not for any cell line datasets or the mouse brain cortex dataset. ‘do_col’ and ‘no_col’ were set to false so that Fast-Higashi automatically decided whether to shift columns. The rank of the embedding space was set to 128 for the mouse brain cortex dataset, 64 for the human bone marrow dataset, and 32 for the K562 dataset. The convergence criterion for the main optimization procedure was set to ‘tol=1e-5’. After convergence, the ‘embed_l2_norm_correct_coverage_fh’ embedding was extracted as the single-cell low-dimensional representation.

**Additional analysis methods for the cell line analysis**

**Cell cycle in K562.** Cell cycle stages were inferred from the Fast-Higashi embedding of scHi-C profiles of the K562 dataset. The Louvain clustering algorithm implemented in Scanpy[9] was performed on the Fast-Higashi embedding with 20 neighbors and a resolution of 1. Every Louvain clustering was annotated to a cell cycle stage based on the percentage of short-range contacts and mitosis-band contacts.

**Additional analysis methods for the mouse cortex GAGE-seq dataset**

**Unsupervised clustering of scRNA-seq profiles.** The Seurat[10] package was used for clustering cells based on their scRNA-seq profiles. The single-cell count matrix of high-quality, non-doublet cells were loaded into the analysis. Only genes expressed in at least 10 cells were kept for subsequent analysis. The expression data were then normalized using the ‘NormalizeData’ and ‘ScaleData’ functions with default parameters. Highly variable genes were identified with the ‘FindVariableFeatures’ function. PCA analysis was performed with the ‘RunPCA’ function with npcs=40 PCs. The number of significant PCs was determined by the

‘JackStraw’ function with 1,000 replicates on the first 30 PCs. The Louvain clustering was performed with functions ‘FindNeighbors’ and ‘FindClusters’ using 20 neighbors, the first 27 PCs, the euclidean distance as the metric, and a resolution of 3. The Louvain clustering step was repeated by 100 times, each running up to 100 iterations until convergence. The best clustering results were kept by the ‘FindClusters’ function. After the clustering step, one cell cluster was found to not express the marker genes of any known cell types, and was therefore annotated as an unknown cluster.

**Unsupervised clustering of scHi-C profiles.** The Louvain clustering on scHi-C profiles was performed with the Scanpy package. For clustering all cells, 20 neighbors and a resolution of 1 were used. For clustering excitatory neurons and inhibitory neurons, 15 neighbors and a resolution of 1.5 were used.

**Identification of differentially expressed genes.** Differentially expressed genes (DEGs) were identified using the MAST[11] algorithm in the Seurat package, unless otherwise stated. The ‘FindMarkers’ function from Seurat was used, without filtering genes by fold-change or the percentage of cells expressing the gene. The default Bonferroni correction was used for correcting p-value for multiple hypothesis testing. For the analysis enabled by the integration with spatial transcriptomic data, marker genes were identified with the ‘rank_genes_groups’ function from Scanpy. Wilcoxon test was used and DEGs were filtered by the ‘filter_rank_genes_groups’ function with parameters min_fold_change=1, min_in_group_fraction=.1, max_out_group_fraction=1.

**Selection of representative gene loci.** To showcase the relationship between expression and 3D genome structures, representative gene loci were selected based on expression, scA/B value, and single-cell insulation score. Specifically, a gene is retained if it 1) over-expressed in inhibitory neurons than in excitatory neurons (*P*-value < 1e-2 and log2(fold-change) > 0.2), 2) has higher scA/B value in inhibitory neurons than in excitatory neurons (t-test *P*-value < 1e-2), 3) is over-expressed in Pvalb than in other inhibitory neurons (*P*-value < 1e-2 and log2(fold-change) > 0.2), and 4) has lower single-cell insulation score in Pvalb than in other inhibitory neurons (t-test *P*-value < 1e-2). Among the remaining genes, the one with the longest gene body was selected for further analysis.

**Identification of chromatin loops.** Chromatin loops were identified from contact maps at the 10kb resolution for each subtype. We utilized the loop calling pipeline included in the scHiCluster software with the default parameters[12], where single cell contact maps at 10kb are first imputed with linear convolution and random-walk-with-restart and are further normalized by distance decay. The differences between the normalized contact maps and their locally smoothed versions are used for a paired *t*-test across all single cells[13]. All entries with FDR less than 0.1 are considered as loop candidates. Adjacent loops, defined as those with a total anchor difference of 20kb or less, were merged, with the merged loop's strength calculated as the average pixel values with the loop.

**Identification of chromatin loops for the Erbb4 locus.** To investigate the correlation of chromatin loops and expression of the *Erbb4* gene, chromatin loops were filtered based on the genomic locations and correlation with the expression of *Erbb4*. Specifically, a chromatin loop was retained if 1) both of its anchors were between 66-71.2 Mb on chromosome 1, and 2) its second anchor (the anchor with the largest genomic location) was between 69.08-69.14 Mb on chromosome 1 (around the TSS of *Erbb4*). For each of the remaining chromatin loops, the correlation between loop strength and expression was calculated, and the only chromatin loop with a significant positive correlation was selected.

#### Additional analysis methods for the bone marrow GAGE-seq dataset

**Cell cycle.** The cell cycle stages for human bone marrow cells were inferred with the 'score_genes_cell_cycle' from Scanpy. Genes from Tirosh et al.[14] were used. Cells at the G1 and G0 stages were kept for downstream analysis.

**Unsupervised clustering of scRNA-seq profiles.** Clustering of the scRNA-seq profiles of the human bone marrow dataset was performed using Scanpy. Genes expressed in at least 10% of cells were used for PCA. A 15-nearest neighbor graph was constructed based on the first 7 PCs. The Louvain clustering algorithm was then performed with a resolution of 1. Clusters that did not highly express any known marker gene sets were annotated as unknown.

### Supplementary Figures

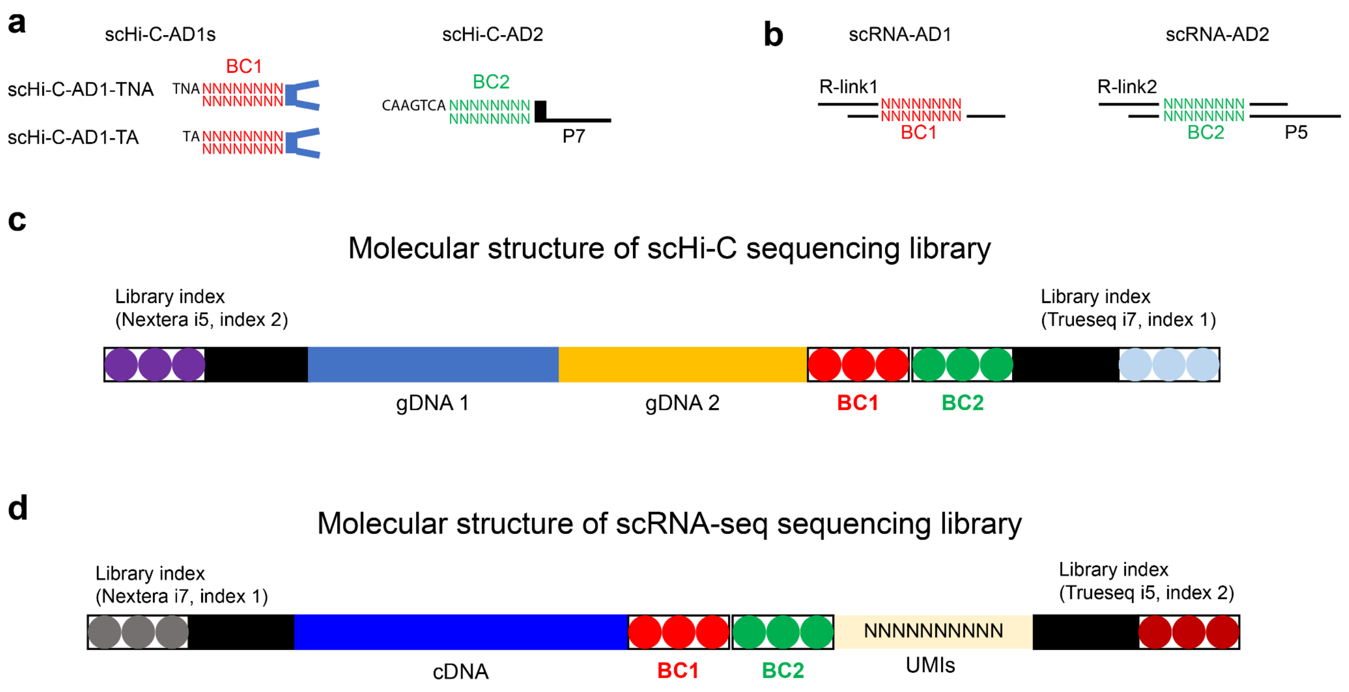


**Figure S1. Molecular design of GAGE-seq adaptors and the molecular structure of the DNA fragments in GAGE-seq scHi-C and scRNA libraries. a-b**. The structure of the two-round barcoded adaptors used in scHi-C (a) and scRNA-seq (b). **c-d**. The overall molecular structure of the DNA fragments in GAGE-seq scHi-C **(c)** and scRNA -seq **(d)** libraries.

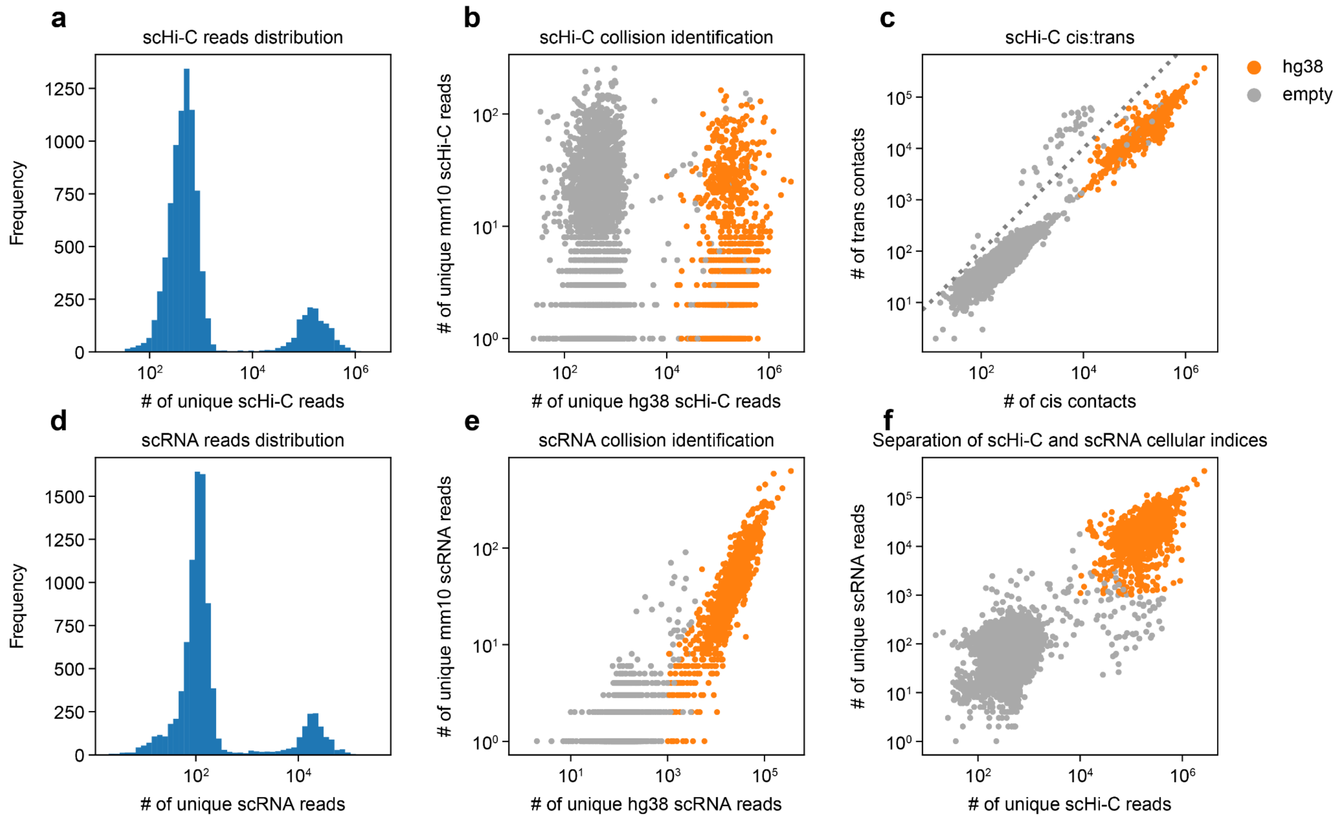


**Figure S2. Quality-control assessment of the K562-GM12878 GAGE-seq libraries**. **a** and **d**. Histogram showing the binomial distribution of GAGE-seq scHi-C (a) and scRNA-seq reads (d). **b** and **e**. Scatter plots representing the collision level in the GAGE-seq scHi-C (b) and scRNA-seq (e) libraries. **c**. Scatter plot showing the cis:trans ratio of scHi-C reads. **f**. Scatter plot indicating the clear separation of DNA and RNA reads of valid cellular indices from those of empty indices. Human data are colored in orange and empty indices in gray.

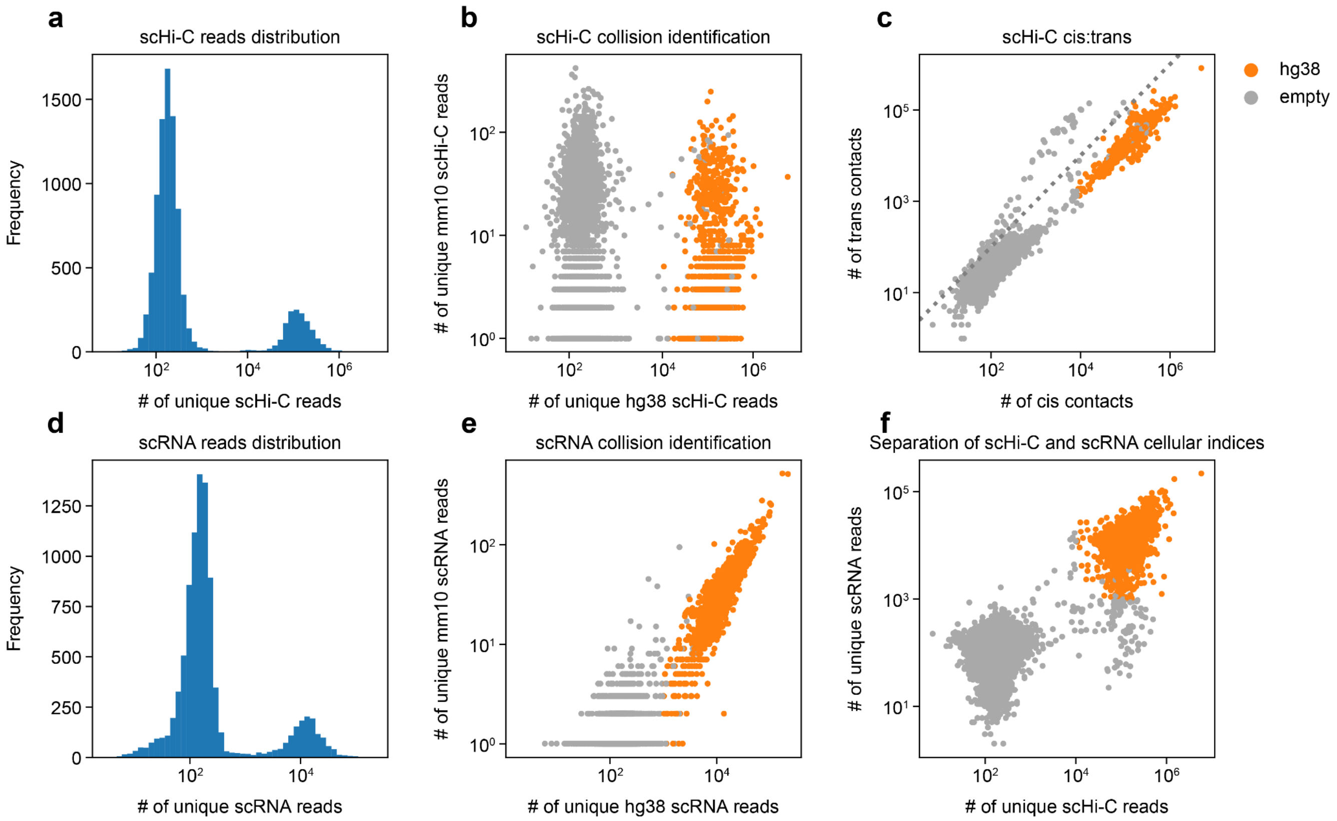


**Figure S3. Quality-control assessment of the MDS-L GAGE-seq library.** **a** and **d**. Histogram showing the binomial distribution of GAGE-seq scHi-C (a) and scRNA reads (d). **b** and **e**. Scatter plots showing the collision level in the GAGE-seq scHi-C (b) and scRNA-seq (e) libraries. **c**. Scatter plot showing the cis:trans ratio of scHi-C reads. **f**. Scatter plot indicating the clear separation of DNA and RNA reads of valid cellular indices from that of empty indices. Human data are colored in orange and empty indices in gray.

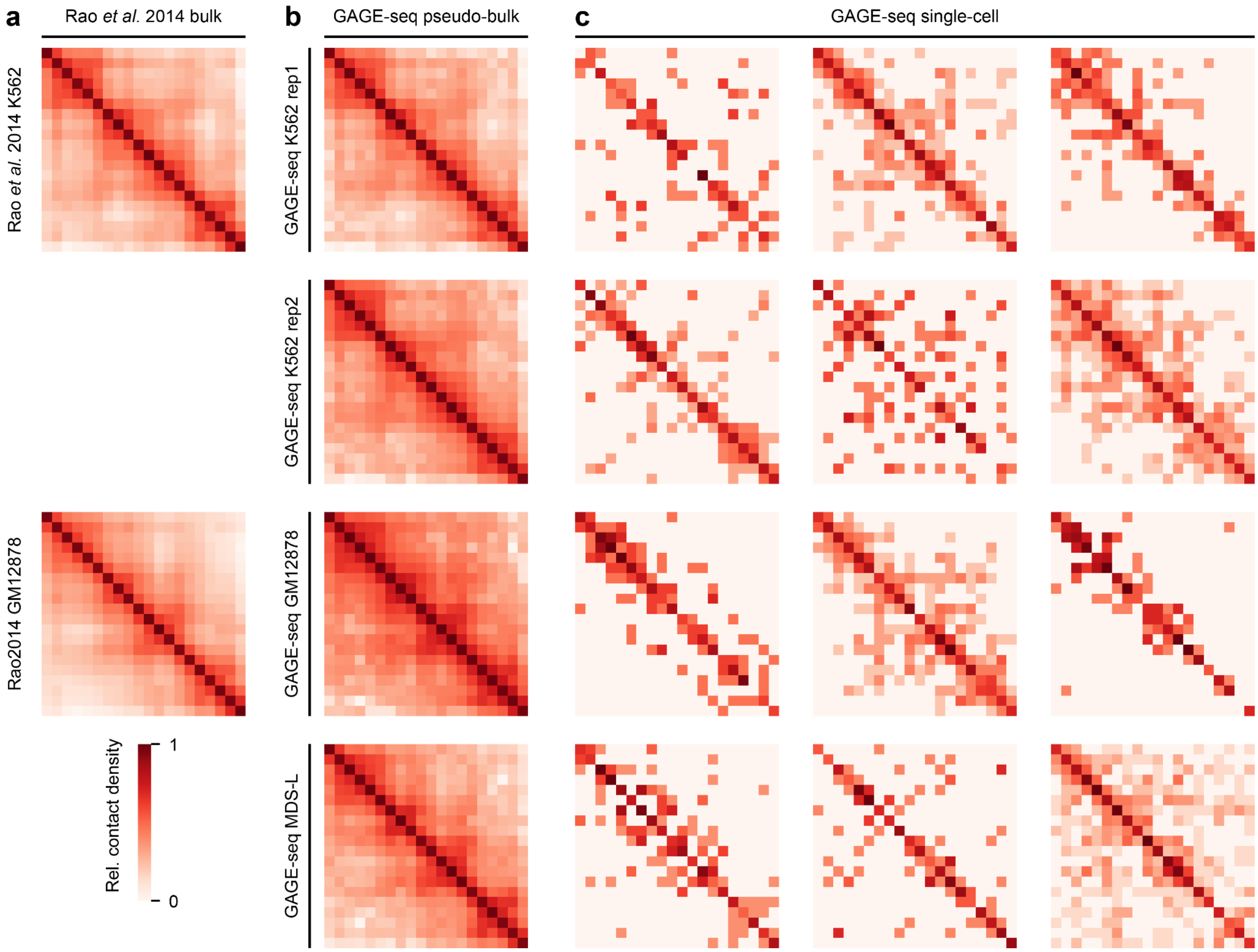


**Figure S4. Single-cell and pseudo-bulk contact maps from the GAGE-seq datasets at the beta globin locus. a**. Benchmarking contact maps from Rao et al[6]. **b.** GAGE-seq pseudo-bulk contact maps. **c**. Representative GAGE-seq single-cell contact maps. The displayed genomic location is human chr11:4.5 - 6.5 Mb, and the resolution is 100 Kb.

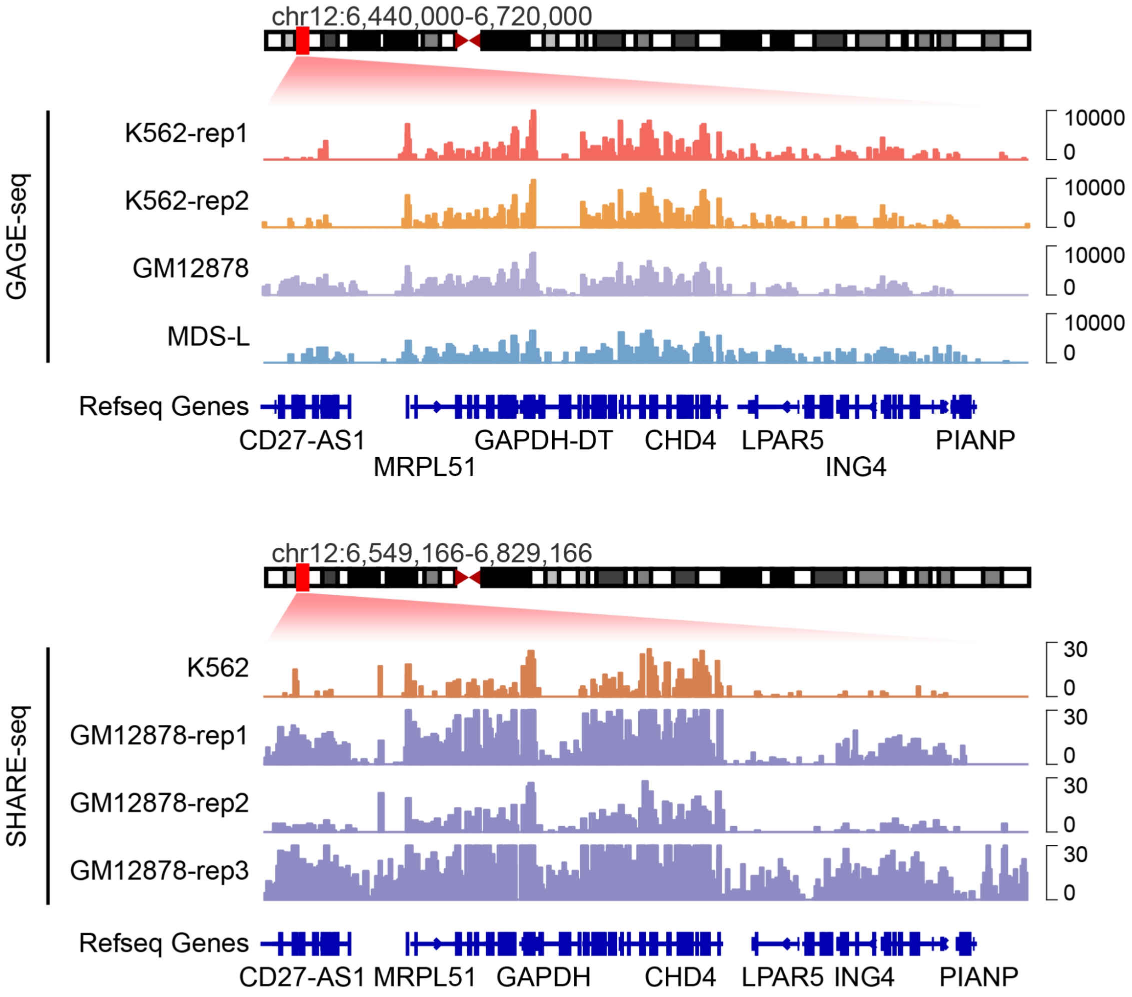


**Figure S5 Aggregated single-cell gene expression profiles of the genes in the GAPDH locus.** Upper panel: scRNA-seq signals from GAGE-seq libraries of K562, GM12878, and MDS-L cells (hg38). Lower panel: scRNA-seq signals from SHARE-seq in K562 and GM12878 cells (hg19)[15].

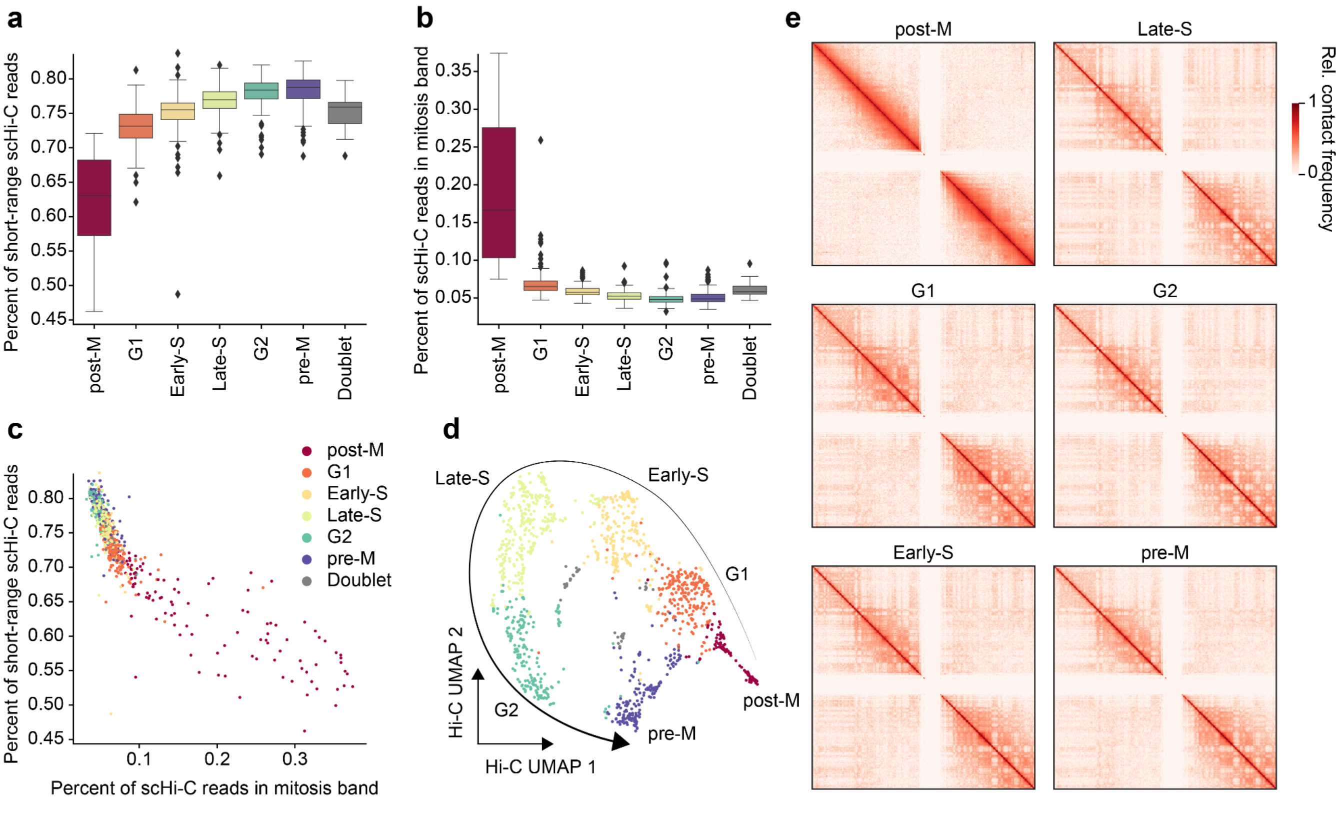


**Figure S6. Cell cycle analysis of the GAGE-seq K562 cells. a**. Percentage of short-range contacts (< 2 Mb) in single cells. **b**. Percentage of mitosis-band contacts (2 to 12 Mb) in single cells. **c**. Joint visualization of the percentage of short-range and mitosis-band contacts in single cells. **d**. UMAP visualization of the Fast-Higashi embeddings. **e**. Aggregated contact maps of the 6 inferred cell cycle phases on chromosome 1 at 1 Mb resolution.

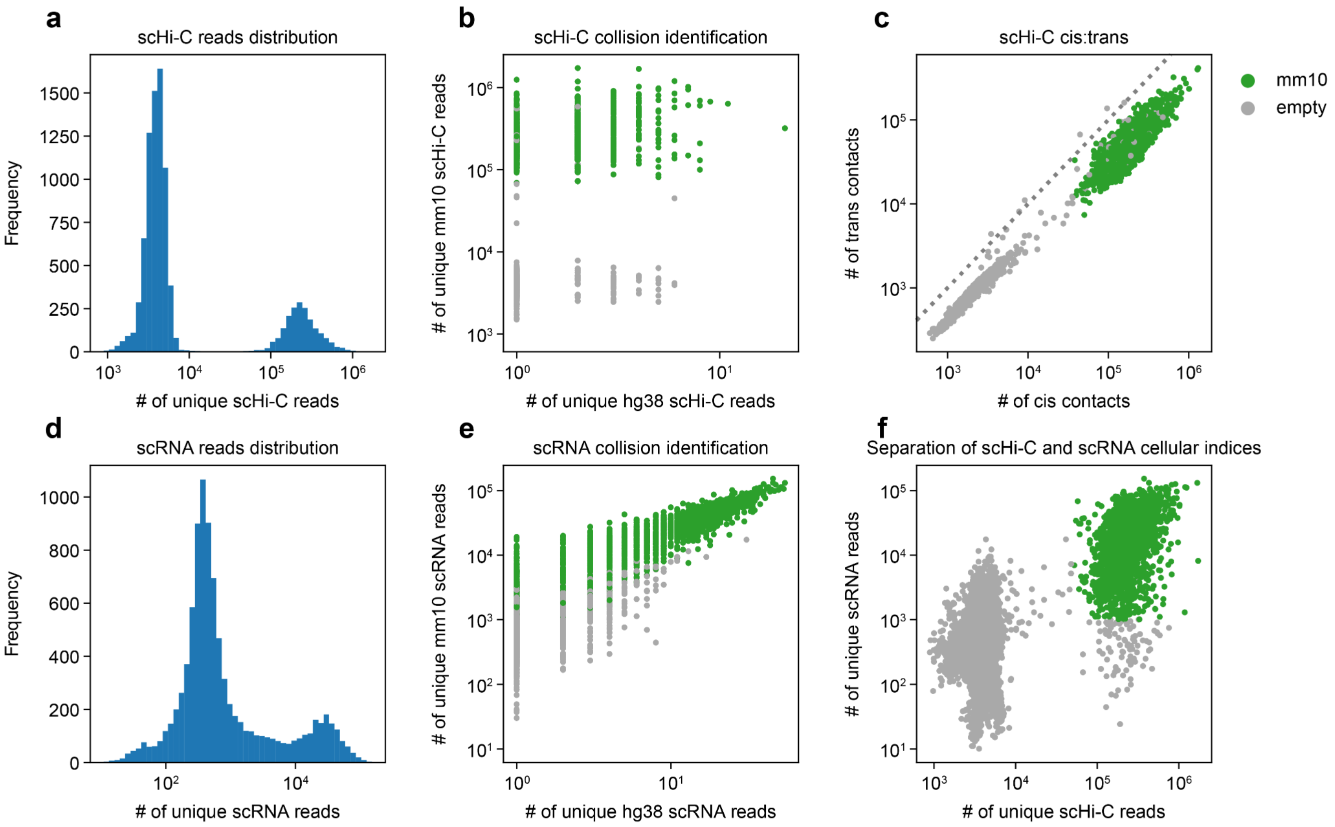


**Figure S7. Quality-control assessment of the GAGE-seq mouse brain cortex library (replicate 1). a** and **d**. Histogram showing the binomial distribution of GAGE-seq scHi-C **(**a**)** and scRNA-seq reads **(**d**)**. **b** and **e**. Scatter plots showing the collision level in the GAGE-seq scHi-C **(**b**)** and scRNA-seq **(**e**)** libraries. **c**. Scatter plot showing the cis:trans ratio of scHi-C reads. **f**. Scatter plot showing the clear separation of DNA and RNA reads of valid cellular indices from that of empty indices. Mouse data are colored in green and empty indices in gray.

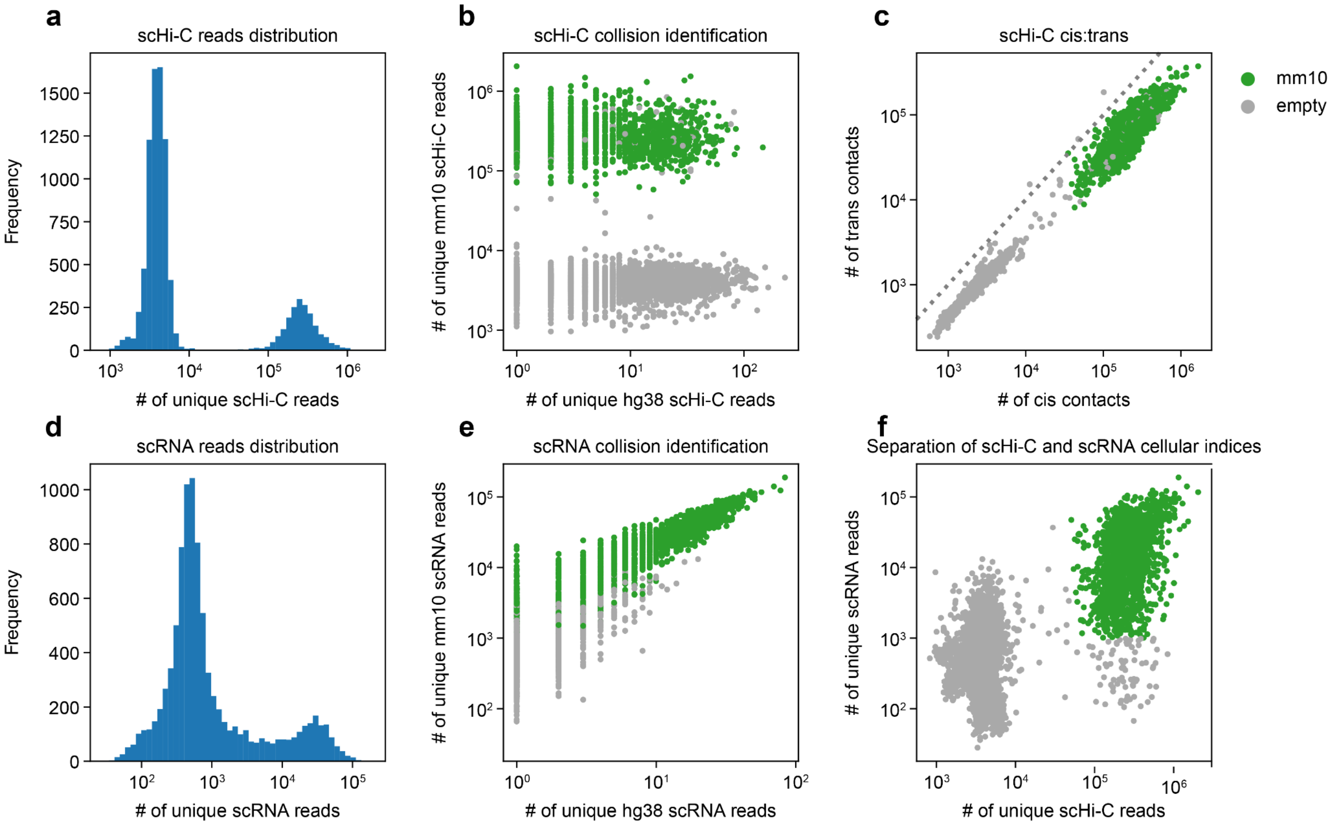


**Figure S8. Quality-control assessment of the GAGE-seq mouse brain cortex library (replicate 2). a** and **d.** Histogram showing the binomial distribution of GAGE-seq scHi-C **(a)** and scRNA-seq reads **(d)**. **b** and **e**. Scatter plots showing the collision level in the GAGE-seq scHi-C **(b)** and scRNA-seq **(e)** libraries. **c**. Scatter plot showing the cis:trans ratio of scHi-C reads. **f**. Scatter plot showing the clear separation of DNA and RNA reads of valid cellular indices from that of empty indices. Mouse data are colored in green and empty indices in gray.

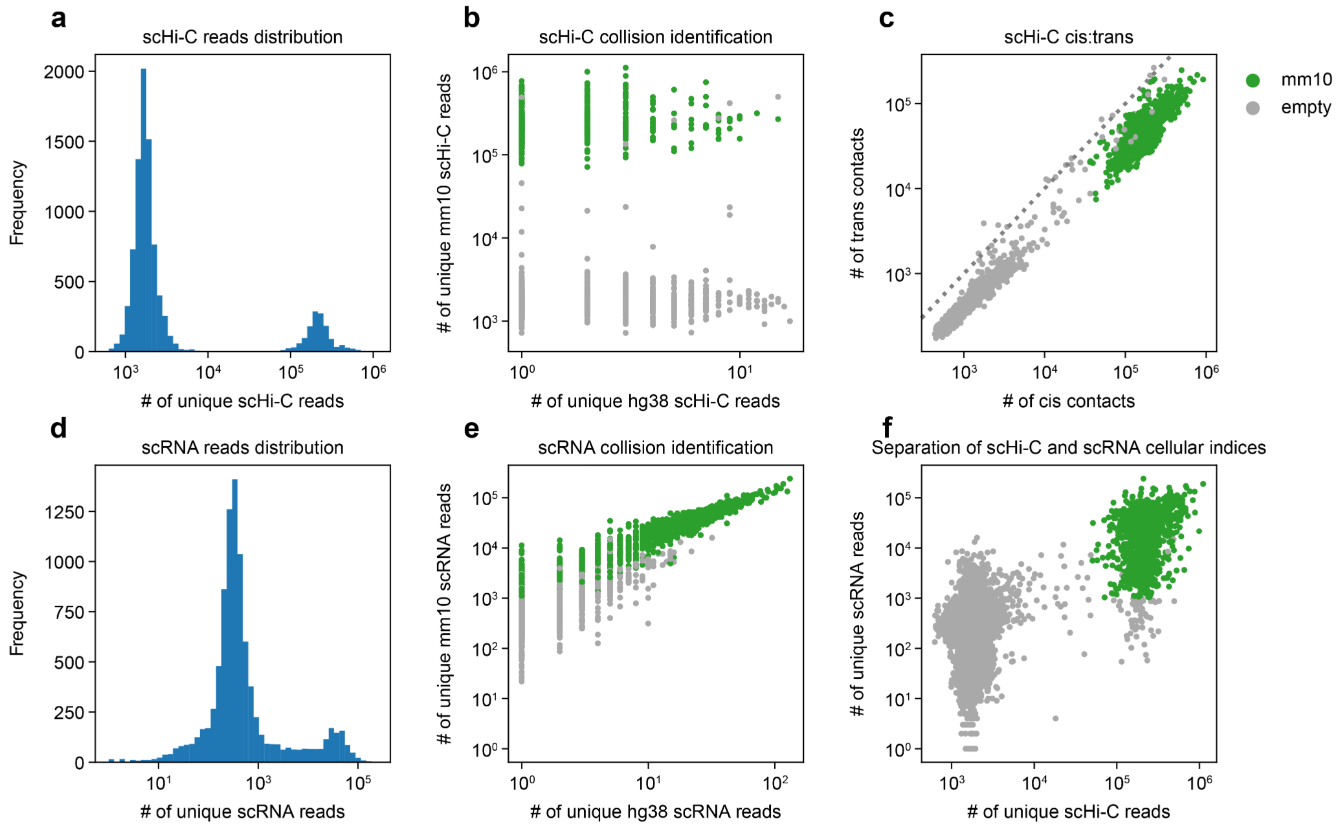


**Figure S9. Quality-control assessment of the GAGE-seq mouse brain cortex library (replicate 3). a** and **d.** Histogram showing the binomial distribution of GAGE-seq scHi-C **(a)** and scRNA-seq reads **(d)**. **b** and **e**. Scatter plots showing the collision level in the GAGE-seq scHi-C **(b)** and scRNA-seq **(e)** libraries. **c.** Scatter plot showing the cis:trans ratio of scHi-C reads. **f**. Scatter plot showing the well-separation of DNA and RNA reads of valid cellular indices from that of empty indices. Mouse data are colored in green and empty indices in gray.

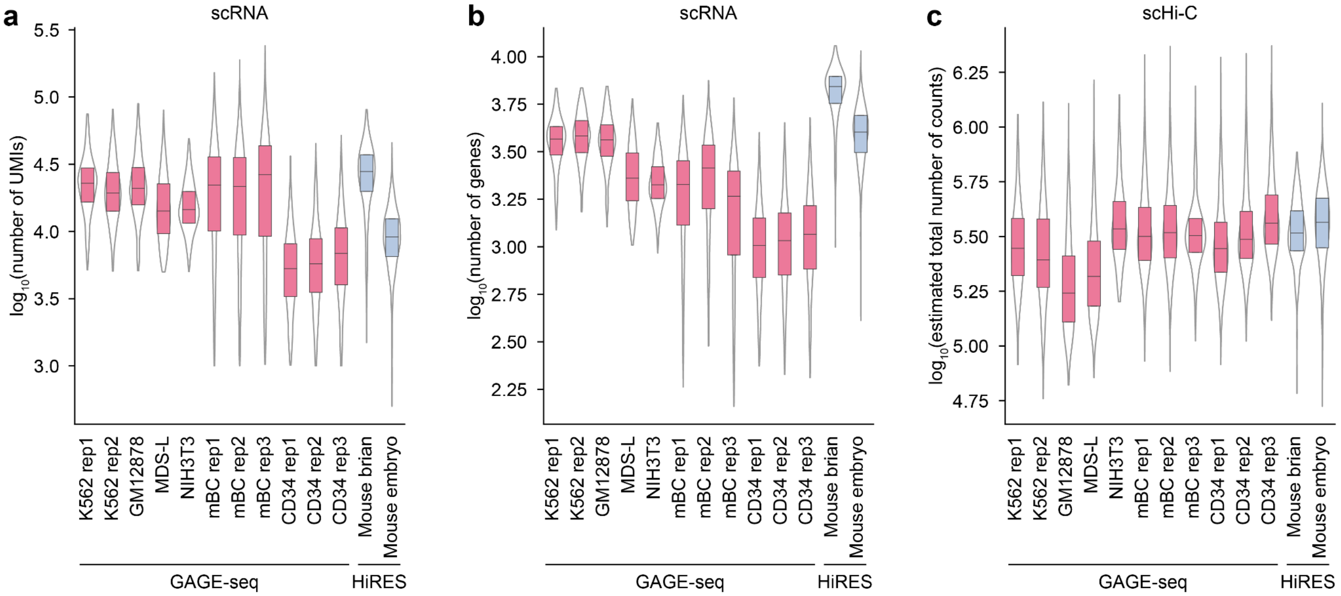


**Figure S10. Comparison of estimated scRNA and scHi-C library complexities between GAGE-seq and HiRES[16]. a**. The number of UMIs detected in single cells. **b**. The number of genes detected in single cells. **c**. The estimated number of chromatinontacts detected in the scHi-C libraries.

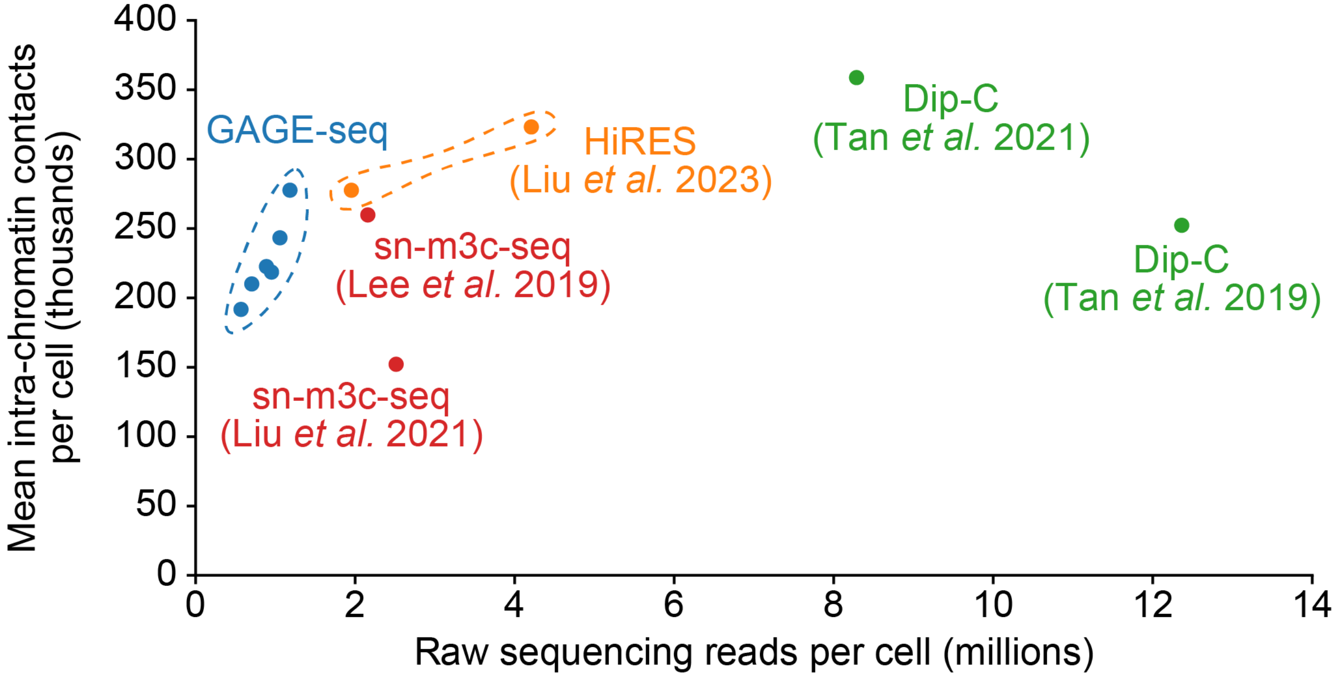


**Figure S11. Comparison between GAGE-seq and other scHi-C related methods in terms of efficiency.** We show intra-chromosomal chromatin contacts per single cell against sequence depth between GAGE-seq scHi-C (blue), Dip-C (green)[4,5] and sn-m3C-seq (red)[17,18] and HiRES (orange)[16]. The six blue dots represent the GAGE-seq libraries with mouse brain cortex and human CD34+ cells from this work.

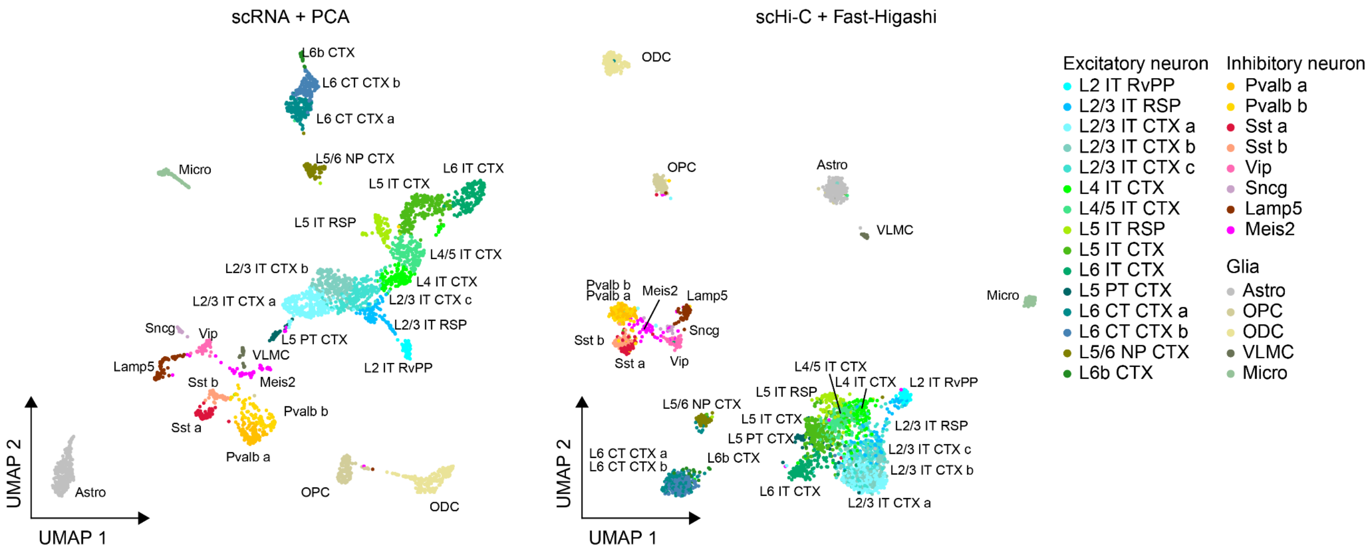


**Figure S12. High resolution cell type identification in the mouse brain cortex using GAGE-seq.** Left panel: UMAP visualization of the PCA embeddings of GAGE-seq scRNA-seq profiles. Right panel, UMAP visualization of the Fast-Higashi embeddings of GAGE-seq sHi-C profiles. A strong correlation between the “structure” (scHi-C-based) and transcriptome (scRNA-based) cell types is observed.

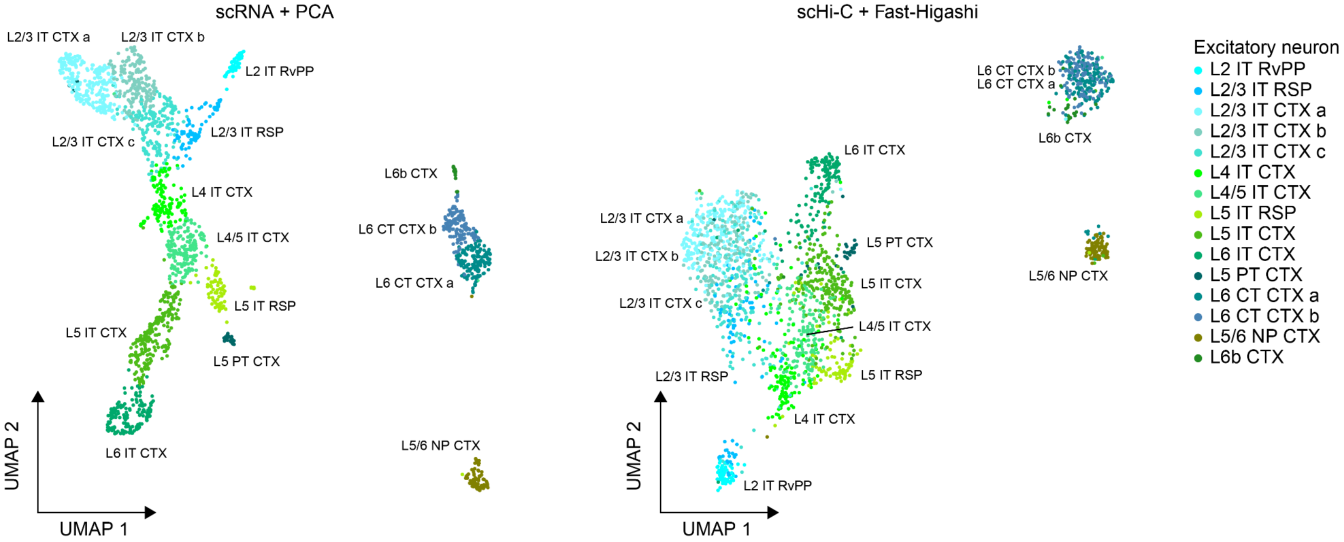


**Figure S13. High resolution excitatory neuron subtypes revealed by GAGE-seq.** Left panel, UMAP visualization of the PCA embeddings of GAGE-seq scRNA-seq profiles. Right panel, UMAP visualization of the Fast-Higashi embeddings of GAGE-seq sHi-C profiles. A strong correlation between the “structure” (scHi-C-based) and transcriptome (scRNA-based) cell types is observed.

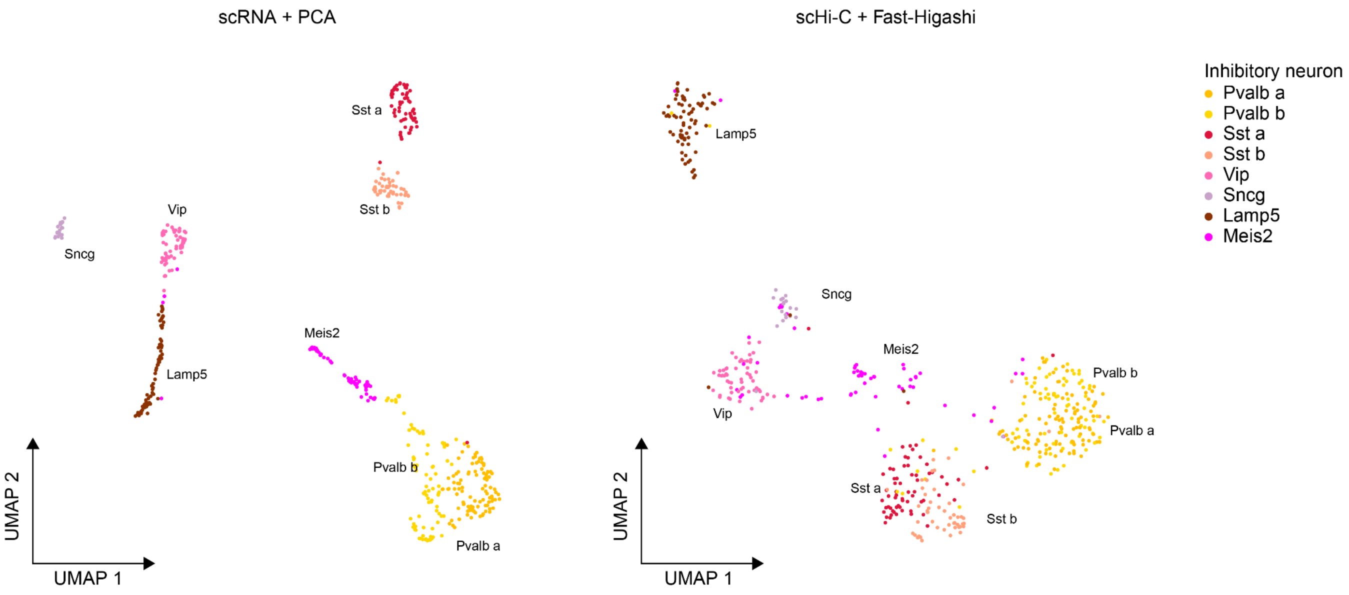


**Figure S14. High resolution inhibitory neuron subtypes revealed by GAGE-seq.** Left panel, UMAP visualization of the PCA embeddings of GAGE-seq scRNA-seq profiles. Right panel, UMAP visualization of the Fast-Higashi embeddings of GAGE-seq sHi-C profiles. A strong correlation between the “structure” (scHi-C-based) and transcriptome (scRNA-based) cell types is observed.

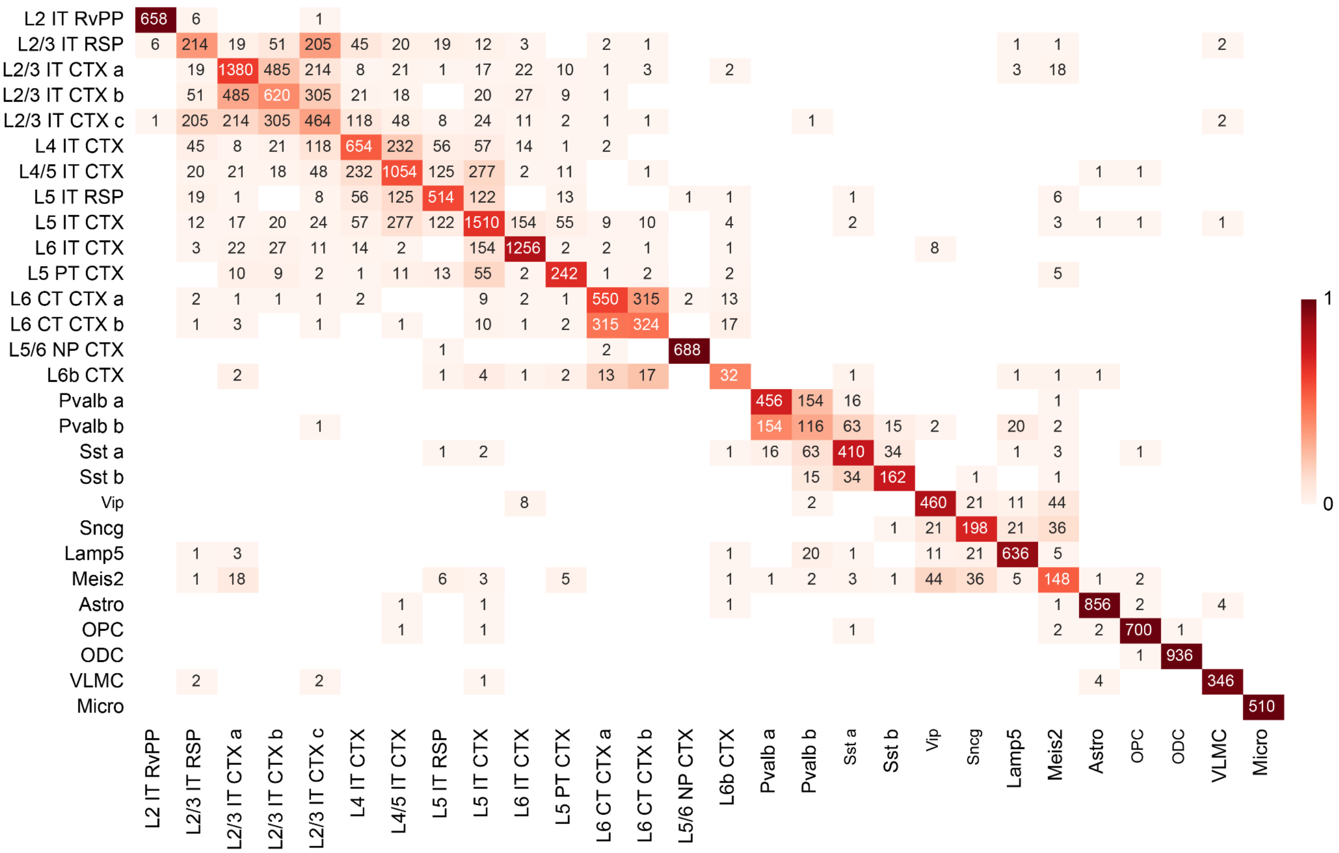


**Figure S15. Congruence between GAGE-seq scRNA-seq clusters and scHi-C embeddings.** The presented adjacency matrix shows the mutual 20-nearest neighbor graph of the Fast-Higashi embeddings, aggregated by cell types. Color intensity is normalized per row. When comparing the pairwise adjacency and self-adjacency, Fisher's exact test was applied to the 2-by-2 adjacency matrix for each pair of cell types. The maximum *P*-value of the one-sided Fisher's exact test was 2e-7, indicating all 28 cell types can be separated in the Fast-Higashi embeddings.

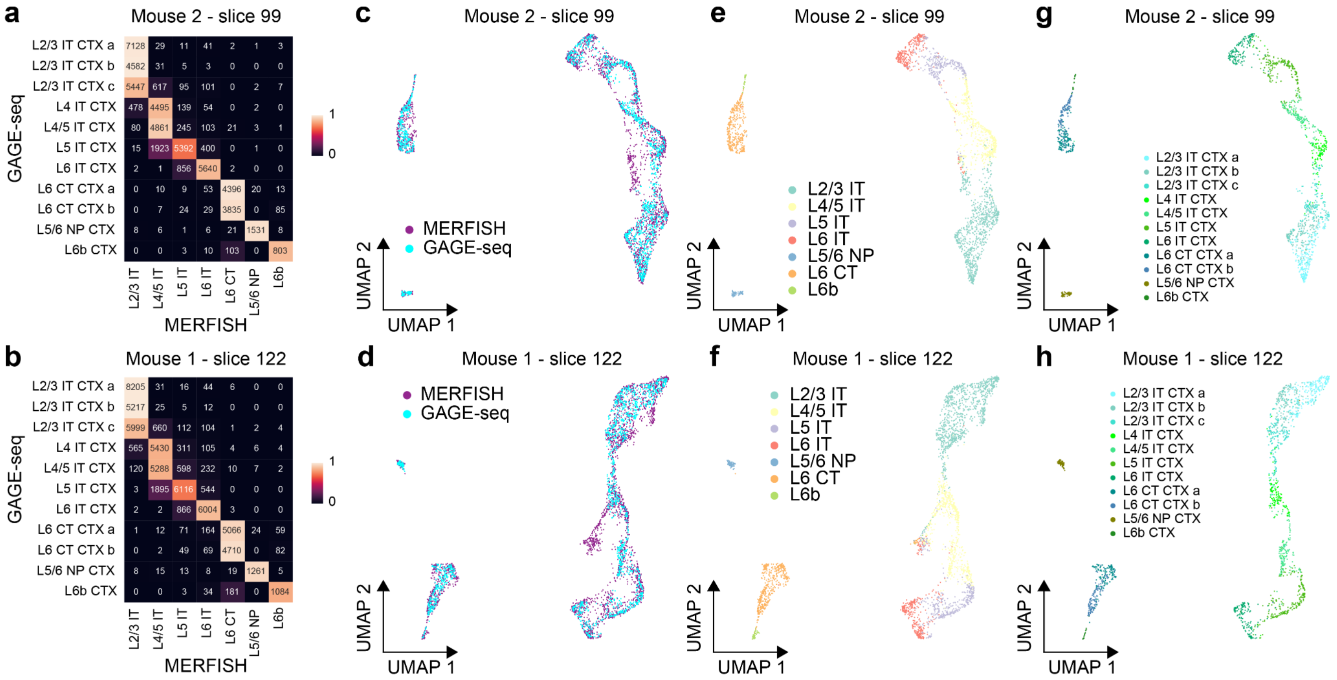


**Figure S16. Membership correspondence between GAGE-seq and MERFISH datasets.** Two tissue slices with the MERFISH dataset[19] are shown. **Top panels**. Slice 99 from mouse 2. **Bottom panels**. Slice 122 from mouse 1. **a-b**. Adjacency matrix of the nearest neighbor graph of the integrated embedding space, aggregated by cell type. For each cell from the MERFISH dataset, its 20 nearest neighbors from the GAGE-seq dataset were included. **c-h**. UMAP visualization of the integrated embedding space. **c-d** . cells from both datasets, colored by dataset. **e-f**. Cells from the MERFISH dataset, colored by the cell type annotation from the original analysis. **g-h**. Cells from the GAGE-seq dataset, colored by the cell type annotation.

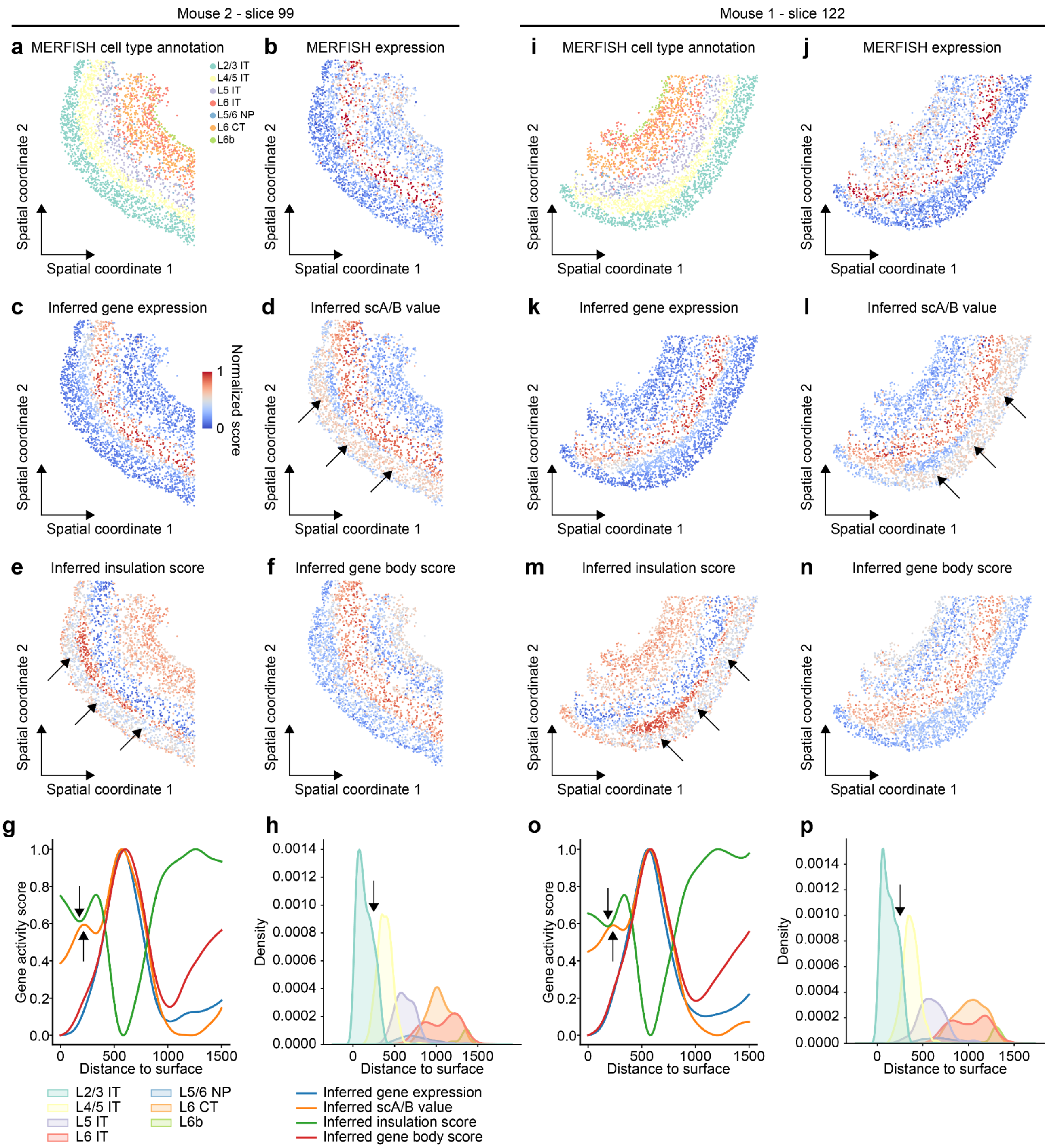


**Figure S17. High correlation between cortical layer-specific gene expression and the *in situ* dynamics of the 3D genome features of excitatory neurons.** *In situ* plots of two tissue slices with the MERFISH dataset[19] are shown. **a-h**. Slice 99 from mouse 2. **i-p**. Slice 122 from mouse 1. **a** and **i**, *in situ* plot of cell type annotations from the original analysis. In panels **b-g** and **j-o**, multiple activity scores of L5 IT CTX marker genes are shown. Activity scores were averaged across genes for each cell. **b** and **j**. Detected expression level. **c** and **k**. Inferred expression level. **d** and **l**. Inferred scA/B value. **e** and **m**. Inferred single-cell insulation score. **f** and **n**. Inferred gene body score. **g** and **n**. The spatial gradient of different features with respect to the distance to the surface. **h** and **p**. The distribution of cell types with respect to the distance to surface.

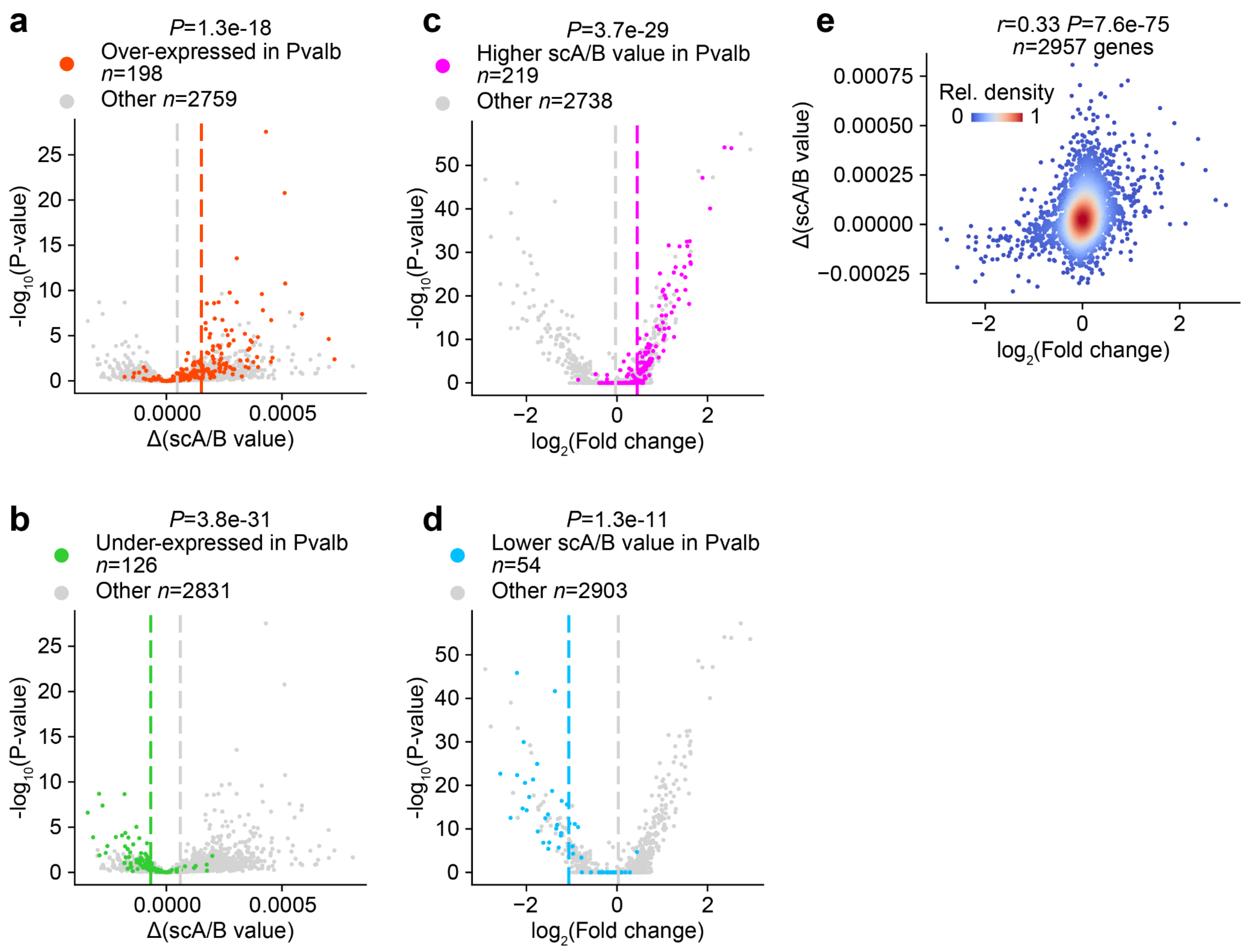


**Figure S18. Correlation between cell type-specific single-cell A/B value and gene expression when comparing Pvalb and the other inhibitory neurons. a** and **b**. Volcano plot of differential scA/B value. **c** and **d**. Volcano plot of differential gene expression. **e**. The whole-transcriptome correlation between differential expression and differential scA/B value. The CpG-based scA/B value was used. *P*-value of one-sided t-test is shown in panels a-d. Pearson's correlation coefficient and one-sided *P*-value are shown in panel e.

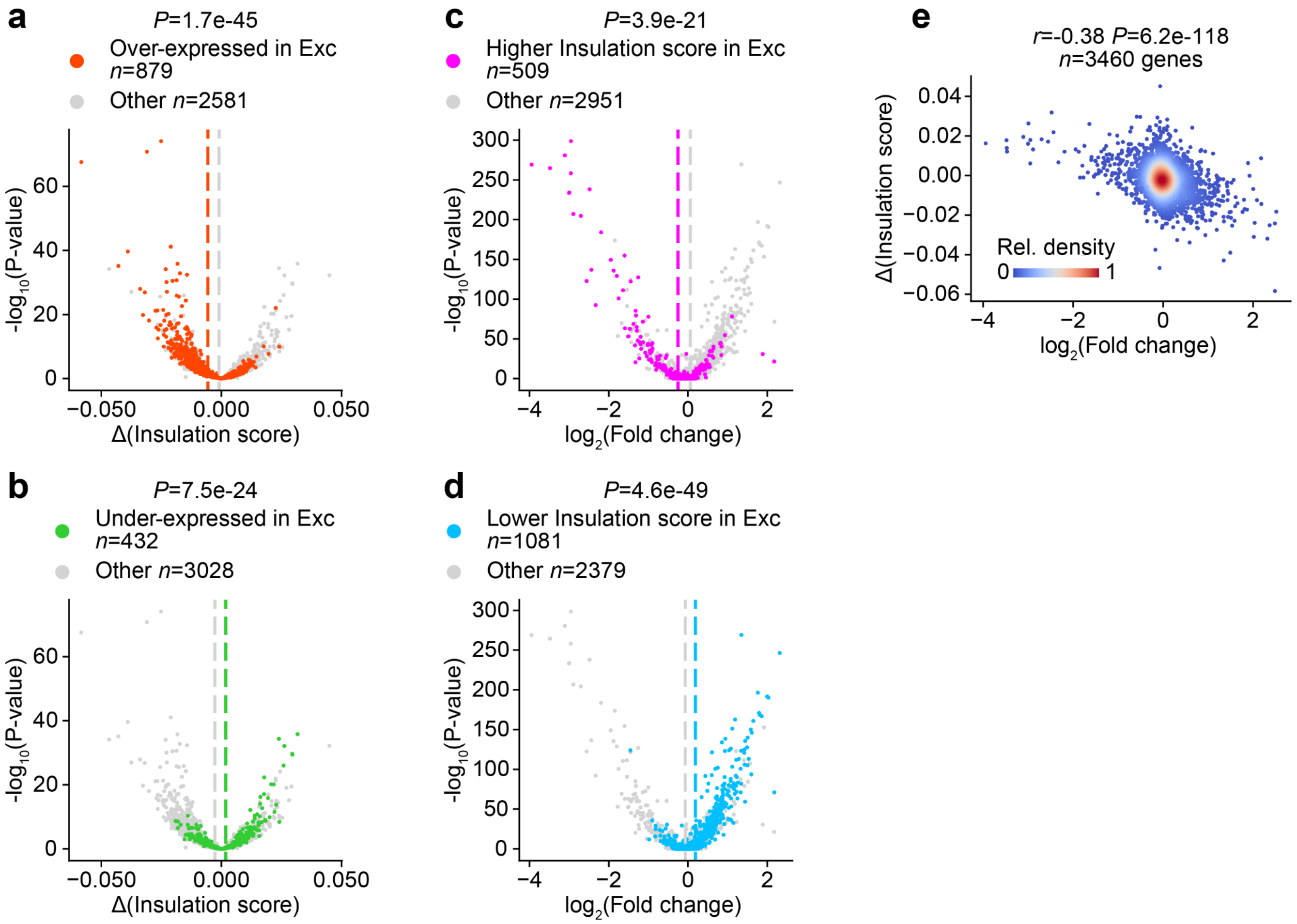


**Figure S19. Correlation between cell-type-specific single-cell insulation score and gene expression when comparing excitatory and inhibitory neurons. a** and **b**. Volcano plot of differential single-cell insulation score. **c** and **d**. Volcano plot of differential gene expression. **e**. The whole-transcriptome correlation between differential expression and differential single-cell insulation score. *P*-value of one-sided t-test is shown in panels a-d. Pearson's correlation coefficient and one-sided *P*-value are shown in panel e.

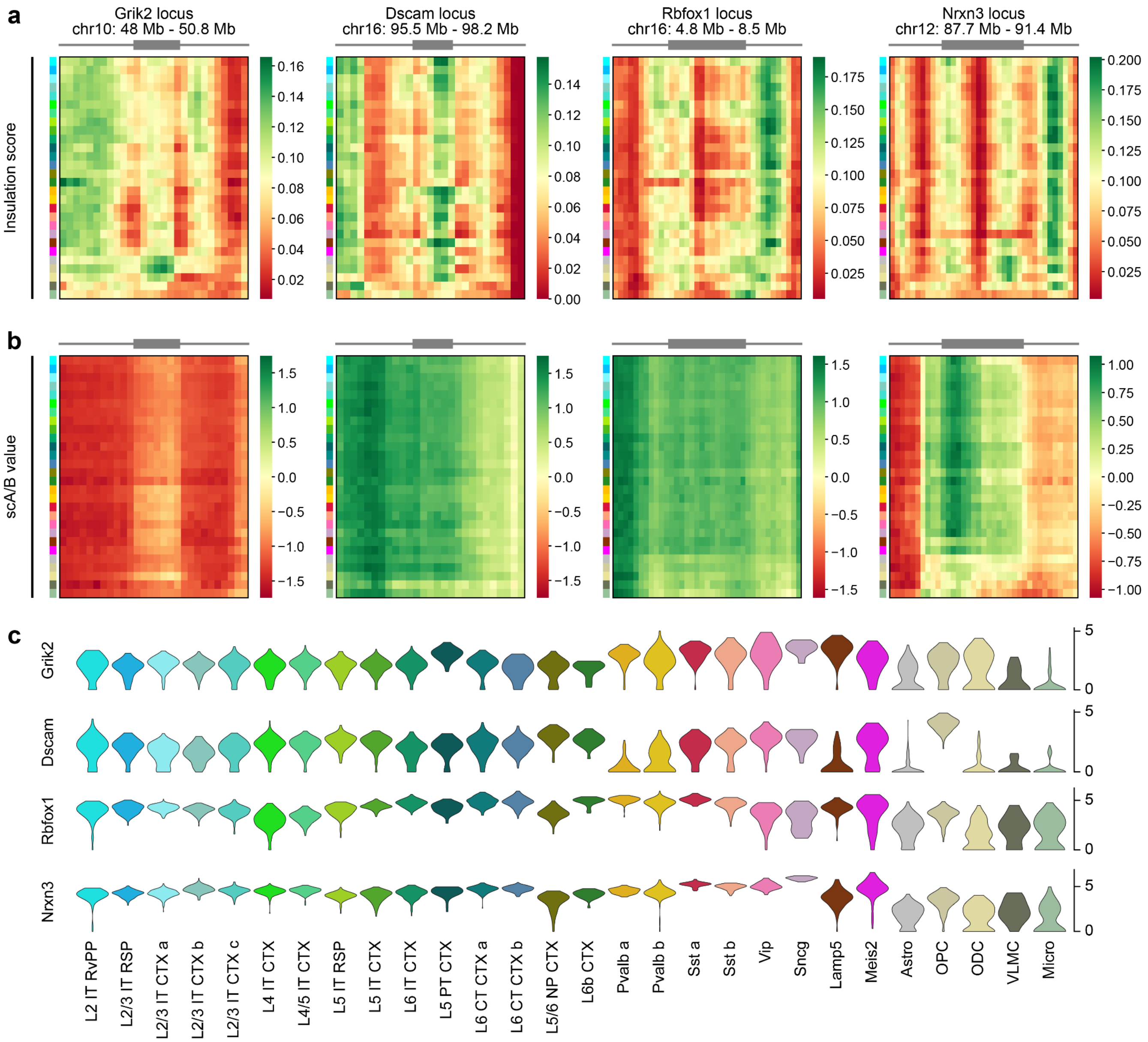


**Figure S20. Aggregated single-cell insulation score and scA/B value of the four gene loci, *Grik2*, *Dscam*, *Rbfox1* and *Nrxn3* in the annotated 28 cell subtypes. a**. Aggregated single-cell insulation score calculated on raw contact maps. **b**. Z-scored eigenvector-based scA/B value calculated on Higashi-imputed contact maps. **c**. Violin plots showing single-cell gene expression profiles of the four genes. Gene bodies are shown as gray boxes above heat maps.

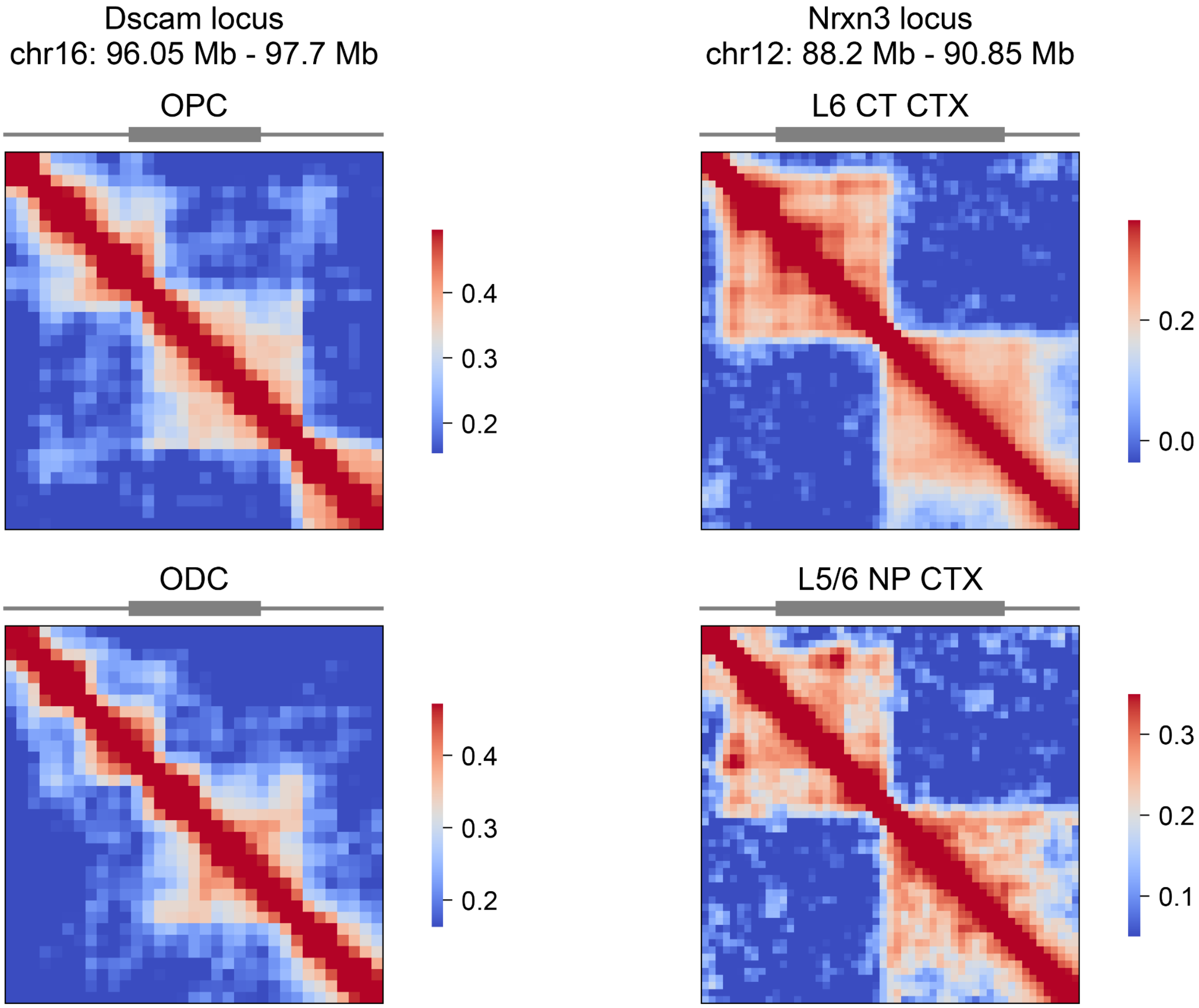


**Figure S21. Aggregated contact maps of the *Dscam* and *Nrxn3* gene loci showing cell type-specific domain organization**. For each locus, two cell types with differential expression were selected and the two aggregated contact maps are shown. All contact maps are at the 50 Kb resolution and are normalized by NPMI. Gene bodies are shown as gray boxes above heat maps.

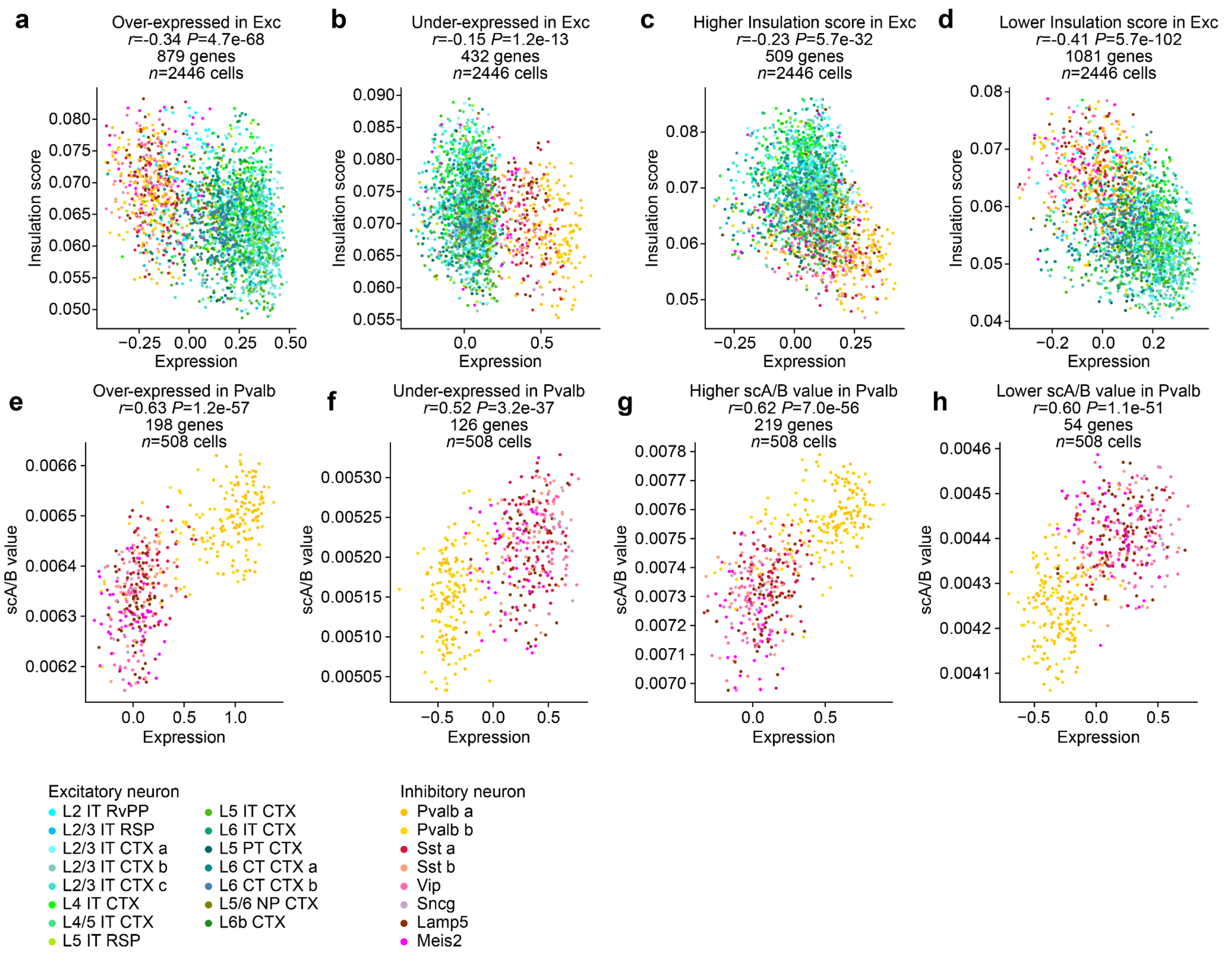


**Figure S22. Correlation between gene expression and 3D genome features at the single-cell level**. For each cell, the average expression and 3D genome features of a particular set of genes are shown. **a-d**. The correlation between expression and single-cell insulation score. **e-h**. The correlation between expression and CpG-based scA/B value. Each cell type was denoted by a distinct color shown at the bottom. The gene set in each panel: **a**. Genes over-expressed in excitatory neurons; **b**. Genes under-expressed in excitatory neurons; **c**. Genes having higher sc-insulation scores in excitatory neurons; **d**. Genes having lower sc-insulation scores in excitatory neurons; **e**. Genes over-expressed in Pvalb; **f**. Genes under-expressed in Pvalb; **g**. Genes having higher scA/B values in Pvalb; **h**. Genes having lower scA/B in Pvalb. Pearson's correlation coefficients and the *P*-values for one-sided test are shown.

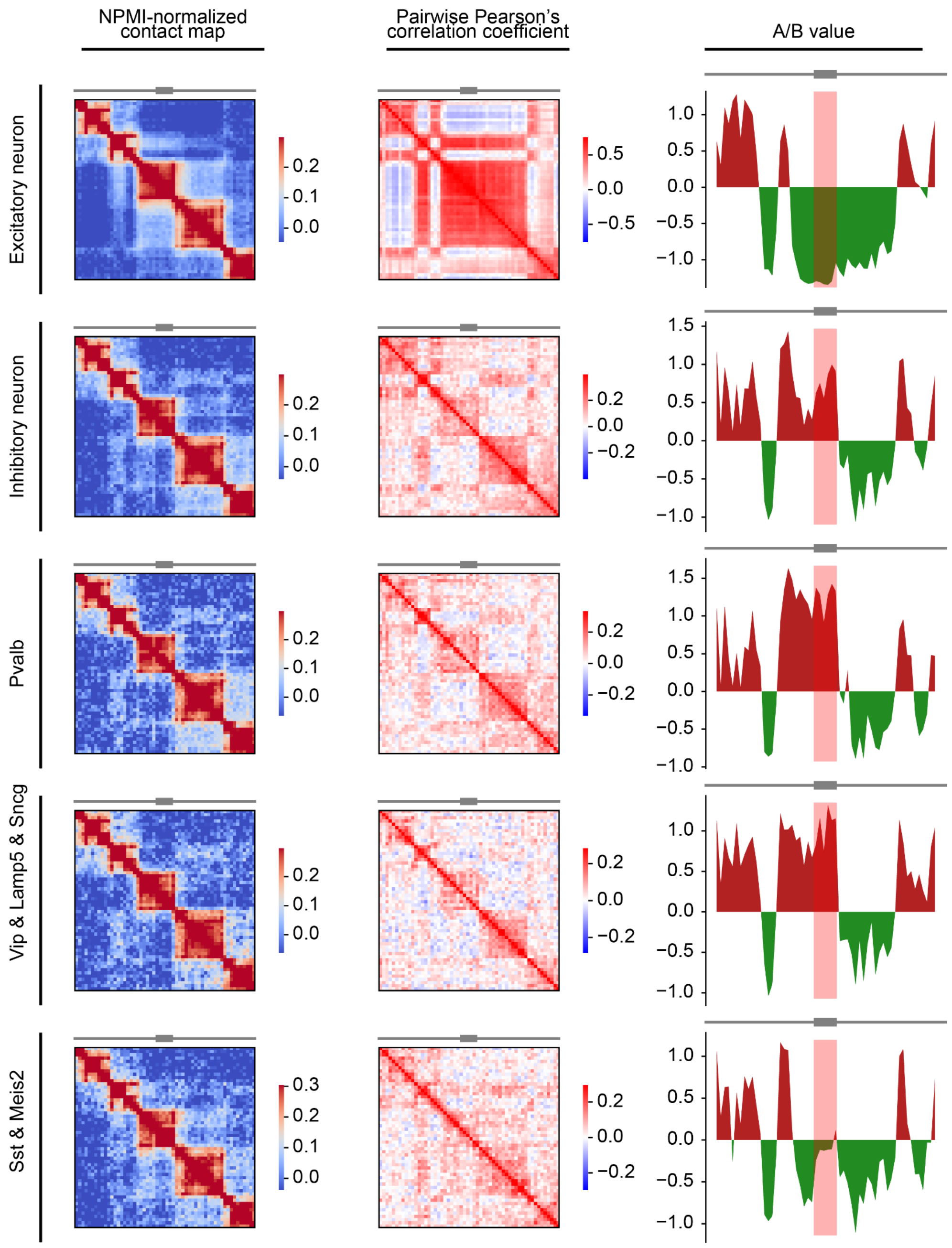


**Figure S23. A/B values and contact maps of the Erbb4 locus.** The local 3D genome features of the Erbb4 locus (chr1: 63 Mb - 74.2 Mb) are shown for 5 groups of cells. The resolution of all panels is 200 Kb. **Left column**, NPMI-normalized contact map. **Middle column**, pairwise Pearson's correlation coefficient. **Right column**, z-scored eigenvalue-based A/B value. The gene body (highlighted by the pink rectangle) is in the active A compartment in all of the last 4 groups of cells, because the A/B value of the gene body either is strongly positive or attains a local maximum.

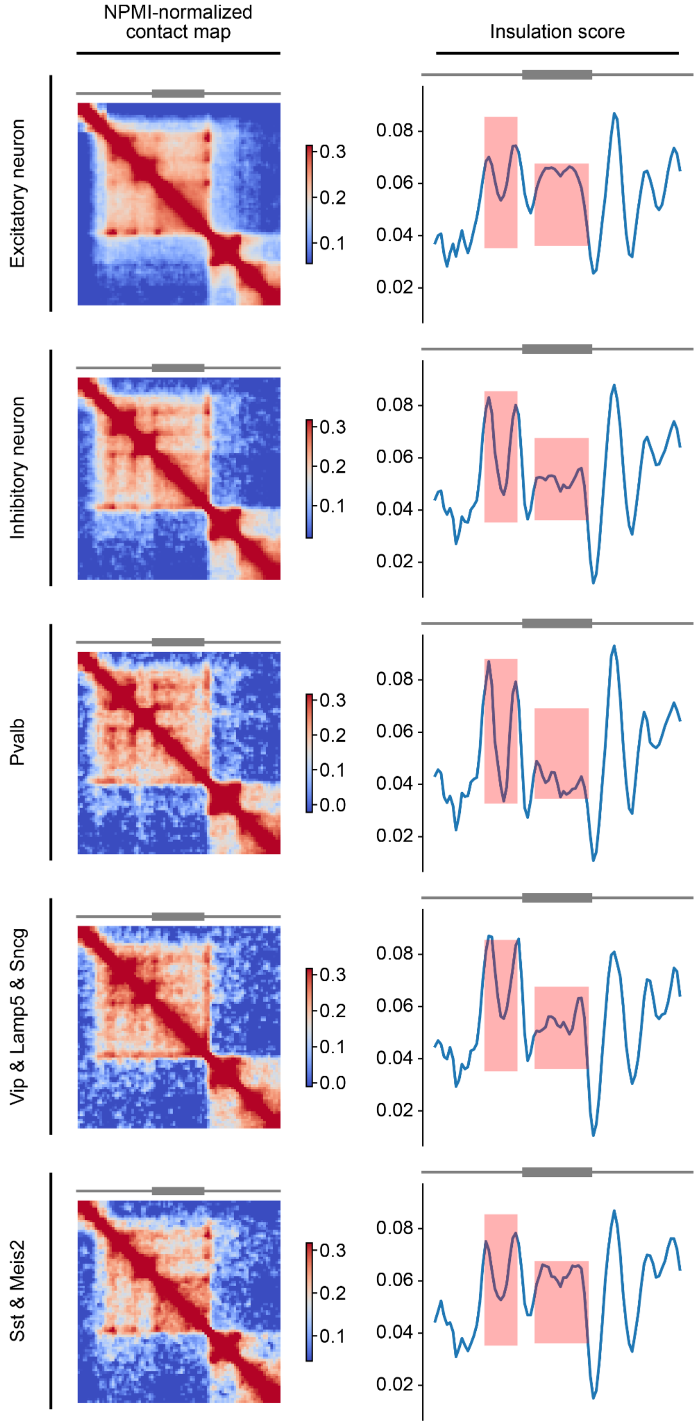


**Figure S24. Insulation scores and contact maps of the Erbb4 locus.** The local 3D genome features of the Erbb4 locus (chr1: 66.5 Mb - 70.65 Mb) are shown for 5 groups of cells. The resolution of all panels is 50 Kb. **Left column**, NPMI-normalized contact map. **Right column**, insulation score. Among inhibitory subtypes (the last 3 rows), the insulation score at the gene body and a downstream region (both highlighted by pink rectangles) are lowest in Pvalb (the 3rd row) and highest in the Sst and Meis2 (the last row).

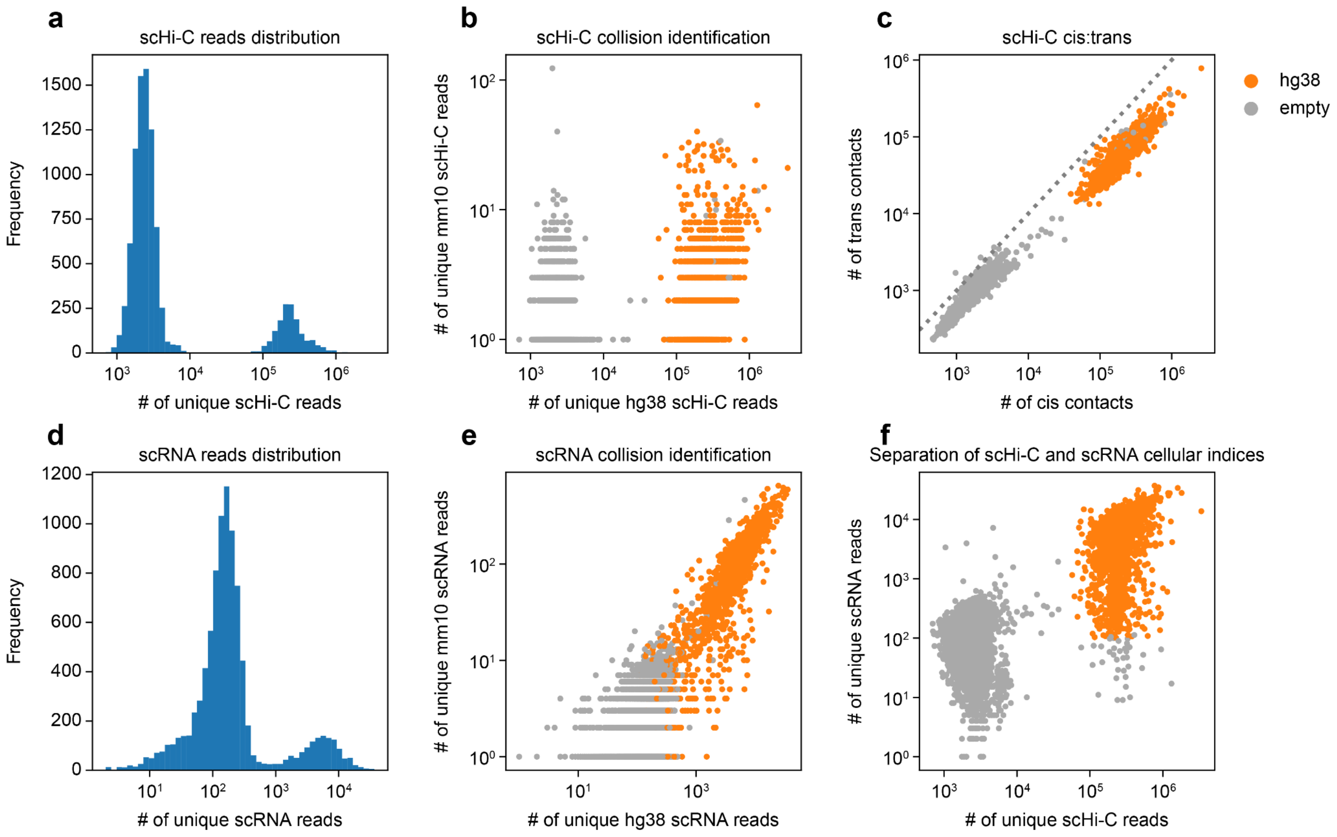


**Figure S25. Quality-control of the GAGE-seq human bone marrow CD34+ cells library (replicate 1). a** and **d.** Histogram showing the binomial distribution of GAGE-seq scHi-C **(**a**)** and scRNA-seq reads **(**d**)**. **b** and **e**. Scatter plots showing the collision level in the GAGE-seq scHi-C **(**b**)** and scRNA-seq **(**e**)** libraries. **c**. Scatter plot showing the cis:trans ratio of scHi-C reads. **f.** Scatter plot showing the well-separation of DNA and RNA reads of valid cellular indices from that of empty indices. Human data are colored in orange and empty indices in gray.

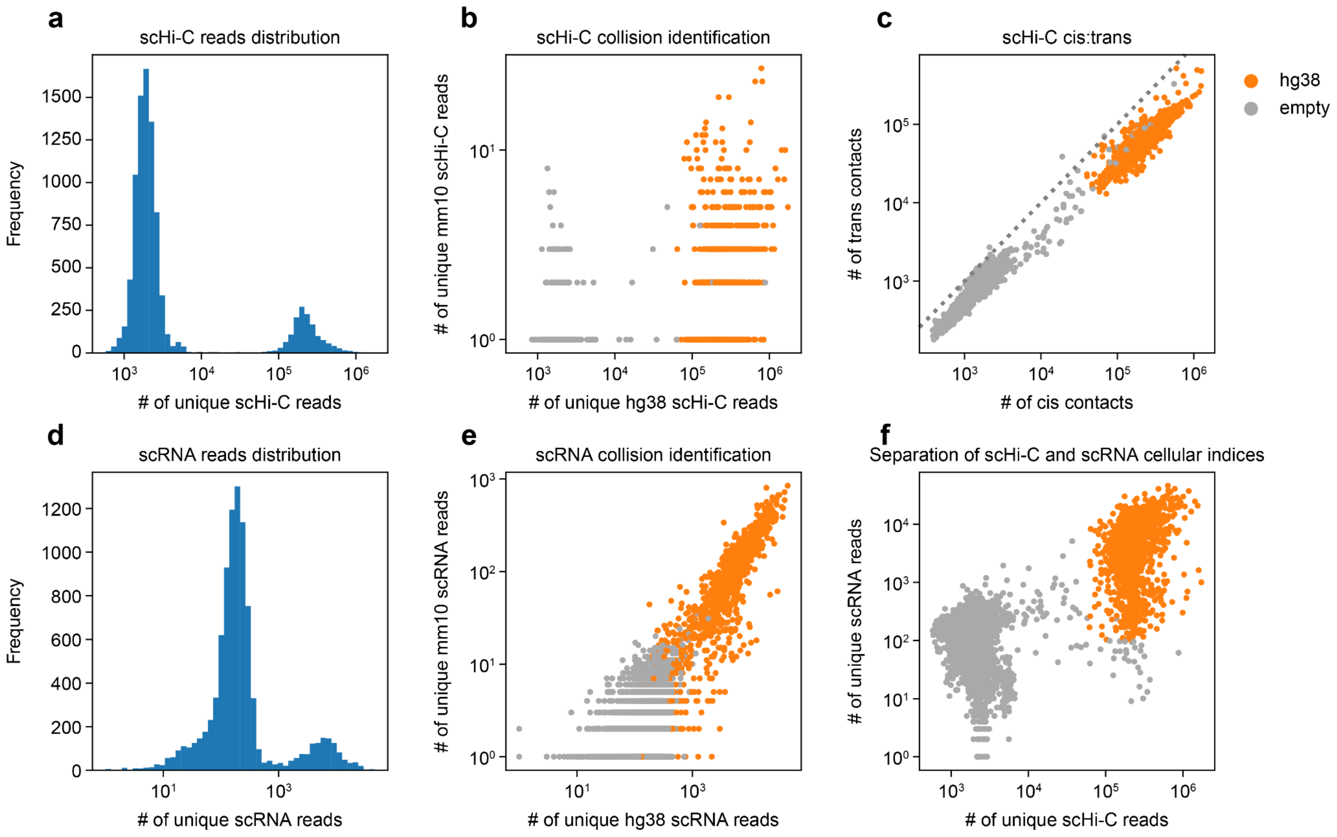


**Figure S26. Quality-control assessment of the GAGE-seq human bone marrow CD34+ cells library (replicate 2). a** and **d**. Histogram showing the binomial distribution of GAGE-seq scHi-C (a) and scRNA-seq reads (d). **b** and **e**. Scatter plots showing the collision level in the GAGE-seq scHi-C (b) and scRNA-seq (e) libraries. **c**. Scatter plot showing the cis:trans ratio of scHi-C reads. **f**. Scatter plot showing the well-separation of DNA and RNA reads of valid cellular indices from that of empty indices. Human data are colored in orange and empty indices in gray.

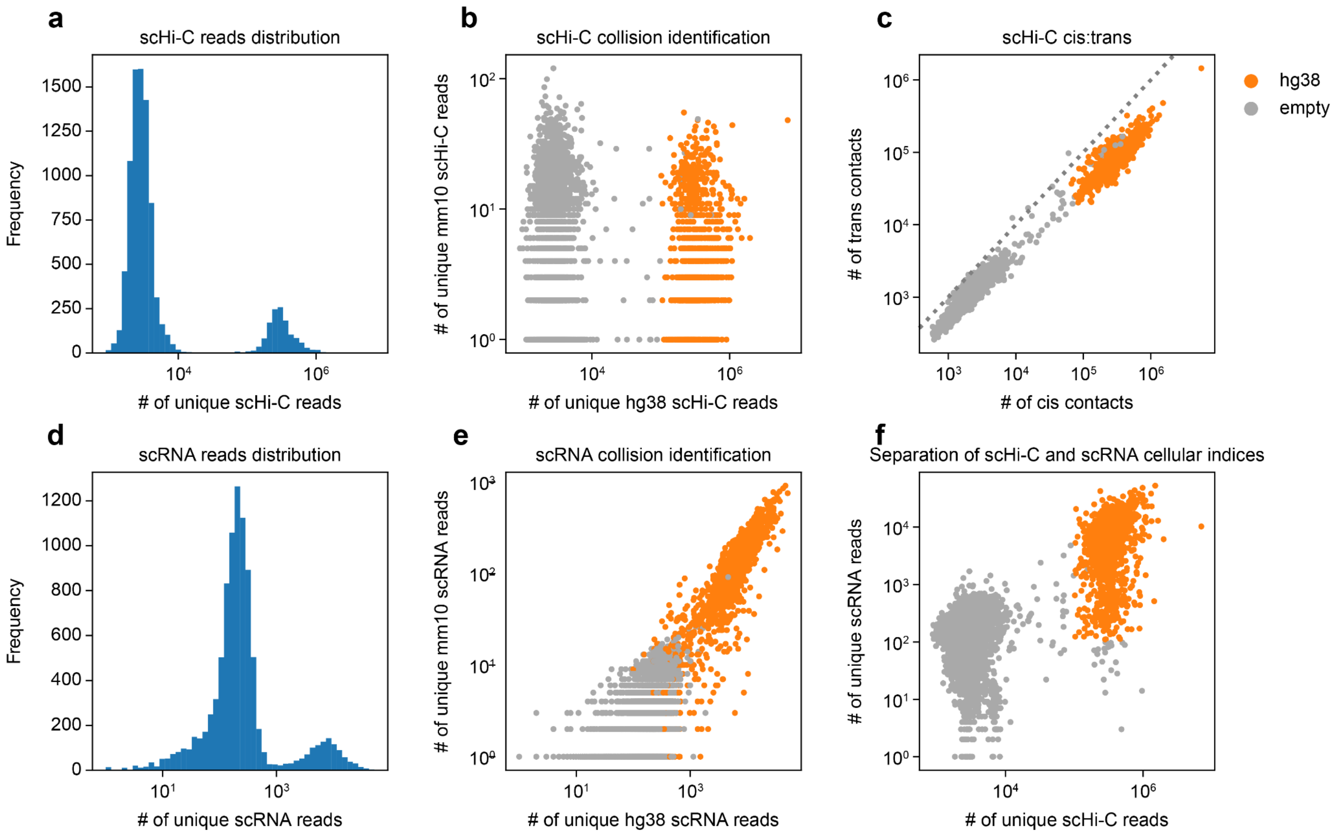


**Figure S27. Quality-control assessment of the GAGE-seq human bone marrow CD34+ cells library (replicate 3). a** and **d**. Histogram showing the binomial distribution of GAGE-seq scHi-C **(**a**)** and scRNA-seq reads **(**d**)**. **b** and **e**. Scatter plots showing the collision level in the GAGE-seq scHi-C **(**b**)** and scRNA-seq **(**e**)** libraries. **c**. Scatter plot showing the cis:trans ratio of scHi-C reads. **f.** Scatter plot showing the well-separation of DNA and RNA reads of valid cellular indices from that of empty indices. Human data are colored in orange and empty indices in gray.

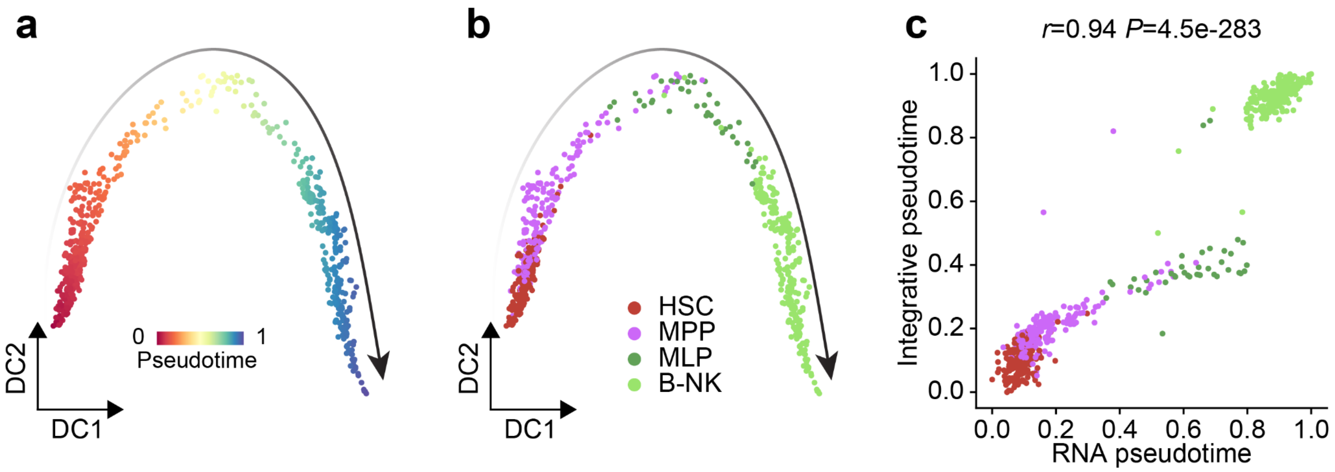


**Figure S28. RNA-based B-NK trajectory of the GAGE-seq human bone marrow datasets. a** and **b.** The diffusion map visualization of the scRNA profiles in the B-NK trajectory, colored by pseudotime (a) and cell type annotation (b). **c**. The high correlation between the RNA-based pseudotime and the integrative pseudotime shown in **Fig 5c**. Spearman's correlation coefficient and the two-sided *P*-value are shown.

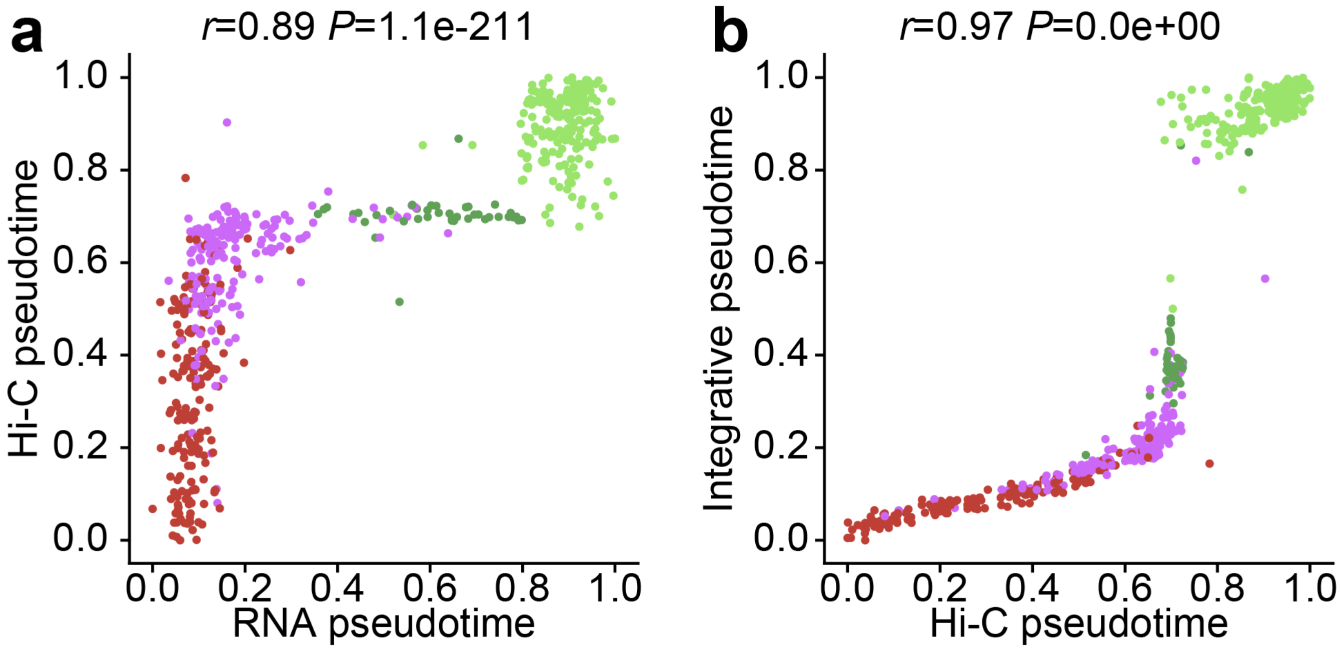


**Figure S29. Hi-C-based B-NK trajectory of the GAGE-seq human bone marrow datasets. a** and **b**. The diffusion map visualization of the scHi-C profiles in the B-NK trajectory, colored by pseudotime (a) and cell type annotation (b). **c** and **d**. The high correlation between the Hi-C-based pseudotime and the RNA-based pseudotime (c) or the integrative pseudotime (d). Spearman's correlation coefficient and the two-sided *P*-value are shown.

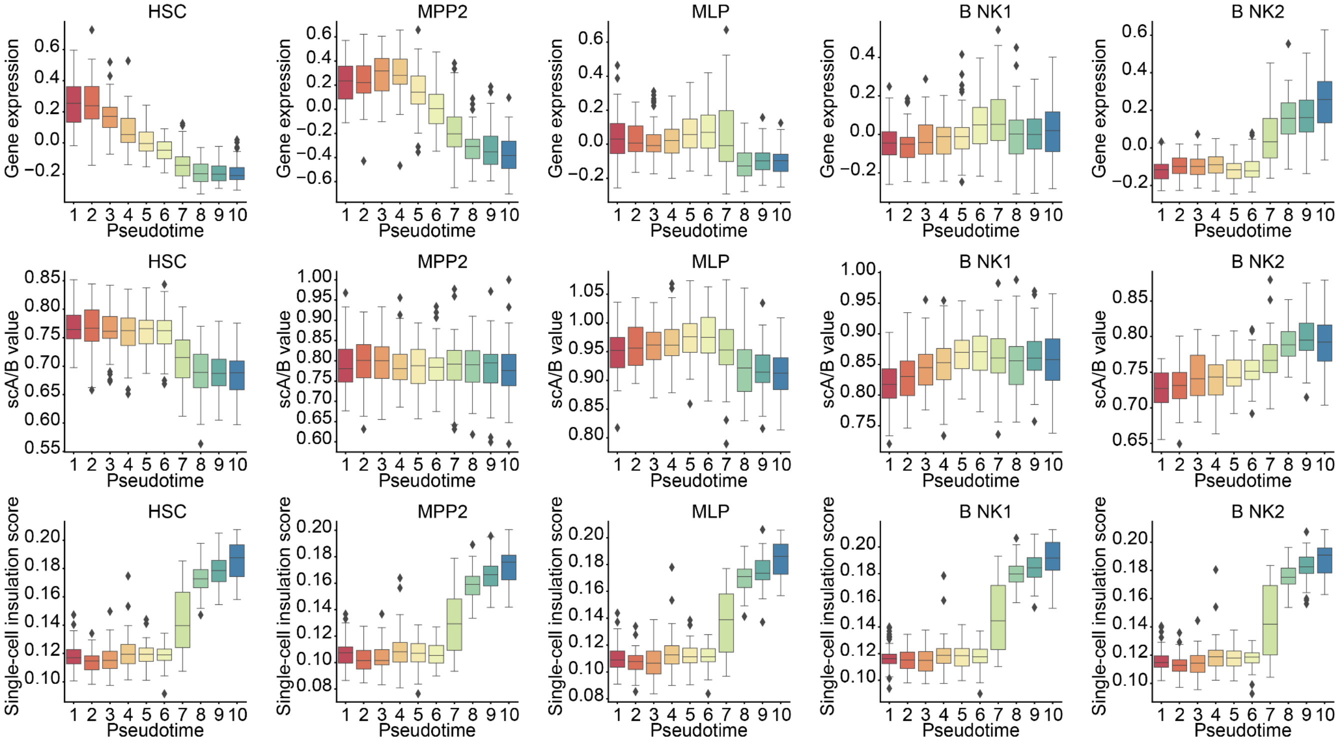


**Figure S30. Temporal pattern of gene expression, scA/B value, and sc-insulation score of the marker gene sets along the B-NK trajectory.** The marker gene sets of five developmental stages along the B-NK trajectory were selected from[20], with one column for each gene set. Gene expression (**top row**), scA/B value (**middle row**), and single-cell insulation score (**bottom row**) are shown. The resolution of all scores is 100 Kb. Cells are grouped according to inferred pseudotime.

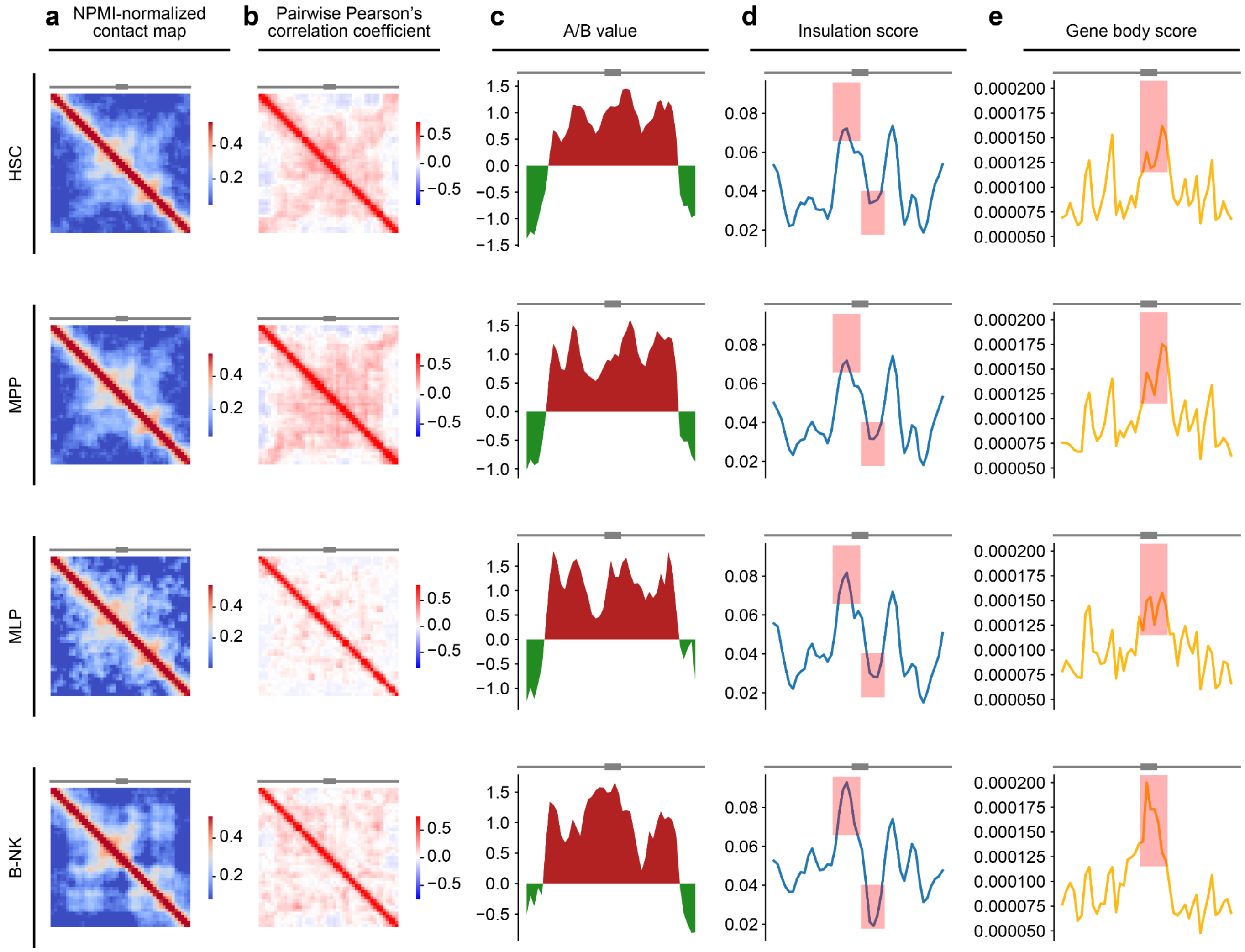


**Figure S31. Aggregated contact maps, A/B compartment, insulation score, and gene body score of the EBF1 locus.** The local 3D genome features of the EBF1 locus (chr5: 156.6 Mb-161.1 Mb) are shown for four differentiation stages, and each row represents a cell stage. The resolution of all panels is 100 Kb. **Column a**, NPMI-normalized contact map. **Column b**, pairwise Pearson's correlation coefficient. **Column c**, z-scored eigenvalue-based A/B value. **Column d**, insulation score. **Column e**, gene body score. Differences between B-NK and other cell stages are highlighted by pink rectangles.

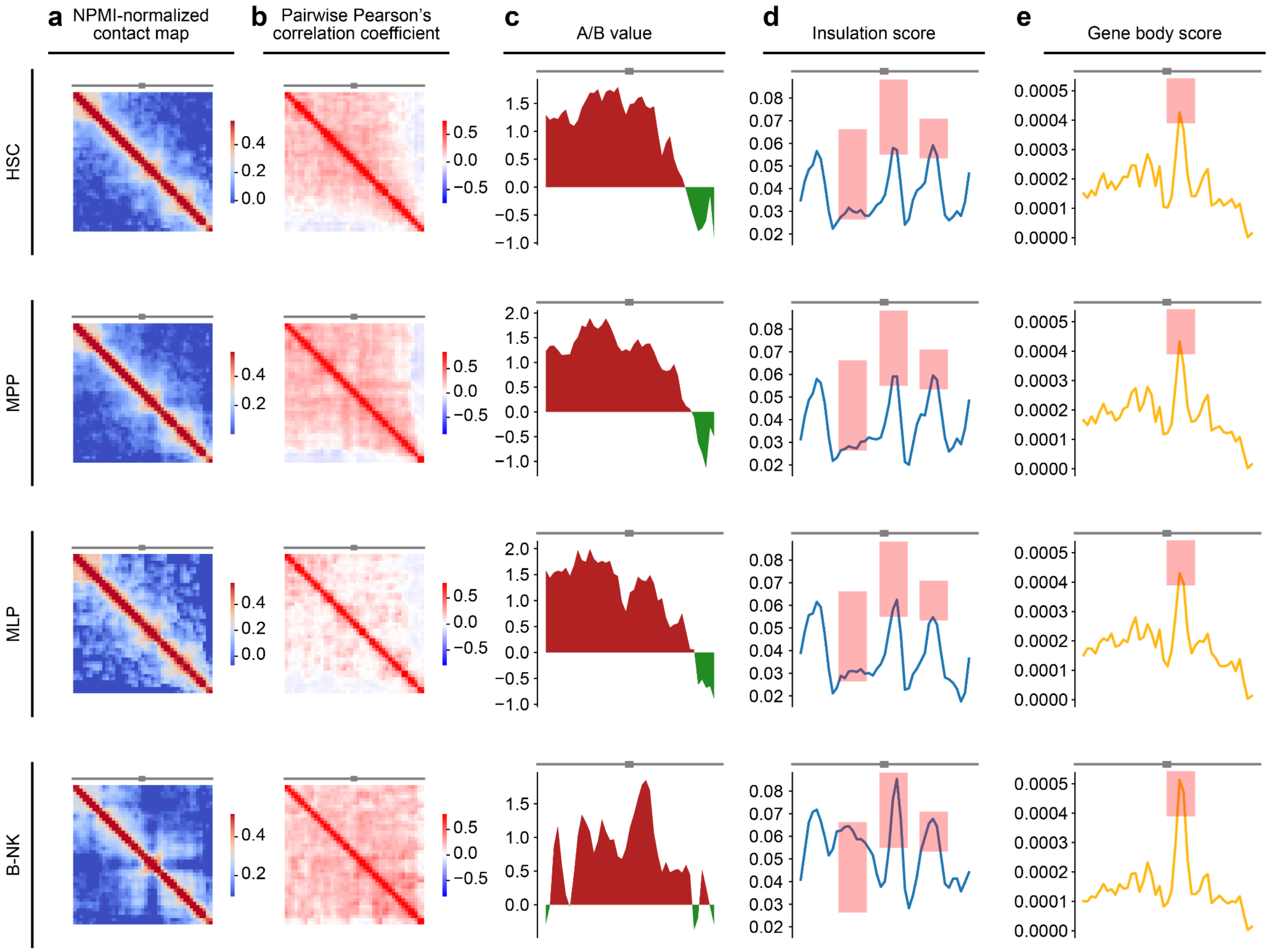


**Figure S32. Aggregated contact maps, A/B, insulation score, and gene body score of the PAX5 locus.** The local 3D genome features of the EBF1 locus (chr 9: 34.8 Mb-39.1 Mb) are shown for four differentiation stages, and each row represents a cell stage. The resolution of all panels is 100 Kb. **Column a**, NPMI-normalized contact map. **Column b**, pairwise Pearson's correlation coefficient. **Column c**, z-scored eigenvalue-based A/B value. **Column d**, insulation score. **Column e**, gene body score. Differences between B-NK and other cell stages are highlighted by pink rectangles.

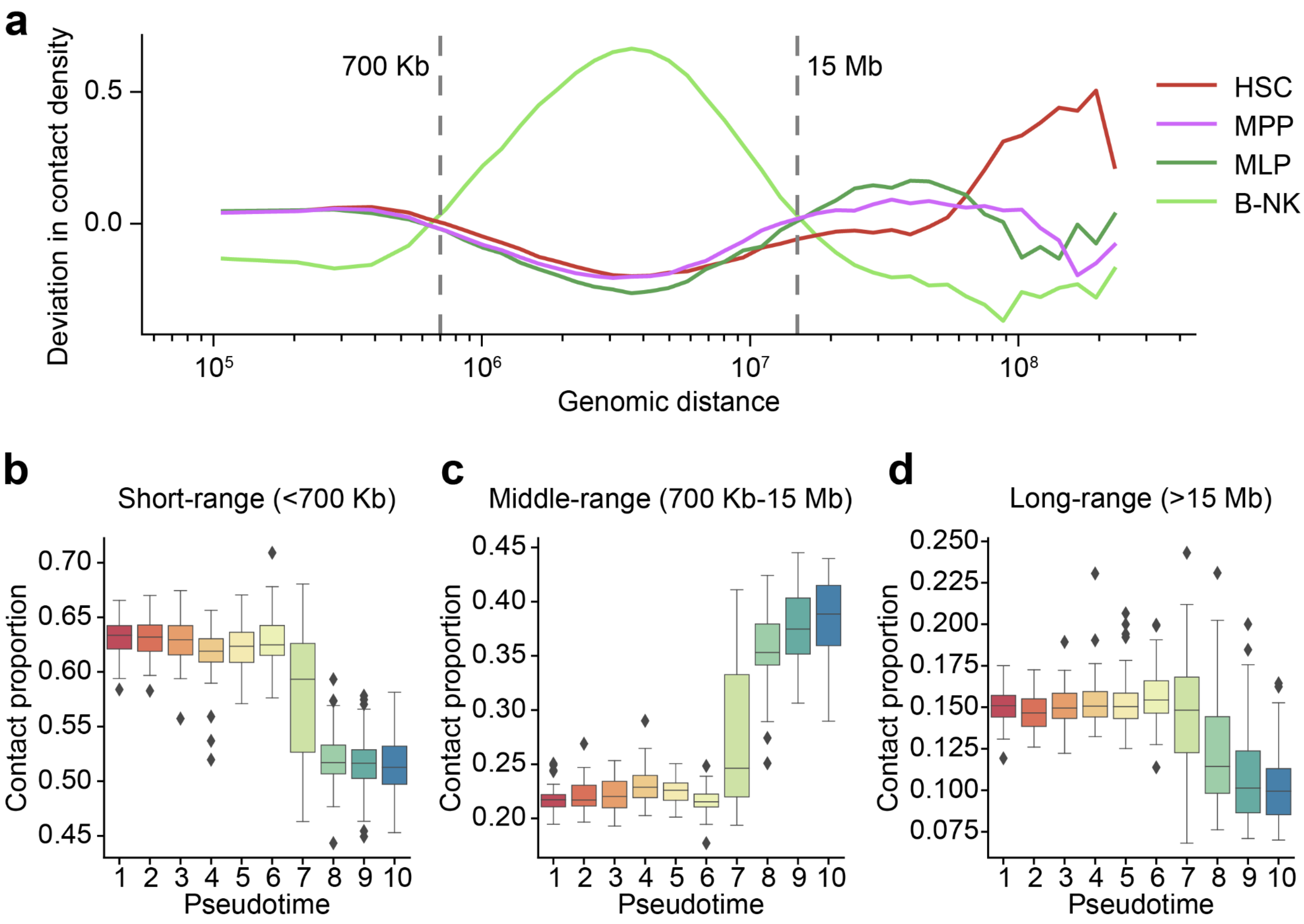


**Figure S33. Global changes in chromatin contacts in B-NK progenitor cells. a**. The differential contact frequency curves for the four cell stages. Pseudo-bulk contact maps were calculated and normalized by coverage for the 4 cell stages. The contact frequency at each genomic distance was calculated. The average frequency curve was defined as the average of the 4 frequency curves. The average frequency curve was subtracted from the frequency curve for each cell type and the difference was divided also by the average curve, which yielded the differential contact frequency curves shown in the figure. **b-d**. The proportion of contacts in the 3 distance groups. The 3 groups are divided based on genomic distance and the 2 thresholds are shown in panel (a).

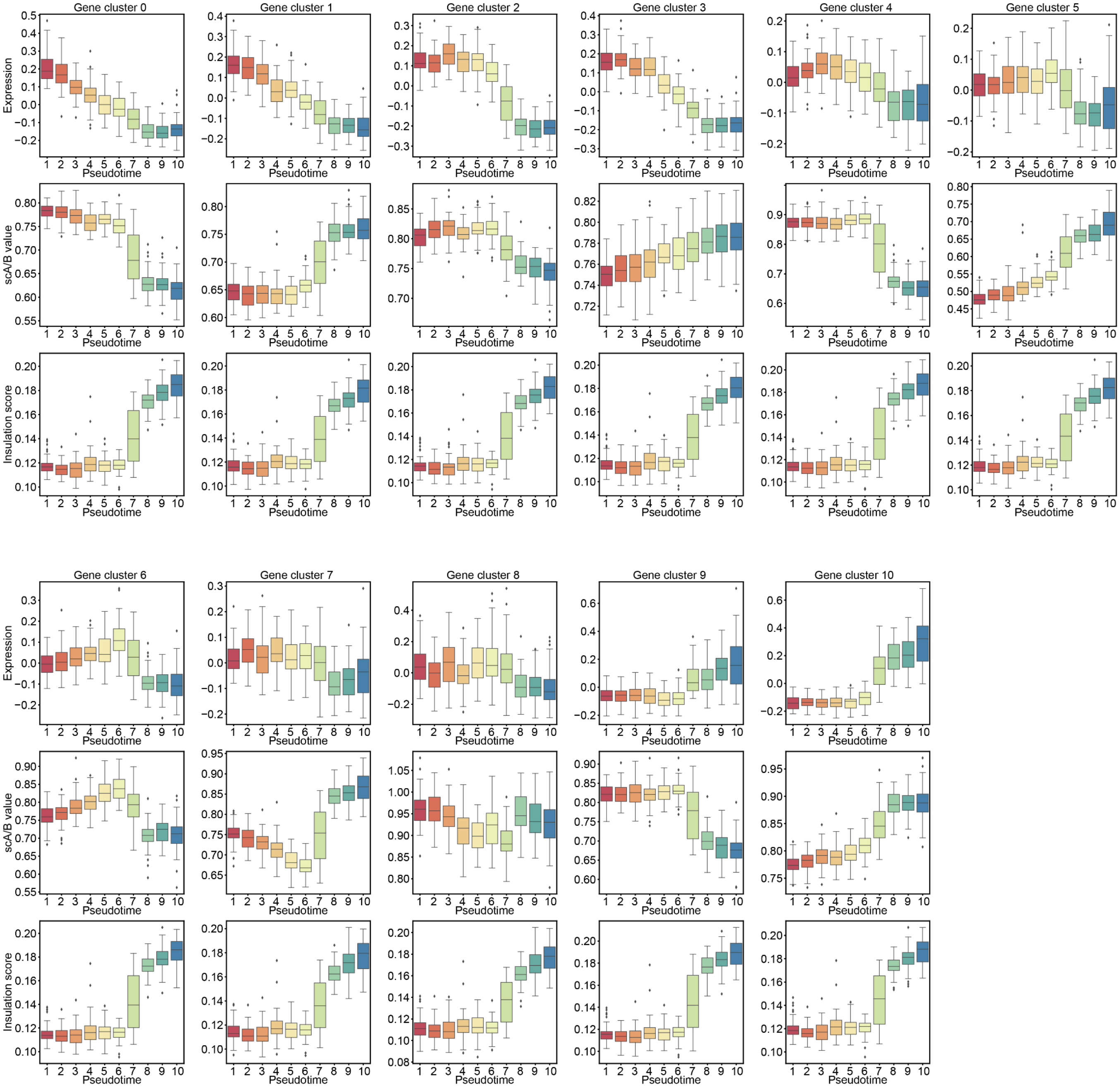


**Figure S34. The temporal trends of multi-modal features of gene clusters along the B-NK trajectory.** The 11 gene clusters (**Fig. 5e**) from the unsupervised clustering procedure are shown, with 6 in the upper half of the figure and 5 in the bottom half. For each gene cluster, expression (**top panel**), scA/B value (**middle panel**), and single-cell insulation score (**bottom panel**) are shown.

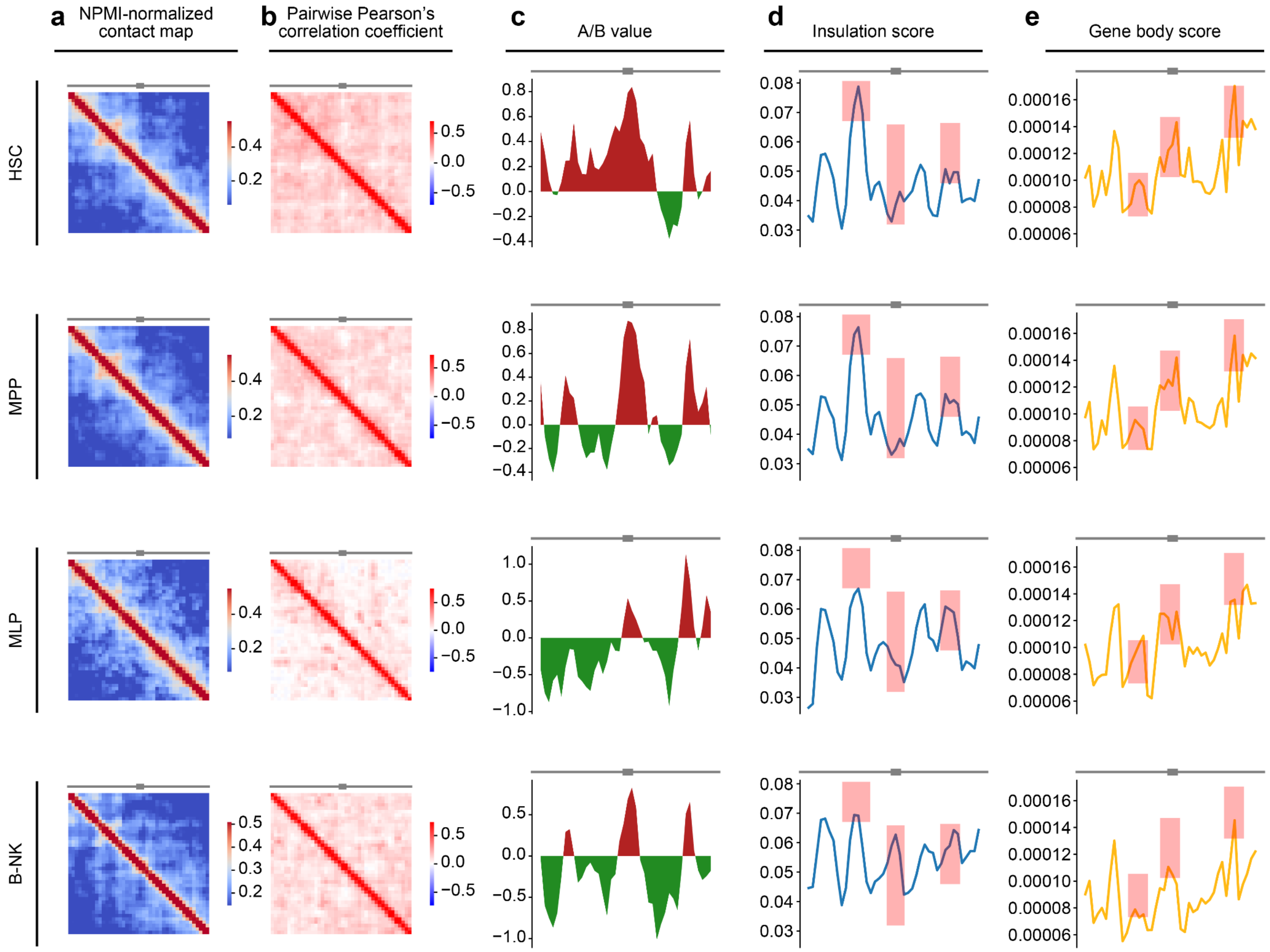


**Figure S35. Aggregated contact maps, A/B, insulation score, and gene body score of the JAK1 locus**. The local 3D genome features of the JAK1 locus (chr1: 62.8 Mb-67 Mb) are shown for four differentiation stages, and each row represents a cell stage. The resolution of all panels is 100 Kb. **Column a**, NPMI-normalized contact map. **Column b**, pairwise Pearson's correlation coefficient. **Column c**, z-scored eigenvalue-based A/B value. **Column d**, insulation score. **Column e**, gene body score. Differences between B-NK and other cell stages are highlighted by pink rectangles.

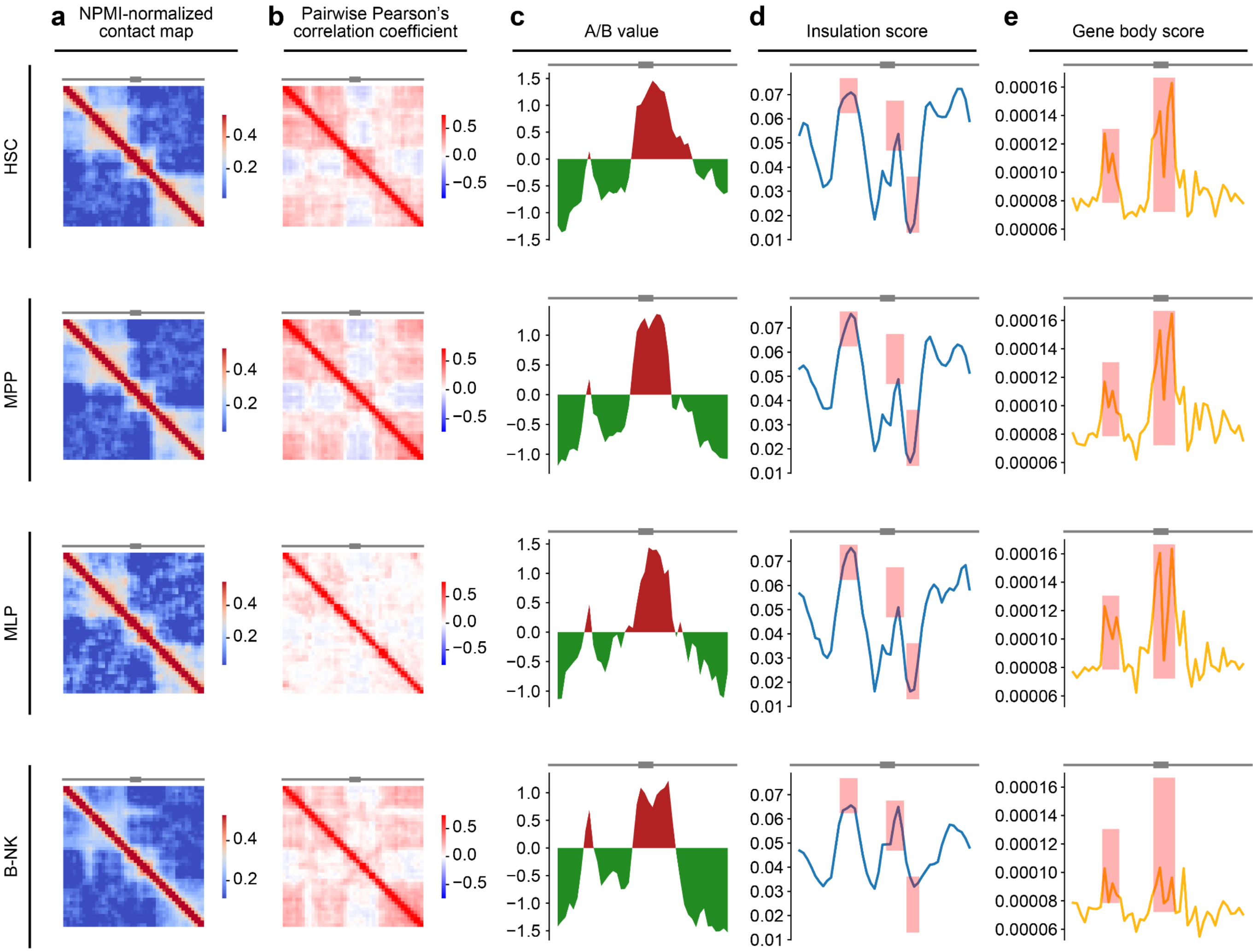


**Figure S36. Aggregated contact maps, A/B, insulation score, and gene body score of the ITPR1 locus**. The local 3D genome features of the ITPR1 locus (chr 3: 2.4 Mb-6.8 Mb) are shown for four differentiation stages, and each row represents a cell stage. The resolution of all panels is 100 Kb. **Column a**, NPMI-normalized contact map. **Column b**, pairwise Pearson's correlation coefficient. **Column c**, z-scored eigenvalue-based A/B value. **Column d**, insulation score. **Column e**, gene body score. Differences between B-NK and other cell stages are highlighted by red rectangles.

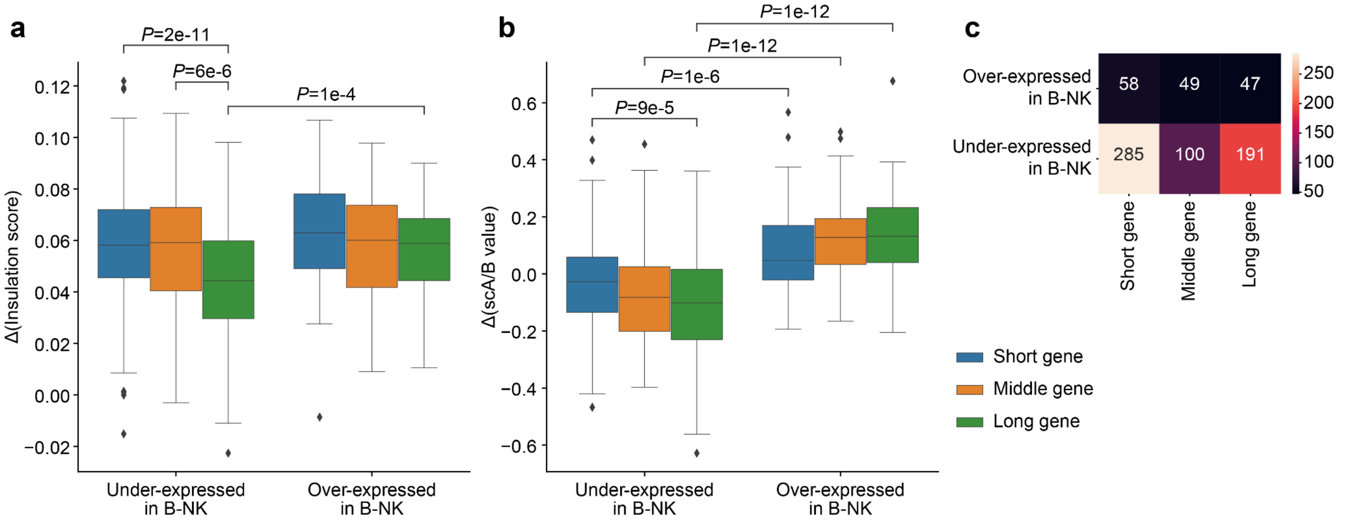


**Figure S37. 3D genome reorganization at the gene loci of the B-NK cell differentially expressed (DE) genes with different gene lengths. a** and **b.** Differential single-cell insulation score (a) and scA/B value (b) of B-NK DEGs. Genes are grouped by both gene length and differential expression. Short genes (length < 100 Kb) are colored in blue. Middle genes (length in 100 Kb - 200 Kb) are colored in orange. Long genes (length > 200 Kb) are colored in green. **c**. The number of genes in each gene group.
